# High-Dimensional Spatiotemporal Single-Cell and 3D Atlas of the Bone Marrow Microenvironment during Leukemic Progression

**DOI:** 10.1101/2025.09.27.678113

**Authors:** Lanzhu Li, Isabelle Rottmann, Geoff Ivison, Huan Wei, Jan C. Schroeder, Gina Dunkel, Yury Goltsev, Garry P. Nolan, Aaron T. Mayer, Borhan R. Saeed, Bettina Weigelin, Christian M. Schürch

## Abstract

The bone marrow microenvironment (BMME) is essential for hematopoiesis and immunity, yet spatiotemporal single-cell analysis during leukemogenesis remains challenging. We characterized the BMME in femurs from wild-type and chronic myeloid leukemia (CML) mice at 7, 14 and 21 days post-induction by highly multiplexed and 3D microscopy. Using a 54-marker CODEX panel, we profiled 2,033,725 cells in 55 tissue regions of interest and identified 41 cell types through unsupervised clustering and supervised annotation. During leukemic progression, we observed an expansion of myeloid and progenitor cell populations, increased PD-L1^+^ leukemic cells, the upregulation of PD-1 on CD4^+^ and CD8^+^ T cells, and a profound loss of B cells, plasma cells and bone cells. Advanced CML exhibited a striking expansion of immature, pericyte-deficient vasculature that disrupted vascular niches and impaired hematopoietic stem/progenitor cell positioning. Spatial mapping revealed leukemia-specific cellular neighborhoods enriched in PD-1^+^CD8^+^ T cells, suggesting localized immune cell exhaustion. Early-stage CML showed increaseds between plasmacytoid dendritic cells and megakaryocytes, whereas advanced CML featured heightened megakaryocyte emperipolesis of non-leukemic granulocytes. Megakaryocytes were morphologically irregular in CML mice and BM trephine biopsies from CML patients. Laser-capture microdissected megakaryocytes from newly diagnosed CML patients had reduced expression of cytoskeleton genes, which was reversed in advanced cases treated with tyrosine kinase inhibitors. 3D imaging revealed vascular disorganization and depleted megakaryocytes in the diaphysis, underscoring region-specific pathology. Together, this study provides a spatiotemporal single-cell atlas of the BMME during leukemic progression, showing how leukemic cells reprogram the niche to support their expansion and immune evasion.

## Introduction

In adult mammals, hematopoiesis occurs in the bone marrow (BM), a complex organ consisting of diverse cell types (CTs) that can be broadly categorized as hematopoietic and non-hematopoietic cells^1^. Hematopoietic cells originate from multipotent hematopoietic stem/progenitor cells (HSPCs), while non-hematopoietic cells (osteoblasts, adipocytes, endothelial cells, pericytes etc.) originate from mesenchymal stem cells^1,2^. Cells from both compartments form the so-called HSPC niches. The BM niche orchestrates the homing, self-renewal, proliferation, and differentiation of HSPCs, ensuring the continuous generation of myeloid, lymphoid, and erythroid lineage blood cells. This tightly regulated process relies on dynamic and reciprocal interactions between HSPCs and their surrounding microenvironment^3^. Additionally, the BM functions as a critical immune organ, contributing to the priming of T cells in response to blood-borne antigens^4^ and playing a key role in shaping immunological memory^5^. Various mature T and B cell subsets, plasma cells, dendritic cells and differentiated myeloid cells within the BM contribute to hematopoiesis regulation by secreting cytokines and through as yet undefined cell-cell interactions^6^.

Chronic myeloid leukemia (CML) is a myeloproliferative disorder that originates from *BCR::ABL1*-transformed leukemic stem cells (LSCs)^7^. The advent of tyrosine kinase inhibitors (TKIs) targeting BCR::ABL1 has revolutionized the treatment of CML, enabling deep molecular remission in most patients^8^. The EURO-SKI study showed that in CML patients treated with TKIs for ≥3 years, nearly half remained in sustained remission up to 2 years after treatment discontinuation^9^. Despite these promising outcomes, TKI-insensitive LSCs persist in most patients over prolonged periods and remain a major barrier to a complete cure^10^. This persistence is closely linked to the BM microenvironment (BMME), which plays a pivotal role in leukemia pathogenesis by regulating and protecting LSCs. The interaction between leukemic cells and the BMME is bidirectional. It has been speculated that leukemic cells hijack and destroy normal HSPC-supportive niches, and transform the BM into leukemia growth-supportive niches^11^. This skewed BMME is associated with drug resistance and relapse, yet the underlying mechanisms and interactions remain incompletely understood. Therefore, characterizing the phenotypic, functional, and spatial properties of each cell within the BMME could facilitate a “two-pronged” treatment targeting both the leukemic cells and their interactions with the BMME to create an unfavorable environment for leukemia progression.

Despite extensive research, precisely mapping the localization of most CTs in the BM and defining their cellular properties and functions remains challenging^1,12^. A major obstacle lies in the limitations of conventional histology, as bone and BM structures are difficult to analyze due to artifacts introduced during sample preparation, including morphological alterations and autofluorescence, which can hinder unbiased quantitative assessment. Similarly, fluorescence-activated cell sorting (FACS), although widely used for high-dimensional single-cell profiling, lacks spatial context and cannot capture the anatomical organization of cells within the BMME. Advances in spatial proteomics, particularly CO-Detection by indEXing (CODEX)^13–15^, in combination with optimized tissue processing protocols, now overcome these limitations by enabling simultaneous visualization of over 50 biomarkers in intact BM sections at single-cell resolution.

In this study, we employed a 54-marker CODEX panel and 3D imaging to construct a spatiotemporal single-cell atlas of the BMME during CML progression. We found that leukemic cells remodel the BMME to create a leukemia-permissive microenvironment by driving T cells toward an exhausted phenotype and concentrating them within the leukemia niche. Leukemia also causes vascular remodeling, leading to immature blood vessels that fail to support hematopoietic stem/progenitor cells.

Additionally, CML megakaryocytes show morphological defects linked to alterations in cytoskeletal gene expression. Together, these findings reveal how leukemia creates immune-privileged niches and highlight vascular normalization and T cell rejuvenation as potential therapeutic approaches.

## Results

### Multiplexed CODEX imaging unveils the spatial landscape in mouse BM during leukemic progression

To comprehensively define the structure and composition of the BM and to characterize the remodeling of an unaltered BMME during leukemic progression, we induced a CML-like disease in immunocompetent mice using a well-established murine transduction/transplantation model. FACS-purified lineage-negative, Sca-1^+^c-kit^+^ (LSK) BM cells were transduced with a *BCR::ABL1-GFP* retroviral vector and injected i.v. into non-irradiated syngeneic mice^16–18^. Femoral bones were harvested from wild-type (WT) and CML mice at 7, 14 and 21 days after leukemia induction (n=3 mice per group) and analyzed using CODEX (**Fig. 1A**).

**Figure 1:**
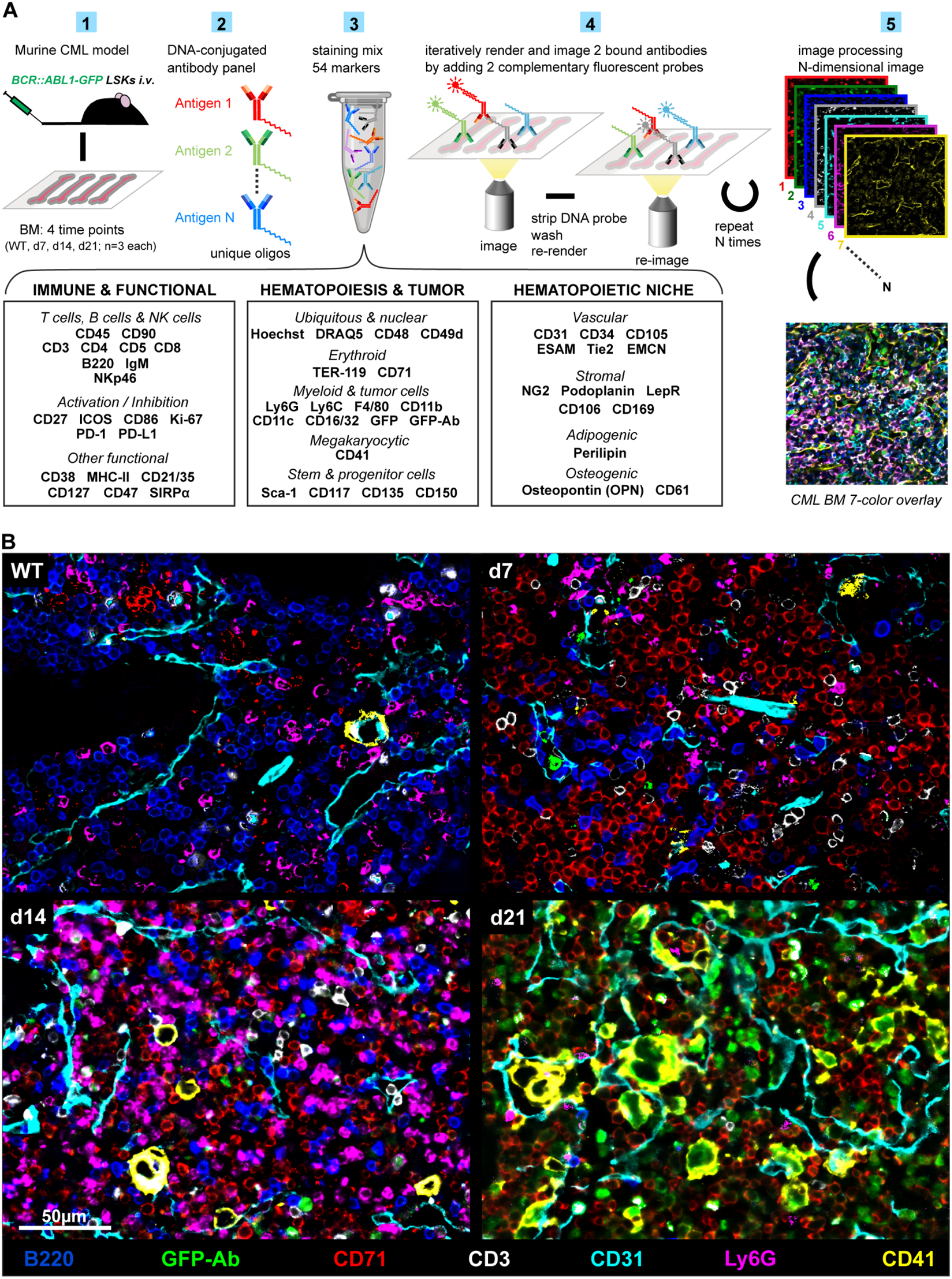
CML model and CODEX workflow. **(A)** (1) 3×10^4^ *BCR::ABL1-GFP* retrovirus-transduced, FACS-purified lineage^-^Sca-1^+^c-kit^hi^ (LSK) cells from naïve C57BL/6 (BL/6) mice were injected i.v. into non-irradiated recipient BL/6 mice. Femoral bones were harvested 7, 14 and 21 after injection, and naïve wild-type (WT) mice were used as controls (n=3 mice each). (2-3) A CODEX panel of 54 markers for fixed-frozen, decalcified BM was established and validated, followed by (4-5) imaging and data processing. **(B)** Representative CODEX seven-color overlay images comparing the four groups of mice. GFP-Ab, anti-GFP antibody. See also **Figure S1** and **Table S1, 4-6**. Scale bar, 50 µm.

We imaged 2-3 regions per bone (n=55 regions) with a 54-marker CODEX panel including markers for leukemia, immune, hematopoiesis, endothelial, and stromal cell components of the BM as well as functional markers such as PD-1, PD-L1, ICOS, and Ki-67 (**Fig. 1A, Table S1**). Following CODEX imaging, we validated the staining quality for each marker in every region and performed morphological correlation between fluorescent markers and H&E staining, exemplified by B220 staining of the B cells, GFP staining of the leukemic cells, CD71 staining of the erythroblasts, CD3 staining of the T cells, CD31 staining of the vessels, Ly6G staining of the granulocytes and CD41 staining of the megakaryocytes (MKs) (**Fig. 1B**). Comprehensive validation of remaining markers is further illustrated in **Fig. S1A-H**, confirming the technical robustness of our staining.

These systematic validations established CODEX as a reliable platform for resolving the BMME’s spatial architecture at single-cell resolution. Therefore, our multiplexed imaging strategy not only successfully maps the complex topography of BM niches, but also provides a quantitative foundation for subsequent single-cell analyses and marker quantification across cellular compartments.

### CODEX imaging enables the spatial annotation of BM-resident cell types at an unprecedented resolution

Our CODEX imaging data analysis pipeline, encompassing nuclei segmentation and fluorescent marker quantification, yielded spatially resolved single-cell data for subsequent analyses (**Fig. 2A.1**). Cell types (CTs) were annotated through a combination of unsupervised clustering and supervised annotation based on biomarker expression density, tissue localization, and morphology (**Fig. S2A**). Following each clustering iteration, we validated results by mapping the clusters to CODEX fluorescent images based on their X/Y coordinates. Clusters conforming to the anticipated antigen distribution patterns were annotated; contrarily, those diverging underwent subsequent clustering iterations until precise cell definitions were achieved (**Fig. 2A.2**). This was followed by an in-depth analysis of CT frequencies between groups (**Fig. 2A.3**) and cell-cell spatial interactions (**Fig. 2A.4**).

**Figure 2:**
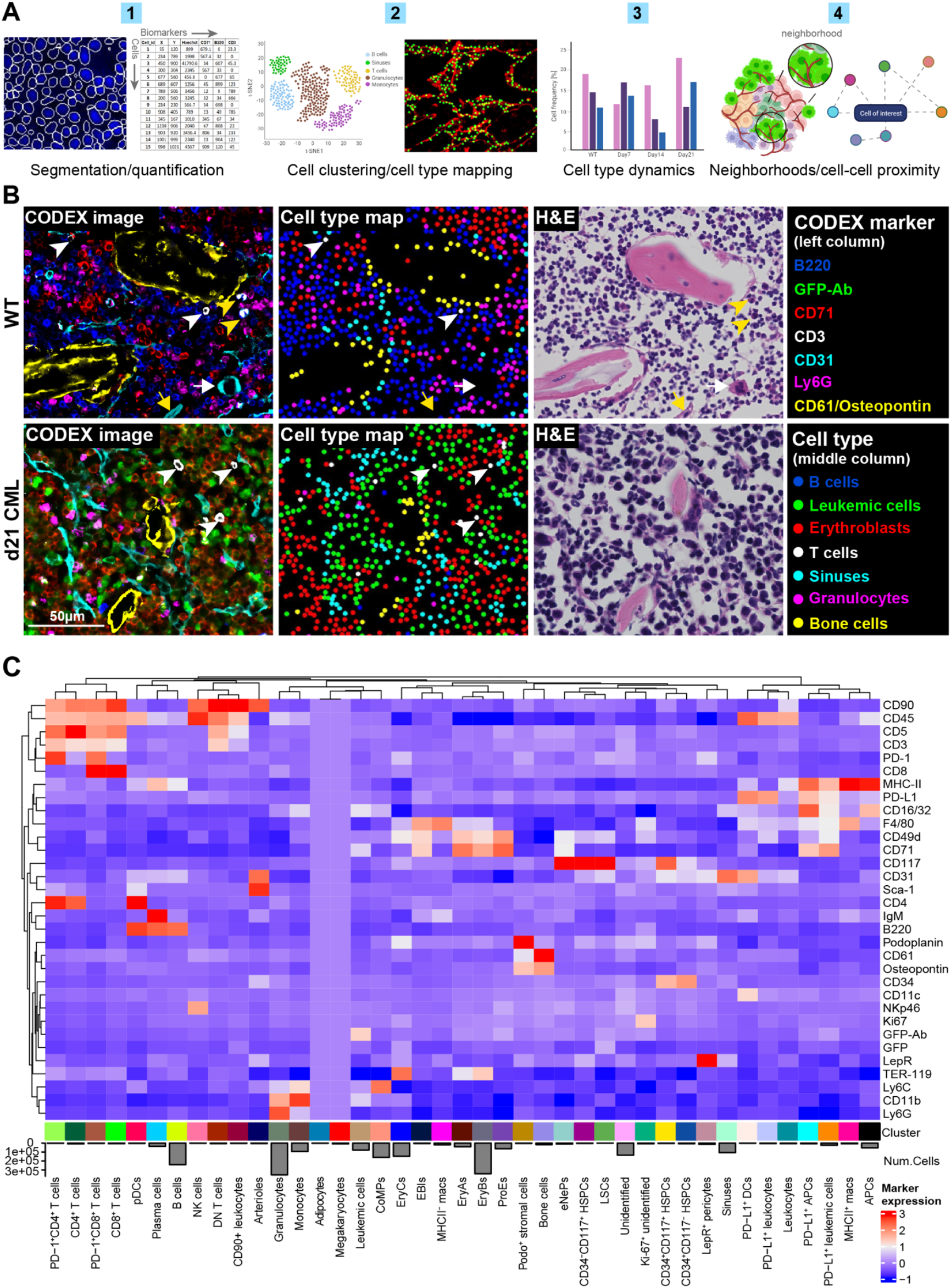
Identification and mapping of 41 cell types in the BM. **(A)** Overview of the CODEX data analysis workflow. (1) Cells were segmented, and marker intensities were quantified using DeepCell / Mesmer^20^. (2) Unsupervised clustering was performed using SpatialMap, based on 32 selected markers for CT identification, followed by supervised cluster / CT annotation. (3) CT dynamics and (4) spatial features were compared in all groups of mice during leukemic progression. **(B)** Comparison of fluorescence images and cluster maps. Left column: CODEX seven-color overlay images from WT (top row) and d21 CML (bottom row) mice. Middle column: corresponding CT maps. Right column: corresponding H&E-stained images. White arrow: MK positive for CD31; white arrowheads: T cells; yellow arrow: arteriole; yellow arrowheads: eosinophils (these cells unspecifically bind to multiple markers). **(C)** Standardized mean fluorescence intensities of the 32 individual markers selected for CT annotation, and numbers of cells for each annotated CT. Adipocytes and MKs were annotated manually and therefore no fluorescence intensity values are depicted. Scale bar, 50 µm. See also **Figures S2, S3** and **Table S2**. Parts of the figure were created with BioRender.com.

A total of 2,033,725 cells were identified and annotated across the four groups of mice (**Fig. S2B**). By mapping annotated, color-coded clusters onto tissues, both CODEX images (**Fig. 2B**, left) and H&E stainings (**Fig. 2B**, right) confirmed accurate assignments of CTs shown in the corresponding CT maps (**Fig. 2B**, middle). Additionally, we generated a “cell passport” for each CT based on the expression of CT-defining markers and appropriate negative controls (**Fig. S3A-X**). Thereby, we identified and verified 41 CTs (**Figs. 2C, S2A, S3A-X, Table S2**). As expected, using their endogenous GFP signal and by adding an anti-GFP CODEX antibody, we identified leukemic cells (GFP^+^GFP-Ab^+^; **Figs. 1B, 2B-C, S3A**), alongside rare LSCs (Lin^-^ CD117^+^/CD34^+^GFP^+^GFP-Ab^+^; **Figs. 2C, S3B**) in the BMME of leukemic mice, but not of WT mice. We also annotated various myeloid cell populations, such as non-leukemic granulocytes (CD11b^+^Ly6C^+^Ly6G^+^), monocytes (CD11b^+^Ly6C^+^Ly6G^-^), PD-L1^+^ dendritic cells (DCs; CD11c^+^MHC-II^+^), plasmacytoid DCs (pDCs; CD3^-^CD4^+^B220^+^Ly6C^lo^CD11c^lo^)^19–21^, macrophages (F4/80^+^), and antigen presenting cells (APCs; CD16/CD32^+^MHC-II^+^) (**Figs. 2C, S3C-G**). In addition, our data uniquely captured two rarely reported lineage-committed progenitors: common monocyte progenitors (CoMPs; lin^-^CD11b^-^Ly6C^+^Ly6G^-^)^22^ and early neutrophil progenitors (eNePs; lin^-^ CD117^+^CD71^+^CD49d^+^CD38^-^), which were recently identified in human BM^23^ (**Figs. 2C, S3H-I**). Moreover, in lin^-^ cells, we identified three distinct HSPCs populations: CD34^+^CD117^-^, CD34^-^CD117^+^ and CD34^+^CD117^+^ (**Figs. 2C, S3J**).

Erythroid progenitor lineages were categorized into four distinct subsets: proerythroblasts (ProEs; CD71^+^TER-119^-^), erythroblasts A and B (EryAs, EryBs; CD71^+^TER-119^+^), and erythroblasts C (EryCs; CD71^-^TER-119^+^). EryA and EryB clusters were distinguished by cell size and population size, with EryA larger but having a larger population^24–26^ (**Figs. 2C, S2C-E, S3K**). Erythroblastic islands (EBIs) were identified by a central macrophage (F4/80^+^) with surrounding erythroblasts (CD71^+^) (**Figs. 2C, S3L**).

MKs were identified manually based on their morphology and the expression of CD41 (see **Methods**; **Figs. 2C, S2F, S3M**). In the lymphoid compartment, we identified B cells (B220^+^IgM^-^), plasma cells (B220^+^IgM^+^) (**Figs. 2C, S3N**), NK cells (CD3^-^NKp46^+^) (**Figs. 2C, S3O**), CD4^+^ T cells (CD3^+^CD4^+^), CD8^+^ T cells (CD3^+^CD8^+^), double negative (DN) T cells (CD3^+^CD4^-^CD8^-^), as well as PD-1^+^CD4^+^ and PD-1^+^CD8^+^ T cells (**Figs. 2C, S3P-Q**).

Among the mesenchymal lineages, we identified two types of blood vessels: arterioles (CD31^+^Sca-1^+^) and sinuses (CD31^+^Sca-1^-^) (**Figs. 1B, 2B-C, S3R**). The sinuses were covered by rare leptin receptor (LepR)-expressing pericytes (**Figs. 2C, S3S**); specialized cells located on the walls of vessels and play a function of trophic support to endothelial cells^27,28^. Interestingly, however, LepR^+^ pericytes were not closely spatially associated with arterioles (**Fig. S2G**). Bone cells were characterized by positivity for CD61^29^ and/or osteopontin (**Figs. 2B-C, S3T**). Adipocytes were identified based on morphology and positivity for perilipin, and were annotated manually (see **Methods; Figs. 2C, S2H, S3U**). Furthermore, we identified Podoplanin-expressing (Podo^+^) stromal cells, a potential lymphatic endothelial progenitor CT (**Figs. 2C, S3V**)^30^.

Other CT clusters included leukocytes (CD45^+^), CD90^+^ leukocytes (CD90^+^CD45^+^), PD-L1^+^ leukocytes (PD-L1^+^CD45^+^) (**Figs. 2C, S3W**), unidentified proliferating cells (Ki-67^+^) and an unidentified cell cluster (**Figs. 2C, S3X**). These CTs were named due to their lack of expression of other lineage markers in our CODEX panel.

In summary, while aspirated BM material offers millions of cells for morphologic evaluation and single-cell analyses, these cells lose their spatial context due to disaggregation, and many CTs, like mesenchymal and endothelial cells, are notoriously difficult to aspirate. Moreover, in such analyses, CTs like adipocytes and MKs are often underrepresented. Our single-cell, spatially resolved CODEX dataset therefore represents a comprehensive murine BM tissue atlas, documenting the healthy BMME and characterizing it during leukemic progression, including many well-known and some previously less-documented BM-resident CTs.

### Leukemic cells remodel the BMME and establish a leukemia-permissive microenvironment

Having identified the various CTs in the BM, we next focused on the temporal changes in the BMME during leukemic progression. We grouped the 41 clusters into 13 categories by pooling data from the four groups of mice and captured a broad representation of cell populations, including myeloid cells (31.98%), erythroblasts (29.13%), B cells and plasma cells (13.63%), unidentified (6.63%), vessels (6.15%), leukemic cells (4.98%), APCs (2.99%), mixed CTs (1.42%), T cells (1.19%), bone cells (0.86%), stem/progenitor cells (0.60%), MKs (0.39%), and adipocytes (0.07%) (**Fig. 3A, Table S2**). When comparing the proportions across these 13 categories within the four groups of mice, B cell and plasma cell, leukemic cell and vessel proportions changed dramatically during disease progression, while the proportions of the other categories remained relatively constant (**Figs. 3B, S4A**).

**Figure 3:**
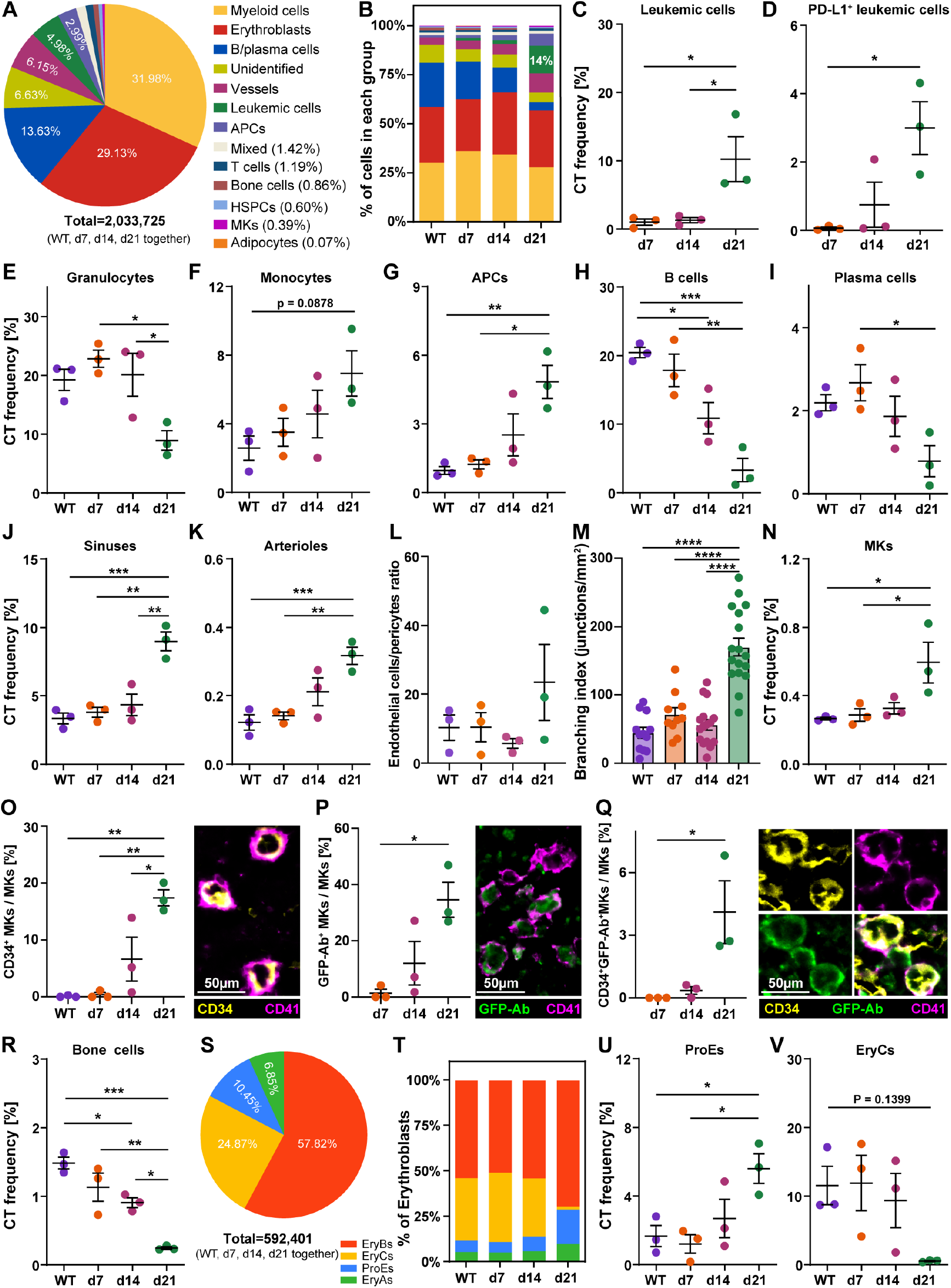
Dynamics of CT frequencies during leukemic progression. **(A)** Overall proportions of the 41 identified CTs consolidated into 13 categories across all groups of mice (n=2,033,725 cells). **(B)** Composition of the 13 categories between different groups (WT=446,729 cells; d7=391,748 cells; d14=595,650 cells; d21=599,598 cells). **(C-K)** Comparison of significantly altered CT frequencies across different groups. (**L**) Ratio of endothelial cells to pericytes. **(M)** Branching index of CD31-stained vessels, quantified using AngioTool. Each data point represents a tissue region (WT=12; d7=10; d14=16; d21=17). **(N)** Frequencies of MKs. **(O-Q)** Quantification of (O) CD34^+^ MKs, (P) GFP^+^ MKs, and (Q) CD34^+^GFP^+^ MKs as a proportion of total MKs across different groups. **(R)** Frequencies of bone cells. **(S)** Composition of the four erythroblast subsets across all groups combined (n=592,401). **(T)** Inter-group comparisons of erythroblast subsets (WT=127,109 cells; d7=103,261 cells; d14=188,305 cells; d21=173,726 cells). **(U-V)** Frequencies of ProEs and EryCs. APCs, antigen-presenting cells; HSPCs, hematopoietic stem/progenitor cells; MKs, megakaryocytes. Scale bars, 50 µm. See also **Figures S4, S5, S6** and **Table S2**. Data are presented as mean ± SEM. Statistics: One-way ANOVA (Tukey’s post hoc test). *p<0.05, **p<0.01, ***p<0.001, ****p<0.0001.

In a detailed comparison of CTs between groups, we found a robust expansion of both leukemic cells and PD-L1^+^ leukemic cells during leukemic progression (**Fig. 3C-D**). Additionally, there was a trend towards an increase in LSCs as the disease developed (**Fig. S4B**). These expansions were accompanied by a corresponding decrease in the proportion of non-leukemic granulocytes (**Fig. 3E**). Moreover, non-leukemic myeloid cells increased over time, including elevated levels of monocytes and APCs (**Fig. 3F-G**). There was also a noticeable tendency for increase in CoMPs, eNePs, macrophages, pDCs, PD-L1^+^ DCs and PD-L1^+^ APCs (**Fig. S4C-I**). In contrast, NK cells exhibited a reduction during leukemic progression, although this change was not statistically significant (**Fig. S4J**). Meanwhile, B cells and plasma cells were significantly reduced (**Fig. 3H-I**). These results are in line with previous reports demonstrating that CML promotes myelopoiesis at the expense of B lymphoid cells^31^. Furthermore, the frequency of the three types of HSPCs did not differ significantly between the groups (**Fig. S4K-M**).

Consistent with previous reports^32–34^, we fond progressive endothelial cell hyperplasia in both sinusoidal and arteriolar compartments during CML progression (**Fig. 3J-K**), with endothelial cell numbers showing strong positive correlation with leukemic burden (**Fig. S5A**). Quantitative vascular morphometry further demonstrated a significant expansion of the vascular network in advanced CML, characterized by increased total vessel length and elongated average vessel dimensions (**Figs. S5B-C, S6A-D**). Paradoxically, LepR^+^ pericytes followed an opposite trend: their frequencies remained constant, leading to an increased endothelial-to-pericyte ratio (**Figs. 3L, S4N**). The ratio rose from ∼10:1 in healthy BM, a value consistent with reported ratios in normal tissues (typically ranging from 1:1 to 10:1)^35,36^, to ∼23:1 in advanced CML (**Fig. 3L**), recapitulating the pericyte-deficient phenotype characteristic of immature vasculature^27,28,37,38^. This pathologic angiogenesis was further evidenced by a 4-fold elevation in vascular branch points (**Fig. 3M**), correlating with the disorganized branching patterns typifying structurally compromised vessels^39,40^. These findings collectively demonstrate that CML-driven BM neovascularization advances by promoting the development of structurally immature vascular networks. These networks are characterized by pericyte deficiency and aberrant branching patterns.

Along with the increase in leukemic cells and vessels in the BM, we noted a substantial increase in the abundance of MKs (**Fig. 3N**), which were positively correlated with vascular cells (**Fig. S5D**) and leukemic cells (**Fig. S5E**). Notably, CD150^+^ MKs, indicative of mature MK progenitors^41^, remained consistently abundant across all groups, accounting for approximately 50% of the total MK population (**Fig. S5F**). Interestingly, MKs more frequently expressed CD34 in CML mice, a phenotype that is consistent with immature MK progenitors^42–44^, with higher proportions of CD34^+^ MKs and CD150^+^CDCD34^+^ MKs at later time points of the disease (**Figs. 3O, S5G**).

In advanced CML, a substantial proportion of MKs also expressed GFP and were derived from the leukemic clone (**Figs. 1B, 3P**); however, only a small minority of MKs were GFP-Ab^+^CD34^+^ (**Fig. 3Q**). Quantification of marker-positive MK subsets was performed manually across all 55 analyzed regions. Moreover, bone cell numbers negatively correlated with leukemic cells (**Fig. S5H**), and were dramatically reduced in advanced CML (**Fig. 3R**), as has been previously reported in both mice and humans with leukemia^45–47^. Concurrently, the presence of Podo^+^ stromal cells, which consistently colocalized with bone cells in the CODEX images (**Fig. S5I**), exhibited a noticeable decline concomitant with the reduction in bone cell population (**Fig. S4O**).

The developmental trajectory from ProE to EryC delineates the maturation from erythroid progenitors to more differentiated erythroid cells in the BM^25,26^. Among the erythroblast populations, EryBs constituted the largest subgroup (57.82%), followed by EryCs (24.87%), ProEs (10.45%) and EryAs (6.85%) (**Fig. 3S**). Despite the advancement of leukemia, the overall proportions of erythroblast subsets remained relatively constant throughout disease progression (**Figs. 3B, S4A**). However, comparing the compositions of the erythroblast subpopulations across the four groups of mice, we noted a significant increase in immature ProEs cells and a near-complete loss of the most mature EryCs cluster in advanced CML (**Fig. 3T-V**). Although the intermediate erythroblast stages (EryA and EryB) also exhibited elevated proportions, these changes did not reach statistical significance (**Figs. 3T, S4P-Q**). These data suggest that the rapid proliferation of leukemic cells leads to an arrest in erythroblast maturation, potentially promoting the severe anemia observed in patients with leukemia^48^. For the remaining 7 CTs (EBIs, adipocytes, leukocytes, CD90^+^ leukocytes, PD-L1^+^ leukocytes, Ki-67^+^ unidentified, and unidentified), we did not observe significant changes during leukemic progression (**Fig. S4R-X**).

### Cellular neighborhood analysis decodes complex structures and identifies functional niches in the BMME

The BM is intricately structured, housing specific areas that create unique microenvironments. These microenvironments are crucial for selectively managing the regulation of different hematopoietic cells^1,49^. Our previous cellular neighborhood (CN) analysis pipeline strongly affirmed that examining the tumor immune microenvironment through the perspective of an intricate spatial architecture, instead of just as a collection of single cells, significantly improves our insight into functional subunits inside intact tissue^13,50^. Utilizing a “window” size of 14, that is, a cell and its 13 nearest neighbors, enabled the identification of 14 unique CNs within the BM through the aggregation of data from the four groups of mice (see **Methods**). Subsequently, each CN was annotated based on its unique enrichment of the 41 original CTs (**Figs. 4A, S7A-B**).

**Figure 4:**
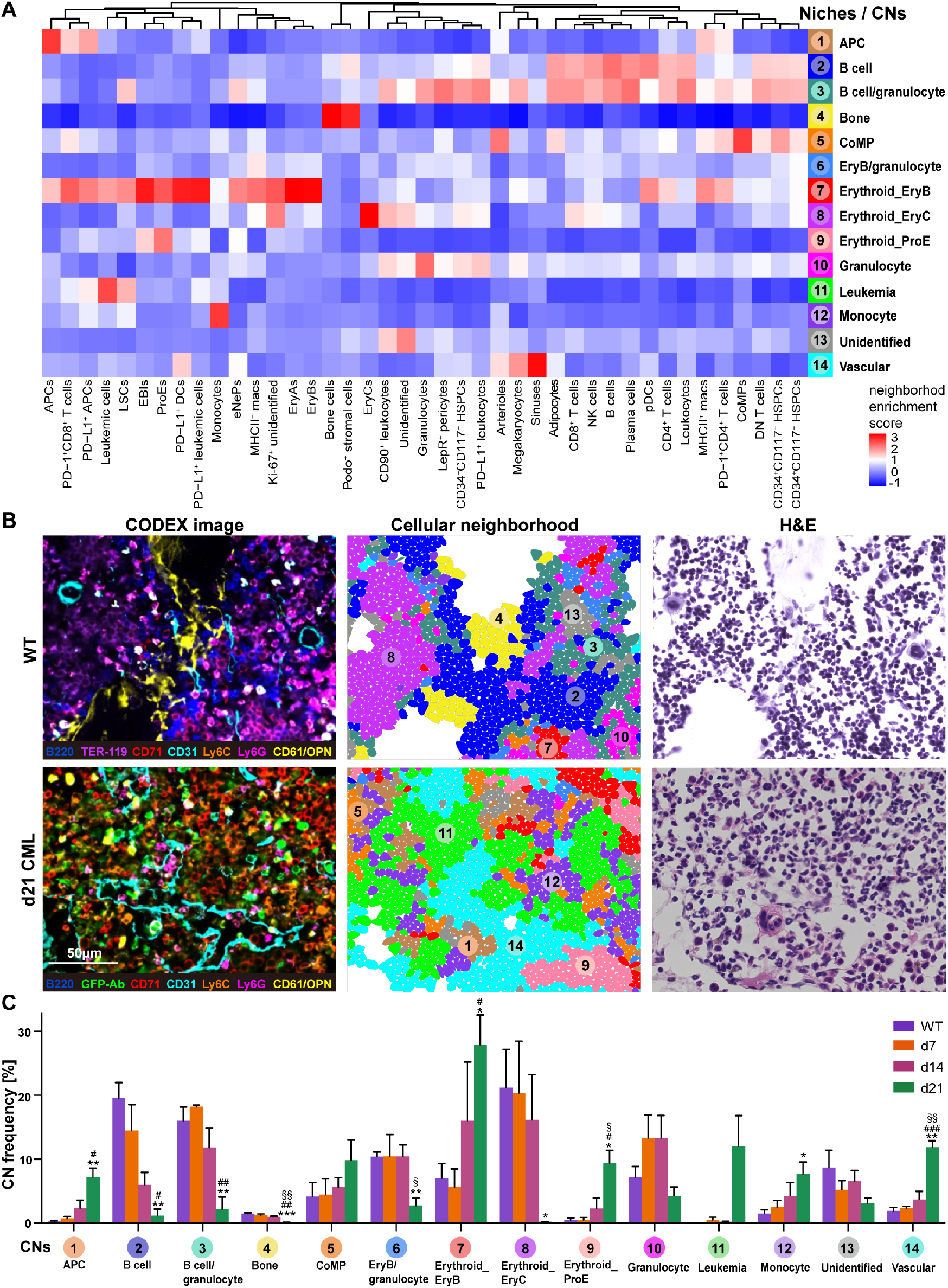
Dynamics of distinct BMME cellular neighborhoods during leukemic progression. **(A)** Identification of 14 BMME niches / CNs based on the local enrichment of the 41 original CTs (pooled data from all groups, normalized across columns). Neighborhood enrichment score indicates CT frequencies within each CN. **(B)** Representative CODEX images (left), Voronoi diagrams of CNs (middle) and corresponding H&E images (right) in WT and d21 CML mice. **(C)** Comparison of CN frequencies across the four groups. See also **Figures S7-S10**. Data are presented as mean ± SEM. Symbols *, ^#^, and ^§^ denote comparisons of the d21 group with the WT, d7, and d14 groups, respectively. Scale bar, 50 µm. Statistics: Student’s *t* test. *^/#/§^p<0.05, **^/##/§§^p<0.01, ***^/###^p<0.001.

The 14 CNs successfully mirrored structures that are directly associated with the architecture of the BM. The Voronoi maps of CNs aligned well with both CODEX fluorescent and H&E staining images (**Figs. 4B, S8**). Specifically, we identified a “leukemia” neighborhood CN-11, mainly composed of leukemic cells and only present in CML mice, especially at d21 (**Figs. 4A-C, S7A-B**). Myeloid cells were enriched in four different CNs: APC-enriched CN-1, CoMP-enriched CN-5, granulocyte-enriched CN-10 and monocyte-enriched CN-12 (**Figs. 4A-B, S7A-B, S8**). In line with the respective increases in APCs, immature progenitors and monocytes during leukemic progression (**Figs. 3F-G, S4C**), APC-enriched CN-1, CoMP-enriched CN-5 and monocyte-enriched CN-12 were more frequent in advanced CML mice (**Fig. 4C**). In contrast, the frequency of granulocyte-enriched CN-10 declined, mirroring the reduction in granulocyte abundance (**Figs. 3E, 4C**).

Erythroid precursors were enriched in four CNs: CN-6 contained EryB, granulocytes and B cells, and was decreased in advanced CML, reflecting the decreases of granulocytes and B cells (**Figs. 3E,H, 4A-C, S7A-B, S8**). CN-7, in which EryB was the predominant CT, was massively increased in advanced CML (**Figs. 4A-C, S7A-B, S8**). EryC-enriched CN-8, as well as ProE-enriched CN-9 were decreased and increased, respectively, consistent with the corresponding CT frequency trends (**Figs. 3U-V, 4A-C, S7A-B**).

B cells were most abundant in B cell-enriched CN-2 and the “B cell/granulocyte” CN-3 (**Figs. 4A-B, S7A-B**). These two CNs were dramatically reduced in advanced CML, reflecting the significant loss of B cells and healthy granulocytes, respectively (**Figs. 4C, 3E,H**). The mesenchymal CN-4 and CN-14 were predominantly enriched in bone cells and vascular endothelial cells (sinuses and arterioles), respectively (**Figs. 4A-B, S7A-B, S8**). In line with the loss of bone cells and the increase of vascular endothelial cells, bone CN-4 nearly disappeared in advanced CML, while vascular CN-14 increased significantly as CML progressed (**Figs. 3J-K,R, 4C)**. Additionally, CN-13, enriched in unidentified CTs and granulocytes, also tended to decrease with disease progression (**Figs. 4A-C, S7A-B, S8**). Furthermore, we illustrated the abundance of each CT across the 14 CNs, highlighting these specific localization patterns (**Fig. S9**). We visualized each BM region through CN Voronoi maps and compared the healthy BMMEs (**Fig. S10A**) vs. early and advanced CML (**Figs. S10B-D**). We noticed that while most BMs in early CML mice (d7 and d14) looked very similar to WT mice, leukemia onset was noticeable in some images. In one d7 mouse, we observed small hotspots of an altered BMME with the occurrence of leukemia CN-11 and EryB CN-7 (**Fig. S10B**, M6_reg1 and M6_reg2). At d14, one mouse exhibited two completely altered regions characterized by EryB CN-7 overgrowth (**Fig. S10C**, M9_reg2 and M9_reg3), whereas the two other regions (**Fig. S10C**, M9_reg1 and M9_reg4) of the same mouse showed almost no changes compared to WT mice. In advanced (d21) CML mice, all BMME regions were significantly altered compared to WT mice (**Fig. S10A**,**D**). Our findings of early CML development are in line with published studies indicating that leukemia onset begins in specific, localized patches within the BM before spreading, rather than being uniformly distributed throughout the BM^51,52^.

In summary, the BMME undergoes dramatic remodeling during CML progression: leukemia-specific niches emerge, erythroid zones expand, while B cell, granulocyte, and bone areas are lost. Concurrent vascular expansion creates a leukemia-permissive milieu that disrupts normal hematopoiesis.

### PD-1+CD8+ T cells increase during leukemic progression and are enriched in the leukemia CN-11

To evaluate changes in T cell subsets during leukemic progression, we categorized T cells into five subclusters: CD4^+^, CD8^+^, DN, PD-1^+^CD8^+^ and PD-1^+^CD4^+^ T cells. These subclusters constituted 35.03%, 25.33%, 20.66%, 12.08% and 6.90% of the total T cell population, respectively (**Fig. 5A,D**). Interestingly, although the overall T cell frequency remained consistent across different disease stages (**Fig. S11A**), we observed significant changes in the composition of T cell subsets between groups (**Fig. 5B-C**). Most notably, there was a significant expansion of PD-1^+^CD8^+^ T cells and PD-1^+^CD4^+^ T cells, accompanied by a corresponding decrease of (PD-1^-^) CD8^+^ T cells, while (PD-1^-^) CD4^+^ T cells remained stable (**Fig. 5B-C**). This progressive shift toward PD-1^+^ exhausted subsets suggests a leukemia-driven immunosuppressive polarization of the T cell repertoire.

**Figure 5:**
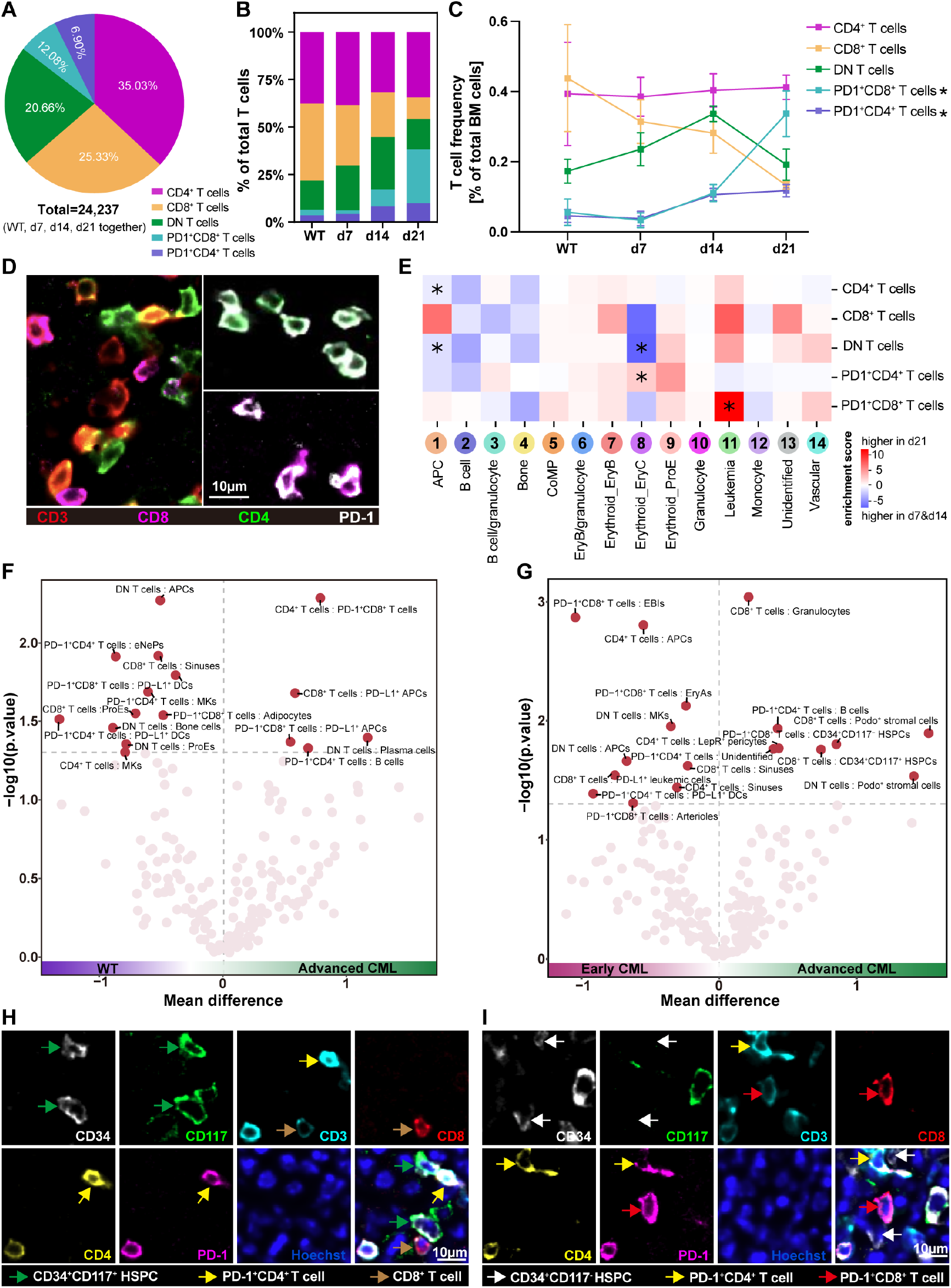
T cell subsets analysis. **(A)** Composition of the five identified T cell subsets within the combined groups (n=24,237). **(B)** Inter-group comparisons of T cell subset frequencies (WT=5,501 cells; d7=4,256 cells; d14=7,400 cells; d21=7,080 cells). **(C)** Changes in T cell subset frequencies during leukemic progression. **(D)** Representative CODEX images for T cell subsets. Left: CD3^+^CD8^+^ T cells, CD3^+^CD4^+^ T cells and DN T cells (CD3^+^ only, center of image). Right: PD-1^+^CD4^+^ T cells (upper panel) and PD-1^+^CD8^+^ T cells (lower panel). **(E)** Heatmap of estimated differential enrichment coefficients for T cell subsets in the respective CNs. A positive coefficient (red) indicates that a T cell subset is more enriched in the corresponding CN in advanced CML (d21); a negative coefficient (blue) denotes greater enrichment of a T cell subset in the corresponding CN in early CML (d7&d14). **(F-G)** Enrichment of cell-cell contacts between T cell and other CTs in (F) WT vs. advanced CML mice and (G) early vs. advanced CML mice. The x-axis shows the mean difference in contact frequency between groups, and the y-axis indicates –log_10_(p-value). A threshold of p < 0.05 (corresponding to - log_10_(p)>1.3) was used to define significant contacts. **(H-I)** Representative CODEX images of d21 CML BM displaying T cells in direct contact with CD34^+^CD117^+^ HSPCs and CD34^+^CD117^-^ HSPCs, respectively. Scale bars, 10 µm. See also **Figure S11**. Data are presented as mean ± SEM. Statistics: (C) Student’s *t* test; (E) Regression analysis; (F-G) Permutation-based test. *p<0.05.

To further investigate how these phenotypic changes manifest within in the spatial architecture of the BMME, we conducted a neighborhood enrichment analysis^13^ (see **Methods**) to determine when a specific T cell subset was “differentially enriched” across various CNs between the groups of mice. To evaluate CN-specific T cell subset frequencies, we applied a linear model that included group classification and overall subset frequency as covariates. The resulting estimated effects were visualized in a heatmap.

A significant coefficient in this model indicates enhanced enrichment of a particular T cell subset within a CN for one group compared to the other. Each of the five T cell subsets was significantly differentially enriched in at least one CN within either WT or advanced CML mice (**Fig. S11B**). Since there is no leukemia CN-11 in WT mice, we focused our analysis on comparing early (d7&d14) vs. advanced (d21) CML mice. In early CML, CD4^+^ T cells were enriched in CN-1 (APC), and DN T cells in CN-1 (APC) and CN-8 (EryC). In advanced CML, there was a significant enrichment of PD-1^+^CD4^+^ T cells in CN-8 (EryC), and particularly, PD-1^+^CD8^+^ T cells in the leukemia CN-11 (**Fig. 5E**). Taken together, these findings indicate that PD-1^+^CD8^+^ T cells not only expand during leukemic progression but also accumulate in the leukemia CN of the BMME, suggesting the creation of an immunosuppressive, leukemia-promoting BMME.

Building upon these spatial distribution patterns, we next examined the cellular interactions of T cell subsets by quantifying pairwise contacts with other CTs using a spatial permutation-based approach. Comparing WT mice to advanced CML mice, we found distinct contacts between T cells and other CTs. In WT mice, PD-1^+^CD4^+^ T cells and PD-1^+^CD8^+^ T cells preferentially contacted PD-L1^+^ DCs (**Fig. 5F**, upper left quadrant). Additionally, PD-1^+^CD4^+^ T cells also contacted eNePs and MKs, while CD8^+^ T cells and DN T cells contacted sinuses and APCs, respectively. In advanced CML, PD-1^+^CD8^+^ T cells contacted PD-L1^+^ APCs and CD4^+^ T cells, CD8^+^ T cells contacted PD-L1^+^ APCs, and PD-1^+^CD4^+^ T cells contacted B cells (**Fig. 5F**, upper right quadrant). Comparing early and advanced CML mice, we found that some of the cell-cell contacts observed in WT mice remained in early CML, but there were also new contacts, e.g., between CD8^+^ T cells and PD-L1^+^ leukemic cells (**Fig. 5G**, upper left quadrant). In advanced CML mice, PD-1^+^CD4^+^ T cells exhibited sustained contact with B cells compared with WT or early CML mice (**Fig. 5F-G**, upper right quadrant). CD8^+^ T cells preferentially contacted granulocytes and CD34^+^CD117^+^ HSPCs, while PD-1^+^CD8^+^ T cells selectively contacted CD34^+^CD117^-^ HSPCs (**Fig. 5G**, upper right quadrant), as unequivocally visualized in our high-resolution CODEX imaging (**Fig. 5H-I**). In summary, characterizing the cross-talk between T cells and other CTs in the CML BM may help understand the mechanisms by which T cells exert their anti-leukemic effects.

### Leukemic progression remodels HSPC niche localization, immune interactions, and functional dynamics in the BMME

HSPCs maintain the production of all hematopoietic and immune cells throughout life, and their function declines during aging, which is associated with changes in their characteristics^53–55^. Our findings prompted a deeper investigation into the three identified HSPC subsets. We analyzed a total of 12,249 HSPCs, categorizing them into CD34^+^CD117^-^, CD34^+^CD117^+^ and CD34^-^CD117^+^ (51.38%, 26.92% and 21.70%, respectively) populations (**Fig. 6A**). The frequencies of these subsets did not significantly differ among the four groups of mice (**Figs. 6B, S4K-M**). However, we observed a significant increase in cell size within the CD34^+^CD117^-^ HPSCs subset, the largest fraction among the three identified subsets, during leukemic progression (**Fig. 6C**). Large HSPCs were shown to have reduced proliferation potential and fitness compared to small HSPCs^55^, suggesting that enlarging CD34^+^CD117^-^ HSPCs gradually lose stem cell function in an increasingly leukemic BMME. Moreover, we observed that all three HSPC subsets displayed elevated levels of MHC-II expression in advanced CML mice (**Figs. 6D, S12A-C**). Recently, MHC-II^hi^ HSPCs have been identified in both mouse and human BM^56,57^, and higher MHC-II expression was linked to their quiescence^57^. In line with these findings, we observed that most MHC-II^hi^ HSPCs were negative for Ki-67 (**Figs. 6D, S12D**).

**Figure 6:**
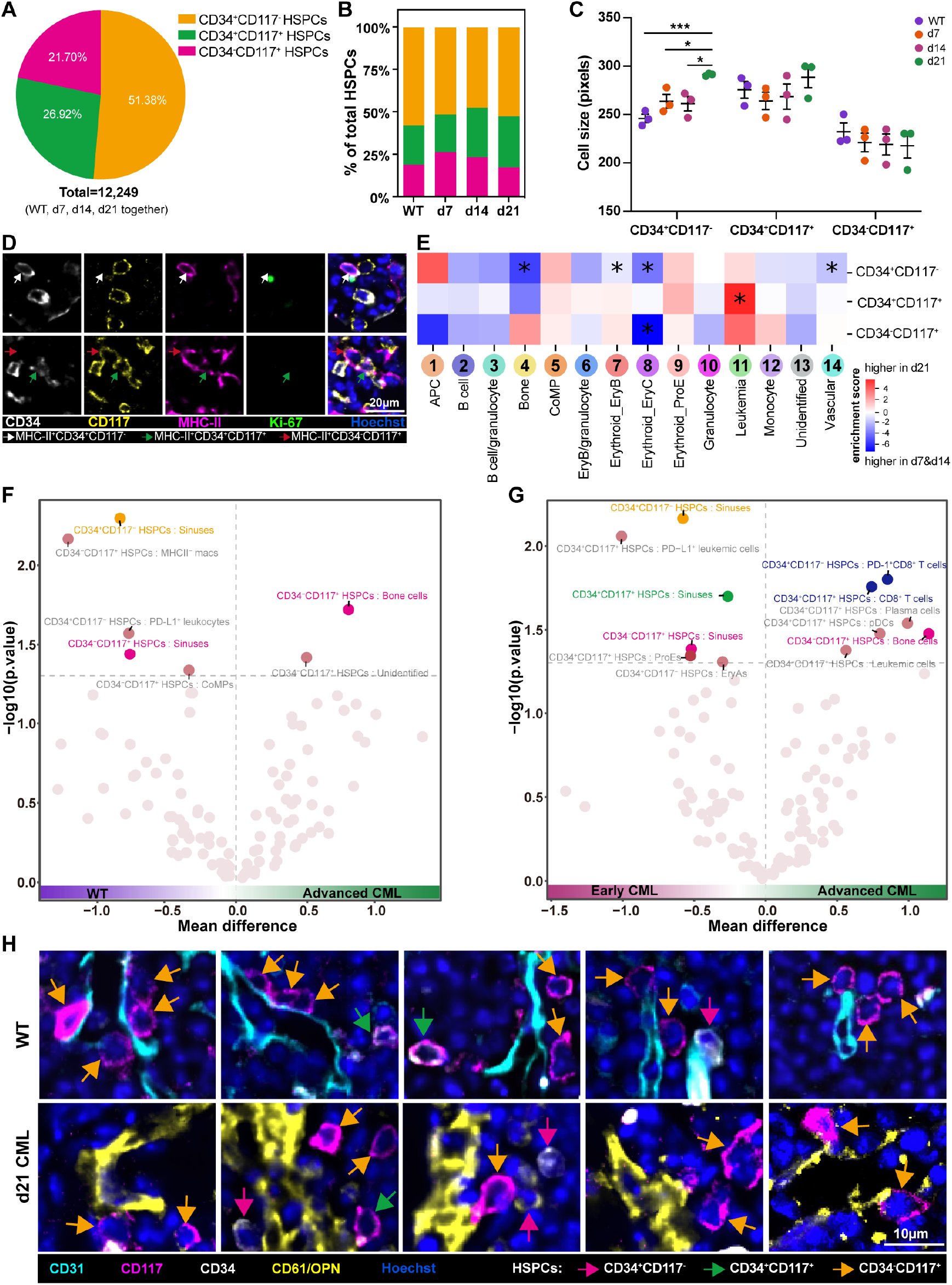
HSPC subsets analysis. **(A)** Composition of HSPC subsets (CD34^+^CD117^-^, CD34^+^CD117^+^, CD34^-^CD117^+^) across all groups combined (n=12,249). **(B)** Inter-group comparisons of HSPC subset frequencies (WT=2,456 cells; d7=2,453 cells; d14=4,328 cells; d21=3,012 cells). **(C)** Comparison of HSPC cell sizes across the four groups. **(D)** Representative CODEX images showing MHC-II^+^ HSPCs. The bright spot in Ki-67 (upper image) constitutes an antibody aggregate / staining artifact irrelevant to the analysis. **(E)** Heatmap of estimated differential enrichment coefficients for HSPC subsets in early vs. advanced CML mice. **(F-G)** Volcano plots showing the enrichment of cell-cell contacts between HSPC subsets and other CTs in (F) WT vs. advanced CML mice and (G) early vs. advanced CML mice. The x-axis shows the mean difference in contact frequency between groups, and the y-axis indicates –log_10_(p-value). A threshold of p < 0.05 (corresponding to –log_10_(p) > 1.3) was used to define significant contacts. **(H)** Representative CODEX images displaying HSPCs close to vessels in WT mice (upper row) and to bone structures in d21 CML mice (lower row). Scale bars, (D) 20 µm, (H) 10 µm. See also **Figure S12**. Data are presented as mean ± SEM. Statistics: (C) Student’s *t* test; (E) Regression analysis; (F-G) Permutation-based test. *p<0.05, **p<0.01, ***p<0.001.

Despite growing evidence for the role of vascular and bone cells regulating HSPCs^1,2,58^, little is known on how leukemic burden affects the localization of HSPCs in the BMME. We therefore investigated the differential enrichment of HSPC subsets across CNs in early vs. advanced CML mice. CD34^+^CD117^-^ HSPCs were more enriched in CN-4 (bone) and CN-14 (vascular), but also in CN-7 (EryB) and CN-8 (EryC) in the early CML BMME (**Fig. 6E**, top row). Interestingly, CD34^+^CD117^+^ HSPCs were more enriched in CN-11 (leukemia) in advanced compared to early CML (**Fig. 6E**, middle row). Moreover, CD34^-^CD117^+^ HSPCs were enriched in CN-8 (EryC) (**Figs. 6E**, bottom row). When comparing WT to advanced CML mice, only two HSPC enrichments were found: CD34^-^CD117^+^ HSPCs in CN-8 (EryC) and CD34^+^CD117^-^ HSPCs in CN-1 (APC) (**Fig. S12E**). These results suggest that the niches for HSPC subsets undergo significant alterations during leukemic progression.

To deepen our understanding of the relevant cell-cell contacts involved in these HSPC locational changes, we examined the contacts between HSPCs and other CTs in the BMME. Despite a significant expansion of the vasculature and a dramatic decrease of bone cells in advanced CML (**Fig. 3J**,**R**), which should provide HSPCs with increased opportunities to form contacts with vessels, we observed the opposite. While all HSPCs subsets maintained close cell-cell contacts with sinuses in either healthy BM (**Figs. 6F**, upper left quadrant, yellow and pink; **6H**, top row) and/or early CML (**Fig. 6G**, upper left quadrant, yellow, green and pink), these contacts were absent in advanced leukemia. Instead, CD34^+^CD117^-^ HSPCs were found to contact PD-1^+^CD8^+^ T cells, while CD34^+^CD117^+^ HSPCs were found to contact CD8^+^ T cells in advanced CML mice (**Figs. 5G-I, 6G**, upper right quadrant, blue). These bidirectional interactions between CD8^+^ T cells and HSPCs likely cause stem cell proliferation and differentiation^59^. Interestingly, CD34^-^CD117^+^ HSPCs in CML mice shifted to contact bone cells (**Fig. 6F-G**, upper right quadrant, pink; **6H**, bottom row). Coupled with a trend towards reduced cell size of CD34^-^CD117^+^ HSPCs in response to leukemic progression (**Fig. 6C**), this supports the notion that HSPCs in the bone niche are more quiescent^60,61^. These findings suggest that the immature, pericyte-deficient, leukemia-remodeled BM vasculature in advanced CML is unsuitable as an HSPC niche.

### Megakaryocyte interactions, and cytoskeletal changes during leukemic progression

MK, characterized by their large size and cellular fragility, present unique challenges for isolation and functional characterization in BM niches due to their low abundance and delicate nature. CODEX multiplexed imaging overcomes these technical limitations, which enabled comprehensive analysis of MK frequency, spatial distribution, morphological features, and microenvironmental interaction while preserving native tissue architecture. Our quantitative analysis revealed significantly increased MK populations in advanced CML (**Fig. 3N-Q**), promoting detailed investigation of their characteristics. Based on visual inspection of CODEX images, we observed close physical contacts between pDCs and MKs (**Figs. 7A, S13A**), with quantitative spatial analysis demonstrating stage-dependent variation in these contacts. Specifically, pDC-MK contacts occurred at significantly higher frequencies in early CML compared to both WT and advanced CML (**Fig. 7B**). This conserved interaction pattern was clinically relevant, as dual immunohistochemical staining for CD61 (MK marker) and CD123 (pDC marker) confirmed similar pDC-MK contacts in human CML BM specimens (**Fig. 7C**). These results mirror prior observations that pDCs regulate MKs and platelet production by releasing type I interferon^62,63^. Moreover, we identified emperipolesis as a distinct MK interaction pattern^64^, characterized by an increased occurrence of MKs engulfing healthy granulocytes in advanced CML (**Figs. 7D-E, S13B**), with only rare leukemia-derived, GFP-Ab^+^ granulocytes (**Fig. S13B**, last row). While emperipolesis has been documented under physiological conditions^65^, it becomes markedly more pronounced in pathological states, including hematologic malignancies and myelofibrosis^66–68^. The functional significance of this enhanced emperipolesis in advanced CML remains to be elucidated.

**Figure 7:**
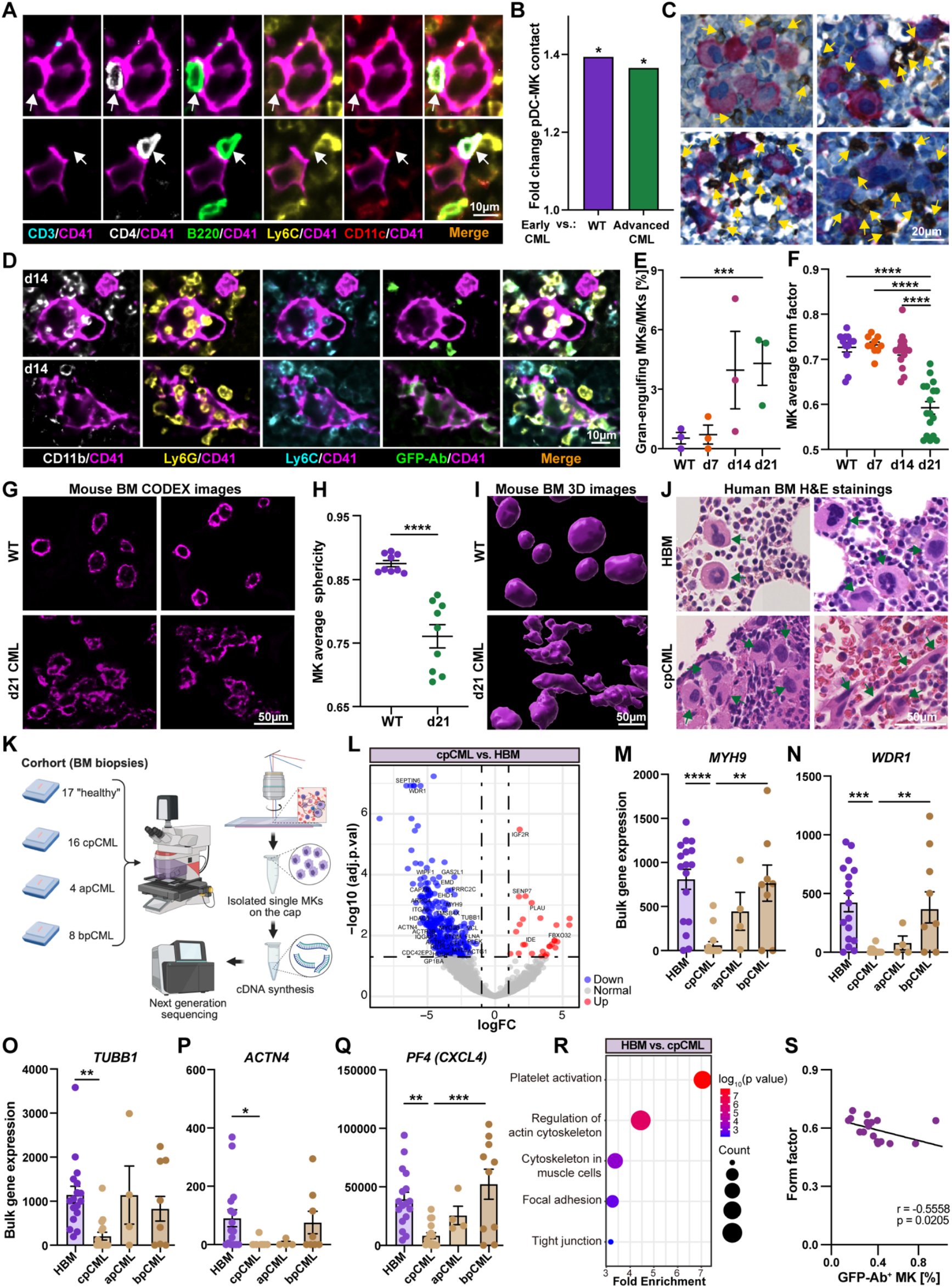
Changes in megakaryocyte cell-cell contacts and morphology during leukemic progression. **(A)** CODEX images showing close physical contacts between MKs and plasmacytoid dendritic cells (pDCs; defined as CD3^-^CD4^+^B220^+^Ly6C^lo^CD11c^lo^). White arrows indicate pDCs. **(B)** Quantification of pDC-MK contacts in early CML compared to WT and advanced CML. Bar graph shows fold change in pDC-MK contacts normalized to early CML. **(C)** Dual immunohistochemistry stainings of MKs (CD61, red) and pDCs (CD123, brown) in BM from human chronic-phase (cp) CML patients (representative images from n=4 patients in a stained cohort of n=48 patients). Yellow arrows indicate CD123^+^ pDCs. **(D)** Representative CODEX images showing granulocyte emperipolesis, with granulocytes internalized by GFP-Ab^-^ MKs (top row) and GFP-Ab^+^ MKs (bottom row). **(E)** Quantification of granulocyte-engulfing total MKs across groups. **(F)** CellProfiler^23^ analysis of the average form factor of MKs, with each data point representing a tissue region within the group. **(G)** Representative CODEX images of CD41 staining in WT and advanced CML mice indicating the shape of MKs. **(H)** Quantification of average sphericity of MKs in WT and advanced CML mice using 3D light sheet microscopy. Three mice per group were analyzed, with three identically sized regions selected per mouse. **(I)** Representative 3D images of femoral BM from WT and d21 CML mice. Surfaces were rendered based on CD42b expression to visualize the morphology of MKs. **(J)** Representative H&E-stained images of human healthy BM (HBM) and cpCML BM. Green arrows indicate MKs. (**K**) Schematic workflow of laser capture microdissection (LCM) followed by sequencing. apCML: advanced-phase CML; bpCML: blast-phase CML. **(L)** Volcano plot showing differentially expressed genes (DEGs) in laser-capture dissected MKs in cpCML vs. HBM samples. **(M-Q)** Normalized bulk expression values of selected genes in LCM-isolated MKs from HBM, cpCML, apCML and bpCML samples. **(R)** Bubble plot of Kyoto Encyclopedia of Genes and Genomes (KEGG) pathway enrichment analysis of downregulated DEGs in HBM vs. cpCML MKs. **(S)** Correlation of MK form factor and proportions of GFP-Ab^+^ MKs in advanced CML. Scale bars, (A, D) 10 µm, (C) 20 µm, (G, I, J) 50 µm. See also **Figures S13-14, and Table S3**. Data are presented as mean ± SEM. Statistics: (B) Permutation-based test; (E, H) Student’s *t* test; (F, M-Q) One-way ANOVA (Tukey’s post hoc test); (R) Wald test; (S) Linear regression analysis (Pearson’s correlation coefficient). *p<0.05, **p<0.01, ***p<0.001, ****p<0.0001.

Moreover, we observed distinct morphological differences in MKs between WT and advanced CML mice, as revealed by both CODEX and 3D light-sheet imaging. While WT MKs appeared largely spherical, MKs in advanced CML were shaped irregularly, making their boundaries difficult to define. This was reflected in a significantly reduced “form factor” metric of advanced CML MKs (**Fig. 7F**) and their ill-defined cell borders (**Fig. 7G**). Similarly, in 3D images obtained from light-sheet microscopy, their sphericity was reduced (**Fig. 7H-I**). Notably, these morphological abnormalities mirrored those observed in human CML BM biopsies (**Fig. 7J**), underscoring their clinical relevance.

To dissect the mechanisms driving these morphological changes, we performed laser-capture microdissection of single MKs from human FFPE BM sections, followed by RNA sequencing (**Figs. 7K, S14A-B**). The cohort included 17 “healthy” controls (see **Methods**), 16 chronic phase (cp), 4 accelerated phase (ap), and 8 blast phase (bp) CML samples (**Figs. 7K, Table S3**). Comparative transcriptomic analysis revealed profound transcriptional divergence between cpCML and healthy MKs, with 287 genes significantly downregulated and 28 upregulated in cpCML (**Fig. 7L**). Interestingly, 29 downregulated genes were linked to cytoskeletal regulation, most notably *MYH9* (*myosin heavy chain 9*)^69–72^, *WDR1* (*WD repeat domain 1*)^73,74^, *TUBB1* (*tubulin beta 1 class VI*)^75,76^, and *ACTN4* (*actinin α4*)^77^ (**Fig. 7M-P**). These genes are critical for cytoskeletal organization, actin filament assembly, cellular shape, structural integrity, and platelet function. Notably, *PF4 (platelet factor 4)*, also known as *CXCL4 (chem-okine [C-X-C motif] ligand 4)*, a small chemokine predominantly released by activated platelets and MKs^78^, was markedly downregulated in cpCML, suggesting impaired MK maturation and function (**Fig. 7Q**). While these genes were significantly downregulated in cpCML, their expression remained largely preserved in apCML and bpCML (**Fig. 7M-Q**). This finding could be attributed to differences in prior treatment history: 75% (9/12) of apCML/bpCML patients had received long-term TKI therapy before disease progression, whereas cpCML samples were collected at initial diagnosis prior to treatment. Consistent with this hypothesis, TKI-treated apCML/bpCML patients exhibited spherical MK morphologies (**Fig. S14C**), contrasting with the irregular shapes seen in untreated cases (**Fig. S14D**). KEGG pathway analysis of downregulated genes in cpCML further implicated dysregulation of platelet activation, actin cytoskeleton remodeling, muscle cell cytoskeletal function, focal adhesion, and tight junction pathways (**Fig. 7R**).

Increasing evidence suggests that the BCR::ABL1 oncoprotein plays a pivotal role in cytoskeletal abnormalities. Specifically, BCR::ABL1 associates with the cytoskeleton by interacting with actin via its COOH-terminal actin-binding domain, while several crucial tyrosine kinase substrates of BCR::ABL1 are cytoskeletal proteins^79,80^. We observed that the most-irregularly shaped MKs were leukemia-derived (**Figs. 1B, 3P**), with a significant inverse correlation between the proportion of GFP-Ab^+^ MKs and their form factor (**Fig. 7S**). This indicates that BCR::ABL1 promotes morphological irregularity of GFP-Ab^+^ MKs in our CML model. In line with this, Hekmatshoar et al. showed that TKI treatment of K562 leukemia cells upregulates myosin-encoding genes^81^, suggesting that TKIs reverse BCR::ABL1-induced cytoskeletal defects. While our findings establish that BCR::ABL1 disrupts cytoskeletal homeostasis and distorts MK morphology, the exact signaling aberrations require further investigation.

### Compartment-specific disruption of vascular architecture and megakaryocyte organization in leukemic BM

Different anatomical regions within the femur provide distinct microenvironmental cues that may influence vascular and MK morphology. To assess region-specific changes in CML, we analyzed whole-mount femurs from WT and advanced CML mice using tissue clearing and 3D light-sheet microscopy (**Fig. 8A-B**). In WT femurs, vessels were well-aligned along the femoral axis, and MKs were evenly distributed across both diaphyseal and epi-/metaphyseal regions (**Figs. 8A.1-2, Movie S1**). In contrast, advanced CML femurs showed severely disrupted vasculature in the diaphysis, with no discernible pattern and reduced MK presence (**Fig. 8B.1, Movie S2**). In the epi-/metaphyseal, MKs appeared abnormally clustered and spatially displaced (**Fig. 8B.2**).

**Figure 8:**
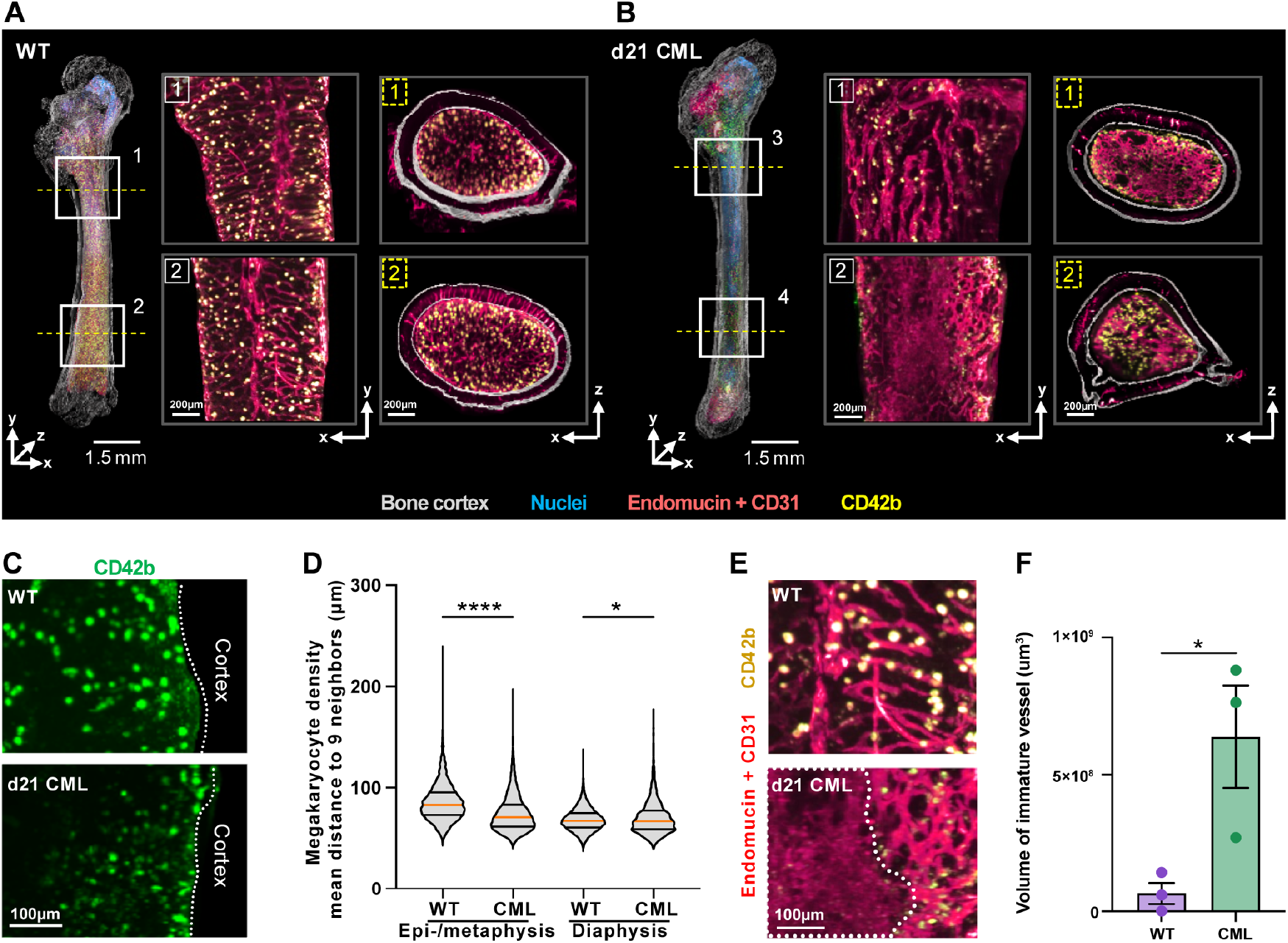
3D light sheet microscopy reveals regional disorganization of vasculature and megakaryocytes in leukemic femurs. **(A-B)** Representative 3D reconstructions of intact (A) healthy (WT, left) and (B) leukemic (d21 CML, right) mouse femurs. Insets and cross-sectional views highlight the vascular structure and MK distribution near the epi-/metaphysis and diaphysis. Colors represent cortical bone surface (grey), nuclei (blue), vessels stained with endomucin and CD31 (red), and MKs identified by CD42b staining (yellow). **(C)** Visualization of MKs (green, CD42b) in the lower femur shaft comparing healthy and leukemic conditions. **(D)** Truncated violin plots represent MK density as the average distance of each cell to its 9 nearest neighbors, with median (orange line) and quartiles (black line) indicated. Data were pooled from two femurs per condition (≥ 800 MKs analyzed per region). **(E)** Comparative visualization of vascular structures (red, endomucin and CD31) and MKs (yellow, CD42b) in healthy versus leukemic femurs. **(F)** Areas with disorganized (dotted line) versus organized vascular structures were classified using a machine learning-based classifier trained on representative image annotations. Quantification shows the volume of disorganized vascular areas per femur. Data are presented as mean ± SEM from three independent femurs per group. Scale bars, (A-B) 200 µm and 1.5 mm, (C, E) 100 µm. Statistics: (D) Kruskal-Wallis test, (F) Student’s *t* test. *p<0.05, ***p<0.001, ****p<0.0001.

We observed pronounced regional heterogeneity in MK distribution. In WT femurs, MKs were densely packed near the cortical bone in the diaphysis (**Fig. 8C**, top). In CML, MKs were distributed unevenly and showed a decreased density in the subregions of the diaphysis (**Fig. 8C**, bottom). Quantitative analysis of MK density, assessed by calculating the mean distances to nine neighboring cells, supported these observations. In the femoral epi-/metaphyseal region, MK in CML femurs were significantly more densely clustered compared to WT controls (**Fig. 8D**, left), as indicated by shorter mean intercellular distances. While statistical testing revealed a significant difference in MK spacing in the diaphysis between WT and CML groups (**Fig. 8D**, right), the effect size was minor, suggesting that MK density in this region differs in subregions but remains overall comparable. These findings indicate that, despite an overall increase in MK frequency in advanced CML (**Fig. 2N**), there are region-specific differences within the BM.

Additionally, advanced CML diaphysis showed extensive regions with either absent or aberrantly branched vasculature (**Fig. 8E**). Quantification of disorganized vascular volume confirmed significantly higher disruption in CML compared to WT (**Fig. 8F**). These observations highlight significant spatial heterogeneity in leukemic pathology and emphasize the influence of local BMME anatomical niches on the distribution and morphology of MKs as well as vascular remodeling.

## Discussion

Numerous pivotal studies have utilized single-cell RNA sequencing to identify and characterize components of the BMME^82,83^. However, only recently researchers have begun to incorporate spatial omics methodologies to study the BM and interrogate the spatial distribution and local intercellular communication networks within the BM^43,84^. A significant challenge in defining the single-cell composition of BM niches lies in isolating sufficient numbers of viable non-hematopoietic cells^85^. In this study, we overcame these limitations by using CODEX, which enables highly multiplexed tissue visualization with single-cell spatial resolution, to uncover leukemia-associated alterations in the BM of CML mice during disease progression.

We validated a 54-marker panel tailored for BM analysis, identifying and characterizing 41 CTs across WT and CML mice at days 7, 14 and 21. This comprehensive profiling captured both hematopoietic and non-hematopoietic populations, while mapping their spatial organization within the BMME. Each CT was rigorously confirmed based on morphology, tissue localization and marker expression, an approach that addresses key limitations of traditional unsupervised CT annotation, including susceptibility to errors from non-specific binding, autofluorescence, and fluorescence spillover. Among the 41 identified CTs, all were present in both WT and leukemic BM, except for leukemia-specific CTs (e.g., LCSs, leukemic CTs, GFP-Ab^+^ MKs), which were exclusively found in leukemic BM. We observed that CML development is associated with a striking myeloid expansion and reduction in B cells and plasma cells, indicating that CML drives myelopoiesis at the expense of B lymphopoiesis^31,86^. Previous research has demonstrated that leukemic cells secrete IL-6, which inhibits the differentiation of B lymphoid cells. Notably, blocking IL-6 with neutralizing antibodies or generating IL-6-deficient CML mouse models restored normal B-cell differentiation^86^. Similarly, bone cells were also significantly reduced in advanced CML, consistent with previous reports indicating that leukemic cells and metastatic tumor cells suppress osteoblast activity^45–47,87^. This suppression increases the risk of fractures and highlights the importance of incorporating bone health management into the comprehensive care of leukemia patients. Furthermore, as the disease progressed, we observed a pronounced differentiation arrest in non-leukemic cells, characterized by an increase in ProEs (erythroid progenitors), CD34^+^ MKs (MK progenitors), and a slight increase in CoMPs (common monocyte progenitors) and eNePs (early neutrophil progenitors). These findings may be explained by the secretion cytokines and growth factors by leukemic cells, such as TGF-β^88^ and IL-6^86,89^, which can alter the stromal niche and impair normal hematopoietic differentiation. For example, TGF-β has been shown to inhibit erythroid and megakaryocytic maturation while promoting an inflammatory microenvironment that favors leukemic cell survival and proliferation^90^.

Angiogenesis is widely recognized as an essential and early step in tumorigenesis, serving as a hallmark of solid tumors^91,92^ and playing a pivotal role in tumor progression and recurrence. While traditionally considered less relevant in hematological malignancies^34,93–95^, often referred to as “liquid tumors”, accumulating evidence, including our own findings, suggests that angiogenesis plays a more significant role in leukemia than previously appreciated. In advanced CML, we observed a marked expansion of both sinusoidal and arteriolar vessels, consistent with prior reports in hematologic cancers. This vascular remodeling is likely driven by the increased metabolic demands of proliferating leukemic cells and may facilitate their dissemination beyond the BM. Despite this increased vessel density, the number of LepR^+^ pericytes, critical for vessel stabilization and maturation, remained unchanged. The absence or detachment of pericytes can destabilize the vasculature, further promoting tumor cell dissemination^27,28^. Notably, whole-bone 3D imaging revealed severe regional disorganization of the vasculature, particularly in the bone diaphysis, where we observed large areas devoid of organized vessels or characterized by highly branched neovascular networks. This vascular disorganization coincided with the absence of all three types of healthy HSPCs from the vascular niche in advanced CML. A possible explanation for this phenomenon is that leukemic cells remodel the BM vascular system in a way that is unfavorable for HSPC colonization, potentially through the promotion of vascular immaturity.

These observations highlight the potential therapeutic value of targeting angiogenesis or employing vasculature normalization strategies in the treatment of leukemia.

Antileukemic T cell responses have been identified in patients with leukemia, and their role in disease development remains an area of active investigation^18,59,96,97^. In our study, while the overall frequency of T cells remained constant during leukemic progression, both CD4^+^ and CD8^+^ T cell subsets exhibited signs of immune exhaustion, characterized by the upregulation of the checkpoint molecule PD-1. Of particular interest, our CN analysis revealed that PD-1^+^CD8^+^ T cells were significantly enriched within the leukemia niche. Consistent with previous studies^16,98,99^, our findings further underscore the profoundly immunosuppressive BMME reconstructed by leukemic cells. Although TKI treatment has revolutionized the prognosis of CML patients, approximately 10% of patients develop resistance after 10 years of treatment^100^. At present, allogeneic hematopoietic stem cell transplantation (allo-HSCT) remains the only curative option for those with apCML or bpCML following TKI failure^100,101^. However, this approach is often impractical for elderly patients, who represent the majority of CML cases. PD-1-targeted immunotherapy has demonstrated clinical efficacy across a range of human cancers^102^. Moreover, its potential has been experimentally explored in murine CML models, where blockade of the PD-1/PD-L1 axis enhanced CD8^+^ T cell function and extended survival^16^. Our findings further underscore the rationale for investigating immune checkpoint inhibitors (ICIs) in combination therapies for relapsed CML or as consolidation treatment for patients discontinuing TKIs.

On the other hand, CD8^+^ T cells and PD-1^+^CD8^+^ T cells showed increased contacts with HSPCs in advanced CML. Bidirectional interactions between HSPCs and CD4^+^ T cells have previously been reported to drive stem cell proliferation, differentiation and eventual exhaustion^56^. As leukemia progresses, we observed a significant increase in the cell size of the largest population of HSPCs (CD34^+^CD117^-^), which has been reported as a potential indicator of declining cellular function^55^. Interestingly, we also observed elevated MHC-II expression in HSPCs under leukemia-induced stress. Notably, high MHC-II expression in HSPCs might has been associated with the acquisition of quiescence^57^. This suggests that under malignant stress, HSPCs may attempt to transition into a quiescent state as a protective mechanism to ensure survival.

CODEX analysis further provide direct evidence of close contact between pDCs and MKs, with pDCs known to play a crucial role in maintaining MK homeostasis^62,63^. In early CML, we observed increased pDCs-MKs contacts compared to WT and advanced CML mice. We hypothesize that pDCs are initially recruited to MKs as part of an effort to maintain MK homeostasis as the disease emerges. However, this regulatory mechanism appears to diminish over time, potentially due to the progressive breakdown of BMME organization as the disease advances. Additionally, we noticed an increased emperipolesis of non-leukemic granulocytes by MKs in advanced CML. Mechanistically, this interaction appears to be mediated by specific adhesion molecules. In both rat and mouse models, emperipolesis-active MKs upregulate intracellular adhesion molecule 1 (ICAM-1), while internalized neutrophils express its cognate receptor, lymphocyte function-associated antigen 1 (LFA-1; CD18/CD11a). Notably, blocking LFA-1 reduces lipopolysaccharide (LPS)-induced emperipolesis, highlighting the key role of ICAM-1/LFA-1 interactions in this process^65,103^. Despite these insights, the biological significance of this process remains unclear. It has been speculated that MKs could serve as a sanctuary for neutrophils in an unfavorable BMME or facilitate their trafficking out of the BM. Further investigation is needed to explore the underlying mechanisms of this process and its clinical significance.

While our CODEX data revealed an overall increase in the frequency of MKs in the BM of advanced CML mice, 3D imaging analyses uncovered pronounced regional heterogeneity in MK distribution. Specifically, the 3D reconstruction showed reduced MK density in the femoral diaphysis of advanced CML mice. This discrepancy may result from the anatomical focus of the CODEX imaging, which primarily captured sections from the femoral epi-/metaphysis. We also observed massive expansion of morphologically irregular MKs in both mouse and human CML samples. RNA sequencing of LCM-isolated MKs from cpCML patients revealed that this irregular morphology was correlated with the downregulation of cytoskeleton-related genes compared to “healthy” BM. Notably, in CML mice, these MKs with poorly defined boundaries were predominantly of leukemic origin, suggesting that the cytoskeleton abnormalities observed in MKs are caused by the BCR::ABL1 oncoprotein. This hypothesis aligns with existing studies demonstrating that BCR::ABL1 impairs cytoskeletal function in leukemia cell lines^79,104^. Notably, the subcellular distribution of BCR::ABL1 is considered crucial for its transforming ability, with at least 70% of the protein associated with the cytoskeleton^105^. These findings may also explain why cytoskeleton-related gene expression changes were not significant in apCML and bpCML samples, as most of the advanced CML patients had undergone prolonged TKI therapy prior to disease progression and biopsy. Since MK maturation and platelet production strongly depend on the dynamic remodeling of the cytoskeleton, investigating the specific signaling pathways disrupted by BCR::ABL1 will be pivotal in understanding the mechanistic underpinnings of these cytoskeletal abnormalities. Further research into these pathways could also provide new therapeutic avenues for mitigating MK dysfunction in CML.

In summary, our study highlights how leukemic cells strategically reprogram stromal, vascular, and immune components to establish a BMME that facilitates leukemic progression. By providing unprecedented, high-dimensional insights into the spatiotemporal dynamics of the BMME during leukemia development, this work serves as a valuable resource for immunologists, hematologists, and cancer researchers. The findings also offer important guidance for designing antibody panels and refining the annotation of CTs in future BM studies. For advanced CML patients, integrating standard therapies with innovative approaches, such as anti-angiogenic therapies, vascular normalization strategies, ICIs, or their synergistic combinations, may more effectively target leukemic cells and their supportive microenvironment, ultimately improving treatment outcomes.

### Limitations of this study

This study has several limitations. First, developing a BM-specific CODEX panel for fixed-frozen, decalcified samples is challenging. Despite testing multiple clones, antibodies for key CTs including regulatory T cells (FOXP3), γδ T cells (TCRγδ) and CXCL12-abundant reticular cells (CXCL12) could not be established. Moreover, although CD117 (c-kit) and Sca-1 were successfully stained, we were unable to confidently identify LSK cells, likely due to their rarity in tissue sections and the relatively weak Sca-1 expression on hematopoietic cells compared to arterioles. Second, CODEX imaging was focused on bone epi-/metaphyseal BM regions, potentially missing regional heterogeneity in CT distribution and vascular density. This was addressed for certain CTs using whole-bone 3D light-sheet imaging. Third, MKs and adipocytes were manually assigned, a labor-intensive process not easily scalable. Therefore, future studies should implement improved and larger antibody panels, optimize staining protocols, and implement advanced segmentation algorithms to further refine CT detection, annotation and identification.

## Methods

### Mice

C57BL/6J (BL/6) mice were purchased from the Jackson Laboratory (Bar Harbor, ME) and were housed in an American Association for the Accreditation of Laboratory Animal Care– accredited animal facility and maintained in specific pathogen-free conditions. The protocol was approved by Stanford University’s Institutional Animal Care and Use Committee under APLAC-15986. Experiments were performed with sex and age-matched (6-8 weeks) animals according to U.S. laws for animal protection.

### LSK cell isolation

Lin^-^, Sca-1^+^, c-Kit^+^ (LSK) cells were isolated as previously described^16–18^. Briefly, WT BL/6 mice were euthanized by CO_2_ inhalation and bones (femurs, tibiae, humeri, pelves and spines) were crushed using a sterile mortar and pestle in sterile phosphate-buffered saline (PBS; Life Technologies, #14190-250). The collected BM cells were centrifuged at 450g for 10 min and the pellet was suspended in 20 mL of ACK lysis buffer (Thermo Fisher Scientific, #A1049201) and incubated at room temperature (RT) for 5 min. After washing with PBS and centrifugation, cells were incubated with biotinylated antibodies targeting red cell precursors (anti-Ter119, BioLegend, #116203, 1:300), B cells (anti-CD19, BioLegend, #115503, 1:300), T cells (anti-CD3ε, BioLegend, #100304, 1:300), and myeloid cells (anti-Gr1, BioLegend, #108404, 1:300) in a final volume of 300 μL of magnetic-activated cell sorting (MACS) buffer (PBS supplemented with 5% fetal bovine serum [FBS; VWR Scientific, #16777-006] and 2 mM ethylenediaminetetraacetic acid [EDTA, PH 7.0; Teknova, #E0308]) for 20 min at 4°C. After washing with MACS buffer, cells were incubated with anti-biotin beads (Miltenyi Biotec, #130-090-485) for 20 min at 4°C, washed, resuspended, loaded onto LS columns (Miltenyi Biotec, #130-042-401) and separated on a QuadroMACS™ manual separator (Miltenyi Biotec, #130-091-051) according to the manufacturer’s instructions. Lin^-^ cells in the flow-through were stained with fluorescent antibodies (anti-Sca-1-APC, BioLegend, #108111, 1:100; anti-c-Kit-APC-Cy7, BioLegend, #105826, 1:300) in MACS buffer for 30 min at 4°C, washed, and sorted with a BD FACSAria III (BD Biosciences). The LSK cells were incubated in transplant media (RPMI [Life Technologies, #21870-092] supplemented with 10% FBS, recombinant murine stem cell factor [100 ng/mL; Peprotech, #250-03B], and recombinant murine thrombopoietin [20 ng/mL; Peprotech, #315-14A]) for 24 h at 37°C, 5% CO_2_.

### LSK cell transduction

The retroviral vector pMSCV-p210BCR/ABL-IRES-GFP and the packaging vector pIK6 were a gift from A. Ochsenbein (University Hospital Bern, Switzerland). Retroviral particles were produced and titrated as previously described^16^. A total of 1×10^5^ LSK cells were transfected twice on two consecutive days with the retroviral particles with polybrene (6.7 μg/mL; Sigma-Aldrich, #2938206) and 0.01 M N-2 hydroxyethylpiperazine-N′-2-ethanesulfonic acid (HEPES; Sigma-Aldrich, #83264) through spin infection (90 min at 1258g, 37°C). Subsequently, 3×10^4^ transduced cells were intravenously injected into non-irradiated syngeneic recipient mice. Leukemia engraftment was monitored by weekly tail vein bleeding and GFP expression in granulocytes was analyzed by FACS. Mice were euthanized by CO_2_ inhalation after 7-, 14-, and 21-days post-transplantation, and femurs were collected, cleaned of surrounding tissue, and processed as follows.

### Bone decalcification and embedding

Briefly, femurs were fixed for 4 h in freshly prepared 4% paraformaldehyde (PFA; Electron Microscopy Sciences, #15710) in PBS on ice. Subsequently, femurs were washed in ice-cold PBS three times for 5 min each. Decalcification was performed by incubating the bones in 0.5 M EDTA (pH 7.4; Sigma-Aldrich, #93302) at 4°C for 24 to 48 h with continuous agitation on a shaker. Following decalcification, femurs were washed three times in ice-cold PBS for 5 min each. To ensure optimal cryoprotection, the bones were immersed in PBS containing 20% (w/v) sucrose (Sigma-Aldrich, #S7903) and 2% (w/v) polyvinylpyrrolidone (PVP; Sigma-Aldrich, #P5288) at 4°C for 24 h. Then, each femur was embedded in optimal cutting temperature (OCT; Sakura, #2568), frozen on a metal block in liquid nitrogen, and stored at -80°C individually sealed in small Ziploc bags (SC Johnson).

### CODEX DNA-conjugated antibodies

Purified, carrier-free monoclonal and polyclonal anti-mouse antibodies (**Table S1**) were conjugated to maleimide-modified short DNA oligonucleotides (TriLink) (**Table S4**, column 2) as described before^13,15,106^. Specifically, each antibody was concentrated on a 50-kDa centrifugal filter column (EMD Millipore, #UFC505096) and sulfhydryl groups were activated in PBS containing 2.5 mM 2-carboxyethyl (Tris) phosphine hydrochloride (TCEP; Sigma-Aldrich, #C4706-10G) and 2.5 mM EDTA (pH 7.0) for 30 min at RT. The corresponding DNA oligonucleotide was resuspended in buffer C containing sodium chloride (NaCl; Thermo Fisher Scientific, #S271-10) at a final concentration of 400 mM and was added onto the filter with the reduced antibody at a 2:1 w/w ratio for 2 h at RT, with at least 100 μg of antibody per reaction. Conjugated antibody was washed and eluted in PBS-based antibody stabilizer solution (Thermo Fisher Scientific, #NC0436689) containing 500 mM NaCl, 5 mM EDTA, and 0.1% (w/v) sodium azide (NaN_3_; Sigma-Aldrich, #S8032). Each conjugated antibody was validated on BM tissue, and the optimal staining dilution was fine-tuned, commencing at a 1:100 dilution and adjusting in accordance with the signal-to-noise ratio.

### CODEX multiplex tissue staining

Coverslips were handled using Dumont coverslip forceps (Fine Science Tools, #11251-33). CODEX buffers (S1, S2, S4, blocking buffer and H2) were prepared as described^15^. Four femurs (WT and CML d7, d14, d21) were assembled into a tissue array inside a research cryostat (Leica Biosystems), sectioned at a thickness of 7 μm, and placed on 22 x 22 mm glass coverslips (Electron Microscopy Sciences, #72204-01) previously coated with poly-L-lysine (Sigma-Aldrich, #P8920). Immediately after melting the section by gently touching the undersurface, the coverslips were carefully immersed in pre-cooled 100% ethanol (Sigma-Aldrich, #E7023) in the cryostat for precisely 15 s. Subsequently, tissues on coverslips were rehydrated in buffer S1 twice for 2 min at RT each, followed by fixation in buffer S1 containing 1.6% (v/v) PFA for 10 min at RT. Coverslips were briefly washed in buffer S1 twice and equilibrated in buffer S2 for up to 30 min at RT. The coverslips were then transferred to a humidity chamber and incubated with 100 μl blocking buffer for 20 min at RT. After blocking, each coverslip was stained with antibody mix 1 (**Table S5**) in a final volume of 100 μL of blocking buffer overnight at 4°C in a humidity chamber on a shaker. Before staining, antibody mixes were pre-concentrated using a 50 kDa filter and subsequently adjusted to 100 μL with blocking buffer. Following the first round of staining, coverslips were washed in buffer S1 three times for 5 min at RT each, fixed in 1.6% PFA in buffer S1 for 10 min, and then incubated with 100 μL blocking buffer for 20 min in a humidity chamber at RT. Subsequently, each coverslip was stained with antibody mix 2 (**Table S5**) in a final volume of 100 μL of blocking buffer for 3 h at RT in a humidity chamber on a shaker. After the second round of staining, coverslips were washed in buffer S2 twice for 2 min each, fixed in buffer S4 containing 1.6% (v/v) PFA for 10 min, briefly washed in PBS three times, and incubated in ice-cold methanol (Thermo Fisher Scientific, #A412-4) for 5 min at 4°C. Following another three brief washes in PBS, tissue on coverslips were fixed using bis(sulfosuccinimidyl)suberate (BS^3^, 3 mg/ml; Thermo Fisher Scientific, #21580) in PBS for 20 min at RT, followed by another three brief washes in PBS, and stored in buffer S4 at 4°C.

### CODEX multicycle reaction and image acquisition

CODEX multicycle reactions and image acquisition were performed as described before^13,15^. Multicycle 96-well plates (Thermo Fisher Scientific, #07-200-762) were prepared by adding 400 nM fluorescent oligonucleotides (Integrated DNA Technologies) (**Table S4**, columns 3 to 5) in 250 μl of buffer H2 containing Hoechst 33342 (1:600; Thermo Fisher Scientific, #62249) and sheared salmon sperm DNA (0.5 mg/ml; Thermo Fisher Scientific, #AM9680), according to the multicycle reaction panel (for details on the order of fluorescent oligonucleotides and microscope light exposure times, see **Table S6**). Coverslips removed from buffer S4 were mounted onto custom-made acrylic plates (Bayview Plastic Solutions) using coverslip mounting gaskets (Qintay, #TMG-22). Subsequently, the tissue was stained with Hoechst 33342 (1:1000) for 1 min, followed by 3 washes with buffer H2. Acrylic plates were mounted onto a custom-designed plate holder and secured onto the stage of a BZ-X710 inverted fluorescence microscope (Keyence). Automated image acquisition and fluidics exchange were performed using a PhenoCycler instrument and driver software (Akoya Biosciences), according to the manufacturer’s instructions, with slight modifications. Tissue overview images were acquired manually using a CFI Plan Apo λ 2x/0.10 objective (Nikon), and automated imaging was performed using a CFI Plan Apo λ 40x/0.95 objective (Nikon) at standard resolution using Zeiss Immersol™ 518 F low fluorescence immersion oil (Thermo Fisher Scientific, #12-624-66A). DRAQ5 nuclear stain (Cell Signaling Technology, #4084L) was added and visualized in the last imaging cycle at a 1:100 dilution. After each multicycle reaction, hematoxylin and eosin (H&E) stainings were performed according to standard pathology procedures, and tissues were reimaged in bright-field mode.

### Computational image processing

Raw TIFF image files were processed, deconvolved, and background subtracted using the Enable Medicine cloud platform (https://www.enablemedicine.com/). After processing, the staining quality for each antibody was visually assessed in each tissue region using the Enable Medicine Visualizer (https://app.enablemedicine.com/portal/visualizer). Nuclear segmentation was performed using DeepCell / Mesmer^107^ based on the Hoechst nuclear stain on the Enable Medicine platform. Whole-cell segmentation masks were derived by dilating the nuclear masks for five rounds. In each round of dilation, each zero-valued pixel was changed to the value of a nearby segmentation mask, with probability proportional to the number of adjacent pixels in a 3×3 box that belong to that segmentation mask. Mean pixel intensity for each marker and corresponding X/Y coordinates were extracted from each segmented whole cell mask, and then stored in Enable Medicine’s cloud, enabling further downstream analysis.

### Unsupervised clustering and cluster validation

Cell phenotypes were identified using unsupervised clustering, in a workflow using the R package SpatialMap (https://docs.enablemedicine.com/spatialmap/). First, single cell expression values (mean pixel intensity for each marker) were pulled from the Enable Medicine atlas. Expression values were then normalized in a three step process. First, expression values were arcsinh transformed, with a cofactor determined by the 20^th^ percentile value of expression within each image. Second, expression values were centered to their mean value within each image and divided by the standard deviation within each image. Third, expression values were centered to their mean value within each cell and divided by the standard deviation within each cell. Each cell’s 30 nearest neighbors (NNs) in normalized expression space were computed by cosine distance. This NN network was then used as the input for leiden clustering. Clusters were manually annotated, and then cell phenotype labels were overlaid on the fluorescence image in the Visualizer (https://app.enablemedicine.com/portal/visualizer). Both morphological features and biomarker expression were validated during this process. Clusters that were found to be poorly annotated were separated out, re-clustered and re-analyzed in the fluorescence image until a comprehensive annotation set was achieved.

### Manual identification of adipocytes and megakaryocytes

Adipocytes and MKs had weak or unusually shaped / multiple nuclear signals, which were poorly captured by our segmentation approach. To remedy this, a manual approach was used to mark the locations of these CTs in the dataset. Each adipocyte and MK was manually identified in the Enable Medicine Visualizer (https://app.enablemedicine.com/portal/visualizer) based on their fluorescent signals (perilipin and CD41, respectively), coupled with their nuclei positions (Hoechst). Subsequently, the centroids of each manually identified cell were calculated and incorporated into our SpatialMap object for further comprehensive data analysis. In addition, subunits of over-segmented MKs obtained by clustering were removed from the dataset.

### Cellular neighborhood identification

Cellular neighborhoods (CNs) were identified as previously described^13^. For each of the 2,033,725 single cells in this experiment, a “window” of each cell and its 13 nearest spatial neighbors was captured, measured by Euclidean distance between X/Y coordinates. These windows were then clustered by their composition with respect to the 41 CTs previously identified by unsupervised k-means nearest neighbor clustering, followed by supervised annotation of the CNs to specific tissue regions. Specifically, each window was converted to a vector of length 41 containing the number of each of the 41 CTs among the 14 neighbors. Subsequently, windows were clustered using Kmeans from the stats module in R, with k = 16. Each cell was then assigned to the same CN as the window in which it was centered. All CNs were manually annotated according to the enrichment of the predominant CTs within them, validated by overlaying them onto the fluorescent images, and visualized by creating CN Voronoi diagrams. During this process, two pairs of CN clusters with very similar cell type compositions were merged, resulting in a final number of 14 CNs with a distinctive phenotype structure.

### Quantification of pairwise cell–cell contacts

Pairwise contact analysis identified preferentially co-located pairs of CTs. First, each cell’s contacts were defined as its 10 nearest neighbors in spatial coordinates. The number of contacts between each pair of CTs was enumerated within each sample. A spatial permutation-based approach was used to generate a null distribution of cell-cell contacts. For each image, the cell type labels in the spatial nearest neighbor network were randomized without changing the structure of the NN network, and the number of contacts between each pair of CTs was enumerated. These contacts were then summed across all images for each sample, to generate a single permuted value for each pair of CTs. This process was repeated for 100 permutations, and a normal distribution was fitted to the permuted values for each pair to obtain a null distribution. Sample-level enrichment p-values were computed as the upper tail of the null distribution. The log_10_ enrichment values were computed as the log of (true number of interactions / mean of the null distribution).

### Differential enrichment analyses

Linear models of the form Yn,c = β0 + β1X + β3Yc + ϵ were fitted, where Yc represents the log-transformed overall frequency of cell type c, X is a categorical variable indicating the patient group, and Yn,c is the log-transformed frequency of cell type c in CN n. The model includes coefficients βi, with ϵ as Gaussian noise with a mean of zero. A pseudocount of 1×10^−3^ was added before applying the logarithm transformation. The models were implemented using the statsmodels package in Python^108^, and the estimated coefficients along with p-values for β1 were extracted and visualized.

### Vascular analysis with AngioTool

AngioTool (version 0.6; National Institutes of Health, National Cancer Institute, Bethesda, MD, USA) is a free software designed for quantifying vascular structures^22^. To ensure accurate analysis, CD31-stained images were manually processed in ImageJ to remove CD31/CD41 double-positive MKs and background artifacts. The cleaned CD31 channel images were then analyzed in AngioTool, quantifying branching index, average vessel length, and total vessel length. The analysis procedure follows the methods described by Zudaire et al.^109^.

### Megakaryocyte morphological analysis with CellProfiler

To quantify the morphology of MKs, CODEX images stained for CD41 were pre-processed in ImageJ to remove background artifacts. The cleaned images were then analyzed using CellProfiler^110^ (version 4) following a standardized workflow. First, the IdentifyPrimaryObjects module was used to segment individual MKs, with the size range parameter carefully optimized to ensure accurate boundary detection while minimizing segmentation errors. Next, the MeasureObjectSizeShape module was applied to extract morphological features. Morphological regularity was assessed using the form factor, calculated as: form factor = (4π × Area) / Perimeter^2. Form factor values range from 0 to 1, where 1 represents a perfect circle, while lower values indicate increasing shape irregularity.

### Human sample collection

Archival human formalin-fixed paraffin-embedded (FFPE) BM biopsy samples were collected from the Department of Pathology, University Hospital Tübingen. Samples from chronic phase CML patients were collected at the time of diagnosis before therapy. Those from accelerated phase and blast phase CML were obtained either at first diagnosis or upon relapse following treatment. “Healthy” BM specimens were morphologically inconspicuous trephines obtained from patients who underwent BM evaluation due to suspected hematologic disorders or as part of lymphoma and extramedullary malignancy staging. Only samples without any pathological findings upon comprehensive diagnostic hematopathology assessment were included (**Table S3**). Informed consent was obtained in accordance with the Declaration of Helsinki protocol. This research has received approval from the Ethics Committee of the University of Tübingen (018/2024B02).

### Laser-capture microdissection (LCM)

Human BM FFPE blocks were cut into two serial 5 μm sections and mounted onto polyethylene naphthalate frame membrane slides (Thermo Fisher Scientific, #LCM0521) that had been irradiated faceup in an ultraviolet hood for 30 min. Each slide was then stored in a 50 mL tube (CELLSTAR, #227261) at -80°C until use. Prior to LCM, the slides were baked for 5 min at 60 °C. Then the slides were immersed in the following solutions sequentially: Xylene (2 min, two times; VWR Chemicals, #1330-20-7), 100% ethanol (1 min, two times), 95% ethanol (1min, two times), 70% ethanol (1 min, two times), nuclease-free water (1min; Thermo Fisher Scientific, #AM9932), hematoxylin (2 min; Diapath, #C0306), RNase free water (10 min), eosin (1 min; Diapath, #C0366), 70% ethanol (1 min), 95% ethanol (1 min), 100% ethanol (1 min, two times). After staining, the slides were loaded onto an Acculift™ LCM spatial profiler system (Targeted Bioscience) and the Acculift™ LCM Caps (Targeted Bioscience, #MED-2310-01) were placed over the regions of interest of the slides. MKs were identified based on their large size, single large oval, kidney-shaped or lobed nucleus with several nucleoli, and a low nucleus-to-cytoplasm ratio. 10-50 MKs per sample were dissected at 40x magnification using a finely focused ultraviolet laser and an infrared laser to adhere the selected membrane regions to the LCM cap. After cutting, the collected MKs were immediately prepared for further cDNA library preparation.

### Preparation of cDNA libraries

Libraries for sequencing were prepared using the Smart-3SEQ2 protocol (obtained from J. W. Foley, Stanford University School of Medicine) with slight modifications, which is an optimized version of the published Smart-3SEQ protocol^111^. Briefly, after LCM, 3 μL of lysis mix (**Table S7**) was added to the center of each LCM cap, where the tissue was located. The LCM caps were sealed with 0.5 mL tubes (Thermo Fisher Scientific, #Q32856) and incubated in an inverted position in a pre-warmed oven at 60°C for 3 h. Then, the tubes were centrifuged, and the ∼3 μL lysates were transferred to 0.2 mL low-retention polymerase chain reaction (PCR) tubes (Thermo Fisher Scientific, #11376044). The PCR tubes were placed in a pre-heated thermal cycler (Biometra TAdvanced), and the program proceeded as follows: 95°C for 1 min, 25°C for 1 min, then a 25°C hold. During the 25°C hold, the PCR tubes were removed, and the program continued with a hold at 42°C. During this time, 2 μL template-switching reverse-transcription (TSRT) mix (**Table S7**) was quickly added to the side of each tube. The tubes were then returned to the thermal cycler and continued with incubation at 42°C for 30 min, then 95°C for 30 s, and finally a hold at 10°C. For amplification, 4 μL PCR mix (**Table S7**), 0.5 μL universal P5 primer (AATGATACGGCGACCACCGAGATCT-ACACTCTTTCCCTACACGACGCTCTTCCGAT C*T) and 0.5 μL P7 primer labeled with unique index (CAAGCAGAAGACGGCATACGAGAT-[index]GTGACTGGAGTTCAGACGTGTGCTCT TCCGATC*T) were then added to each sample. 25 PCR cycles (for 31 to 50 MKs) or 27 cycles (for 10 to 30 MKs) were performed using specific PCR program (37°C for 6 min; 98°C for 45 s; 25 or 27 cycles at 98°C for 15 s, 60°C for 30 s, 72°C for 15 s; then 72°C for 60 s, and 4°C hold). Amplified cDNA was next purified with SPRI bead mix (Beckman Coulter, #B23317) and a magnetic separation block (ALPAQUA, #A32782). Finally, the samples were washed with fresh 80% ethanol and resuspended in a 10 μL DNA storage buffer (**Table S7**) to yield the sequencing-ready library.

### Library quantification and quality control

The library from each sample was quantified and assessed for quality using the Agilent 4150 TapeStation System with the High Sensitivity D1000 Kit (Agilent Technologies), which provides accurate measurement of cDNA concentration and fragment size. The final amplified libraries yielded approximately 10 μL per sample, with concentrations ranging from 1 to 150 nM and fragment sizes between 200 and 600 base pairs.

### Pool preparation and RNA sequencing

Libraries were pooled equimolarly, and final pools were checked for correct fragment length using an Agilent 2100 BioAnalyzer (Agilent Technologies). Sequencing pools were diluted to 0.5 nM and denatured according to the Illumina NextSeq 500 manual with addition of 5% PhiX spike-in. Pools were sequenced on an Illumina NextSeq 500 device using Illumina NextSeq 500 High Output Kit v2.5 (75 cycles) with a run mode 75,6,0,0 (75nt Read1 and 6nt Index read). The sequencing data were demultiplexed using the latest version of the Nextflow pipeline: nf-core/demultiplex^112^. The bcl2fastq conversion software was used for demultiplexing, and the quality was checked with fastp^113^. The average sequencing depth obtained was between 3.0 and 5.0 million reads per sample.

### Bulk RNA-seq analysis

Raw sequencing reads were quality-checked using FastQC (v0.12.1)^114^. Adapter sequences and low-quality reads were removed using Trim Galore (v0.6.10; https://github.com/FelixKrueger/TrimGalore). The filtered reads were then aligned to the human genome (GRCh38) using HISAT2 (v2.2.1)^115^; Samtools (v1.21)^116^ was used for sorting and bam conversion, while FeatureCounts from Subread (v2.0.6)^117^ was used to allocate reads to genes. Genes with zero counts in more than 40% of samples were excluded to ensure data robustness. Differential gene expression analysis was performed using DESeq2 package in Bioconductor^118^, with significant differentially expressed genes (DEGs) defined by adjusted p-values (padj < 0.05) and absolute log2 fold change (log_2_FC) > |1|^118^. To compare gene expression levels across different groups, RNA counts were normalized to transcripts per million by accounting for gene length. Enrichment analysis of DEGs was performed using the DAVID database (https://david.ncifcrf.gov/tools.jsp) to identify significantly enriched Kyoto Encyclopedia of Enrichment of Genes and Genomes (KEGG) pathways^119^.

### 3D whole-mount tissue clearing and antibody staining

OCT-embedded femurs were briefly thawed at RT for 2 min to allow rough trimming of the surrounding OCT. The trimmed samples were incubated in 4% PFA at RT for 20 min. Once the remaining OCT became transparent, it was gently removed from the femur using spatulas. Tweezers were not used to prevent tissue damage. Samples were then post-fixed overnight in 4% PFA at 4°C with continuous shaking. After fixation, femurs were washed three times with PBS for 30 min at 4°C. Decalcification was performed using 0.5 M EDTA for 6 days at 4°C, with the solution replaced every second day. After decalcification, samples were washed in PBS at RT three times for 30 min each. To remove residual heme, femurs were incubated in 20% N,N,N’,N’-tetrakis(2-hydroxypropyl)ethylenediamine (Quadrol; Sigma-Aldrich, #122262) in water at 37°C for 36 h, followed by three 30 min PBS washes at RT. Samples were incubated in urea buffer (25% urea [Sigma-Aldrich, #U5378], 15% glycerol [Sigma-Aldrich, #G5516], 15% Triton X-100 [Sigma-Aldrich, #X100] in VE-water [Stakpure, OmniaPure UV/UF Reinstwassersystem]) for 6 h at 4°C followed by washing 5 min in PBS at RT. To enhance tissue permeability, femurs were treated with 0.2% Collagenase A (Merck KGaA, #10103578001) in PBS for 1 h at 37°C, then washed twice with 2% FBS in PBS for 5 min at RT. For antibody staining, samples were incubated overnight at 37°C in blocking buffer (10% DMSO [Sigma-Aldrich, #D8418], 6% FBS in PBS containing 0.1% Triton X-100). Primary antibodies (anti-CD42b [Abcam, clone SP219, host rabbit, #ab183345, 1:50]; anti-CD31 [BD Biosciences, clone MEC13.3, host rat, #550274, 1:50] and anti-Endomucin [Santa Cruz Biotechnology, clone V.7C7, host rat, #sc-65495, 1:100]) and Sytox Green nuclear staining (Thermo Fisher Scientific, #S7020, 1:3000) were diluted in antibody diluent (5% DMSO, 3% FBS, 0.2% Tween 20, 0.05% sodium azide [Carl Roth, #K305.1] in PBS), and samples were incubated for 7 days at 37°C. Following primary antibody incubation, samples were washed six times for 1 h each and overnight in washing buffer (0.2% Tween 20, 100 µg/ml heparin sodium salt [Sigma-Aldrich, #H3393], 0.05% sodium azide in PBS). Secondary antibody staining (Donkey anti-Rabbit Alexa Fluor™ Plus 680 [Thermo Fisher Scientific, clone polyclonal, #A32802, 1:200] and Donkey anti-Rat DyLight™ 755 [Thermo Fisher Scientific, clone polyclonal, #SA5-10031, 1:200]) was performed for 6 days at 37°C, followed by washing six times for 1 h and once overnight in the washing buffer. After staining, samples were dehydrated in a graded ethanol series (pH > 9; 50%, 70%, 100%, 100%) with each step incubating 24 h at 4°C. For refractive index matching, femurs were incubated in ethyl cinnamate (ECi; Sigma-Aldrich, #W243000) at RT for at least 48 h. ECi was replaced twice, after 24 h each.

### Light-sheet microscopy image acquisition

Femurs were imaged using the Blaze Ultramicroscope II (Miltenyi Biotec) equipped with a 1x objective lens (NA 0.1) and an additional 2.5x magnification. Image acquisition was performed with a 4.2-megapixel sCMOS camera. Fluorescent signals were detected using the following excitation and emission filter combinations: 650/45 nm excitation and 720/60 nm emission for Alexa Fluor 680, 740/40 nm excitation and 780 nm long-pass emission for DyLight 755, and 470/40 nm excitation with 525/50 nm emission for Sytox Green nuclear staining.

### Light-sheet microscopy data analysis

Light-sheet images were stitched, processed and quantified using Imaris 10.1.1 (Bitplane AG). BM and cortical bone were segmented using the Imaris machine learning-based pixel classification tool. Foreground and background pixels were determined by training the model on all channels (nuclei, Endomucin + CD31, and CD42b) and surfaces were calculated with a surface grain size of 25 µm. MKs were counted using the Imaris spot detection tool. Using the CD42b channel, spot detection was performed with an estimated spot size of 23.6 µm and a spot quality filter set to 550. The same settings for spot detection were applied in all samples. The densities of MKs close to the femur epi-/metaphysis region and the diaphysis were determined by calculating the average distance of each spot to its 9 nearest neighbors within a volume of 9.7 x 10^8^ um^3^. The volume of regions with immature vessel structures were determined by training a classifier using the Imaris machine learning-based classification tool. The classifier was trained on representative healthy and immature vessel regions using the vessel-channel stained for Endomucin and CD31 positive endothelial cells. The classifier was then applied to the entire BM region in all femurs to determine the volume of immature vessel regions.

### Statistical analysis

All statistical analyses were performed with R (4.3.2) and GraphPad Prism v9 (GraphPad Software, Inc). Data are reported as means ± SEM. Comparisons between two groups were conducted using unpaired, two-tailed Student’s t test or Kruskal-Wallis test. For multiple group comparisons, one-way ANOVA followed by Tukey’s post hoc test was applied. Correlation analyses were performed using Pearson’s correlation coefficient for normally distributed data. Neighborhood enrichment analysis was evaluated using linear regression analysis to assess associations across groups. Cell–cell contact enrichment was tested using permutation-based methods, with significance determined by comparing observed contact frequencies to a randomized null distribution. Differential gene expression for volcano plots was assessed using the Wald test as implemented in DESeq2. Differences were considered statistically significant at p < 0.05. Only statistically significant comparisons are displayed in the figures, and all specific information is provided in the respective figure legends.

## Acknowledgments

C.M.S. was supported by the Faculty of Medicine, University of Tübingen; the Institute of Pathology, University Hospital Tübingen; the State of Baden-Württemberg within the Centers for Personalized Medicine Baden-Württemberg (ZPM); the Mach-Gaensslen-Stiftung Schweiz; the Swiss National Science Foundation (P300PB_171189; P400PM_183915); the Lady Tata Memorial Trust; the Dres. Bayer-Foundation; Swiss Cancer Research (KFS-5114-08-2020); the European Union (ERC, 101116768); the German Research Foundation (Cluster of Excellence iFIT, ‘Image-Guided and Functionally Instructed Tumor Therapies’, EXC 2180 - 390900677, University of Tübingen; INST 37/1228-1 FUGG; INST 37/1302-1 FUGG; INST 37/1310-1 FUGG); the Brigitte und Dr. Konstanze Wegener-Stiftung; the Werner Jackstädt-Stiftung; the Bundesinstitut für Risikobewertung (Bf3R, 60-0102-01.P641-12572660); and the Leukemia Research Foundation (1077776). L.L. and H.W. were supported by the China Scholarship Council (202108320058 and 202108340026, respectively). B.W. was supported by the German Research Foundation (Cluster of Excellence iFIT, ‘Image-Guided and Functionally Instructed Tumor Therapies’, EXC 2180 - 390900677, University of Tübingen) and the IZKF Junior Research Group grant (2561-0-0). G.P.N. was supported by the NIH (2U19AI057229-16, 5P01HL10879707, 5R01GM10983604, 5R33CA18365403, 5U01AI101984-07, 5UH2AR06767604, 5R01CA19665703, 5U54CA20997103, 5F99CA212231-02, 1F32CA233203-01, 5U01AI140498-02, 1U54HG010426-01, 5U19AI100627-07, 1R01HL120724-01A1, R33CA183692, R01HL128173-04, 5P01AI131374-02, 5UG3DK114937-02, 1U19AI135976-01, IDIQ17X149, 1U2CCA233238-01, and 1U2CCA233195-01); the DOD (W81XWH-14-1-0180 and W81XWH-12-1-0591); the FDA (HHSF223201610018C and DSTL/AGR/00980/01); Cancer Research UK (C27165/A29073); the Bill and Melinda Gates Foundation (OPP1113682); the Cancer Research Institute; the Parker Institute for Cancer Immunotherapy; the Kenneth Rainin Foundation (2018-575); the Silicon Valley Community Foundation (2017-175329 and 2017-177799-5022); the Beckman Center for Molecular and Genetic Medicine; Juno Therapeutics (122401); Pfizer (123214); Celgene (133826 and 134073); Vaxart (137364); and the Rachford & Carlotta A. Harris Endowed Chair. Sequencing and sequencingrelated workflows were performed with the support of the DFG-funded NGS Competence Center Tübingen (INST 37/1049-1). Data management and storage of raw data for this project were supported by the Quantitative Biology Center (QBiC), University of Tübingen, Germany.

## Author Contributions

Conceptualization: C.M.S.

Methodology: L.L., I.R., G.I., H.W., G.D., Y.G., B.R.S., C.M.S.

Formal Analysis: L.L., I.R., G.I., H.W., B.W., C.M.S.

Investigation: L.L., I.R., B.R.S., C.M.S.

Resources: C.M.S., J.C.S., G.P.N., A.T.M., B.W.

Data Curation: L.L., I.R., G.I.

Writing – Original Draft: L.L., C.M.S.

Writing – Review & Editing: All authors

Visualization: L.L., I.R., B.W.

Supervision: C.M.S., B.W.

Project Administration: L.L., C.M.S.

Funding Acquisition: C.M.S., G.P.N., B.W.

## Supplemental Information

Supplemental data includes 14 figures, 7 tables and 2 movies.

