## Supplemental Data for "High-Dimensional Spatiotemporal Single-Cell and 3D Atlas of the Bone Marrow Microenvironment during Leukemic Progression"

---

#### SUPPLEMENTAL FIGURE LEGENDS

**Figure S1: Validation of BM CODEX antibody panel, related to Figure 1 and Table S1A.** Representative CODEX seven-color overlay images comparing the four groups with Hoechst (blue) as a nuclear marker. **(A)** Ki-67 (green), IgM (red), CD5 (white), ESAM (cyan), CD86 (magenta) and osteopontin (yellow). **(B)** CD27 (green), Tie2 (red), CD8a (white), PD-L1 (cyan), Sirpα (magenta) and CD38 (yellow). **(C)** CD117 (green), CD16/32 (red), NKp46 (white), LepR (cyan), CD34 (magenta) and CD61 (yellow). **(D)** Perilipin (green), CD11c (red), CD150 (white), F4/80 (cyan), Ly6C (magenta) and CD48 (yellow). **(E)** CD4 (green), CD47 (red), PD-1 (white), EMCN (cyan), Sca-1 (magenta) and podoplanin (yellow). **(F)** TER-119 (green), CD127 (red), CD135 (white), CD106 (cyan), CD11b (magenta) and CD21/35 (yellow). **(G)** CD45 (green), CD90 (red), ICOS (white), CD105 (cyan), CD169 (magenta) and MHC-II (yellow). **(H)** GFP (green), CD49d (red) and NG2 (yellow). Scale bar, 50 µm.

**Figure S2: Cell lineage assignment, erythroblast subset classification, manual identification of cell types and pericyte distance analysis relative to arterioles and sinuses, related to Figure 2 and Table S3. (A)** Schematic illustrating the strategy for cell lineage assignment. (**B)** Number of segmented single cells per mouse. **(C)** $CD71^{+}TER\text{-}119^{+}$ cells were further classified into EryAs (larger cells) and EryBs (smaller cells) based on cell size. Cells with an area size smaller than 350 pixels were annotated as EryBs, while those equal to or larger than 350 pixels were annotated as EryAs. **(D)** Proportion of EryAs and EryBs within the $CD71^{+}TER119^{+}$ population. **(E)** Schematic representation of erythroblast subtypes based on CD71 and TER-119 expression, as well as cell size, during maturation. **(F, H)** Examples of manually identified MKs ($CD41^{+}$) and adipocytes (perilipin$^{+}$), respectively (see **Methods**). **(G)** Average distances of pericytes to arterioles and sinuses, respectively, with each data point representing the average of distances from one tissue image. Statistics: Student's *t* test. ****p<0.0001.

**Figure S3: Supervised annotation of BM clusters based on marker expression, tissue localization and morphology, related to Figure 2 and Table S3.** After unsupervised clustering, each cluster was overlaid on the CODEX fluorescent images based on X/Y positions and visually verified based on marker expression profiles (for details on each cluster, see **Table S3**), morphology, and tissue localization. Similar clusters were manually merged, resulting in 41 final clusters. Each cluster is depicted as red dots, representing the X/Y coordinates of the cells contained in that cluster. For each cluster, three or five representative markers essential for cluster identification and Hoechst nuclear stain are shown (right panels), as well as an overview of cellular distribution of that cluster within a larger tissue area (red dots on black background, left panels). **(A)** Leukemic cell and PD-L1$^+$ leukemic cell clusters with GFP, GFP-Ab, Ly6G, and PD-L1, respectively. **(B)** LSC cluster with GFP-Ab, CD117 and CD34. **(C)** Granulocyte and monocyte clusters with CD11b, Ly6C and Ly6G. **(D)** PD-L1$^+$ DC cluster with CD11c, MHC-II and PD-L1. **(E)** pDC cluster with B220, CD4, Ly6C, CD11c and CD3. **(F)** MHC-II$^+$ macrophage and MHC-II$^-$ macrophage clusters with F4/80, MHC-II and CD11b. **(G)** APC and PD-L1$^+$ APC clusters with CD16/CD32, MHC-II, F4/80 and PD-L1, respectively. **(H)** CoMP cluster with Ly6C, CD11b and Ly6G. **(I)** eNeP cluster with CD71, CD117, TER-119, CD49d and CD38. **(J)** CD34$^+$CD117$^-$ HSPC, CD34$^+$CD117$^+$ HSPC and CD34$^-$CD117$^+$ HSPC clusters with CD34, CD117, Sca-1 and CD71, respectively. **(K)** EryA, EryB, EryC and ProE clusters with CD71, TER-119 and B220. **(L)** EBI cluster with F4/80, CD169 and CD71. **(M)** Megakaryocyte cluster with CD41, perilipin and B220. **(N)** B cell and plasma cell clusters with B220, IgM and CD71. **(O)** NK cell cluster with NKp46, CD3 and CD45. **(P)** CD4$^+$ T cell, CD8$^+$ T cell and DN T cell clusters with CD3, CD8a and CD4. **(Q)** PD-1$^+$CD4$^+$ T cell and PD-1$^+$CD8$^+$ T cell cluster with CD3, CD4, CD8 and PD-1 respectively. **(R)** Sinus and arteriole clusters with CD31, CD105, Sca-1 and CD71, respectively. **(S)** LepR$^+$ pericyte cluster with LepR, CD31 and B220. **(T)** Bone cell cluster with CD61, osteopontin and podoplanin. **(U)** Adipocyte cluster with perilipin, CD41 and Ly6G. **(V)** Podo$^+$ stromal cell cluster with podoplanin, CD31 and CD61. **(W)** Leukocyte, CD90$^+$ leukocyte and PD-L1$^+$ leukocyte clusters with CD45, B220, Ly6G, CD90, PD-L1 and GFP-Ab, respectively. **(X)** Ki-67+ unidentified cluster with Ki-67, GFP-Ab and CD71.

**Figure S4: Comparison of cell types across the four groups, related to Figure 3. (A)** Composition of the 13 categories in each mouse. **(B-X)** Comparison of individual CTs among the four different groups. Data are presented as mean ± SEM. Statistics: One-way ANOVA (Tukey's post hoc test). No significant differences were observed between the groups.

**Figure S5: Correlation of CT proportions, vessel length, MK subsets and spatial colocalization of Podo$^+$ stromal cells with bone cells, related to Figure 3. (A, D-E, H)** Correlation analysis showing relationships between CT proportions in groups. (A)

Leukemic cells and vascular cells, (D) MKs and vascular cells, (E) leukemic cells and MKs, (H) leukemic cells and bone cells. Each data point represents the frequency of CT from a tissue region within the corresponding group. **(B-C)** Quantification of CD31-stained vessel length across groups. (B) Total vessel length (mm), (C) Average vessel length (mm). Each data point represents a tissue region. **(F-G)** Quantification of (F) $CD150^+$ MKs and (G) $CD150^+CD34^+$ MKs as a proportion of total MKs across different groups. **(I)** Representative CODEX images showing colocalization of $Podo^+$ stromal cells and $CD61^+$/$osteopontin^+$ bone cells. Scale bar, 50 μm. Data are presented as mean ± SEM. Statistics: (A, D-E, H) Linear regression analysis (Pearson's correlation coefficient); (B, F-G) One-way ANOVA (Tukey's post hoc test). ***$p<0.001$, ****$p<0.0001$.

**Figure S6: Visualization of blood vessels in CODEX regions across groups, related to Figure 3.** CODEX images highlighting blood vessels identified by CD31 staining. **(A)** WT, **(B)** d7 CML, **(C)** d14 CML and **(D)** d21 CML mice BM. Scale bar, 100 μm.

**Figure S7: CT composition within each CN, related to Figure 4. (A)** Bubble plot of cellular neighborhoods. The size of the circles represents the frequency of a given CT within the respective CN (horizontally). The color of the circles indicates the frequency of a given CT in a particular CN compared to the total number of cells of that CT in all CNs (vertically). **(B)** Absolute cell numbers of the 41 CTs in each CN.

**Figure S8: CN Voronoi maps comparing BM from d7 and d14 CML mice, related to Figure 4.** Representative CODEX images (left column), Voronoi diagrams of CNs (middle column) and corresponding H&E images (right column) in d7 CML (upper row) and d14 CML (lower row) mice. Scale bar, 50 μm.

**Figure S9: CT distribution across CNs, related to Figure 4.** Numbers of cells of each CT within each of the 14 CNs.

**Figure S10: Voronoi diagrams and distribution of CNs, related to Figure 4.** Voronoi diagrams illustrate the spatial distribution of the 14 CNs in the BM of **(A)** WT , **(B)** d7 CML, **(C)** d14 CML and **(D)** d21 CML mice BM. Scale bar, 100 µm.

**Figure S11: T cell frequency and neighborhood enrichment analysis, related to Figure 5. (A)** Comparison of the T cell frequencies (% of all nucleated cells) across the different groups. **(B)** Heatmap of estimated differential enrichment coefficients for T cell subsets in WT vs. advanced CML mice. Data are presented as mean ± SEM. Statistics: Regression analysis, not adjusted for multiple tests. *p<0.05.

**Figure S12: MHC-II and Ki-67 expression in HSPC subsets and neighborhood enrichment analysis, related to Figure 6. (A-C)** Comparison of MHC-II mean fluorescent intensities (MFI) in (A) $CD34^{+}CD117^{-}$ HSPCs, (B) $CD34^{+}CD117^{+}$ HSPCs, and (C) $CD34^{-}CD117^{+}$ HSPCs across the four groups. **(D)** MFI of MHC-II and Ki-67 in all HSPCs, analyzed collectively, with each data point representing an individual HSPC. **(E)** Heatmap of estimated differential enrichment coefficients for HSPCs subsets in WT vs. advanced CML mice. Statistics: (A-C) Student's *t* test (WT vs. d21 CML); (E) Regression analysis, not adjusted for multiple tests. *p<0.05, ****p<0.0001.

**Figure S13: Representative CODEX images of pDC–MK interactions and granulocyte emperipolesis by MKs, related to Figure 7. (A)** Close contacts between pDCs and MKs (white arrows indicate pDCs). **(B)** Granulocytes internalized by MKs. Scale bar, 10 µm.

**Figure S14: MK isolation by laser capture microdissection, and morphological changes in MKs, related to Figure 7. (A)** Step-by-step procedure for isolating human MKs from FFPE BM trephine biopsy sections using laser capture microdissection (LCM). Step 1, identifying a target MK under the microscope; Step 2, marking the MK for laser dissection (red outline); Step 3, precise laser cutting around the MK border; Step 4, successful laser capture of the isolated MK. **(B)** Isolated single MKs collected on the LCM cap. **(C-D)** H&E staining of BM sections from patients with TKI-relapsed and newly diagnosed apCML and bpCML patients, respectively. Green arrows indicate MKs. Scale bar, 50 µm.

#### SUPPLEMENTAL TABLES

**Table S1:** Antibodies, clones, manufacturers and corresponding CODEX oligonucleotides.

**Table S2:** Categorized cell types and surface markers.

**Table S3:** Patient cohort characteristics.

**Table S4:** Sequences of CODEX oligonucleotides.

**Table S5:** Composition of antibody mixes.

**Table S6:** CODEX multi-cycle panels.

**Table S7:** cDNA library preparation solutions.

#### SUPPLEMENTAL MOVIES

**Movie S1: 3D view of a whole WT femur, related to Figure 7.** Bone cortex is shown in gray, endomucin + CD31 in red, and CD42b in yellow.

**Movie S2: 3D view of a whole d21 CML femur, related Figure 7.** Bone cortex is shown in gray, endomucin + CD31 in red, and CD42b in yellow.

Figure S1A-B

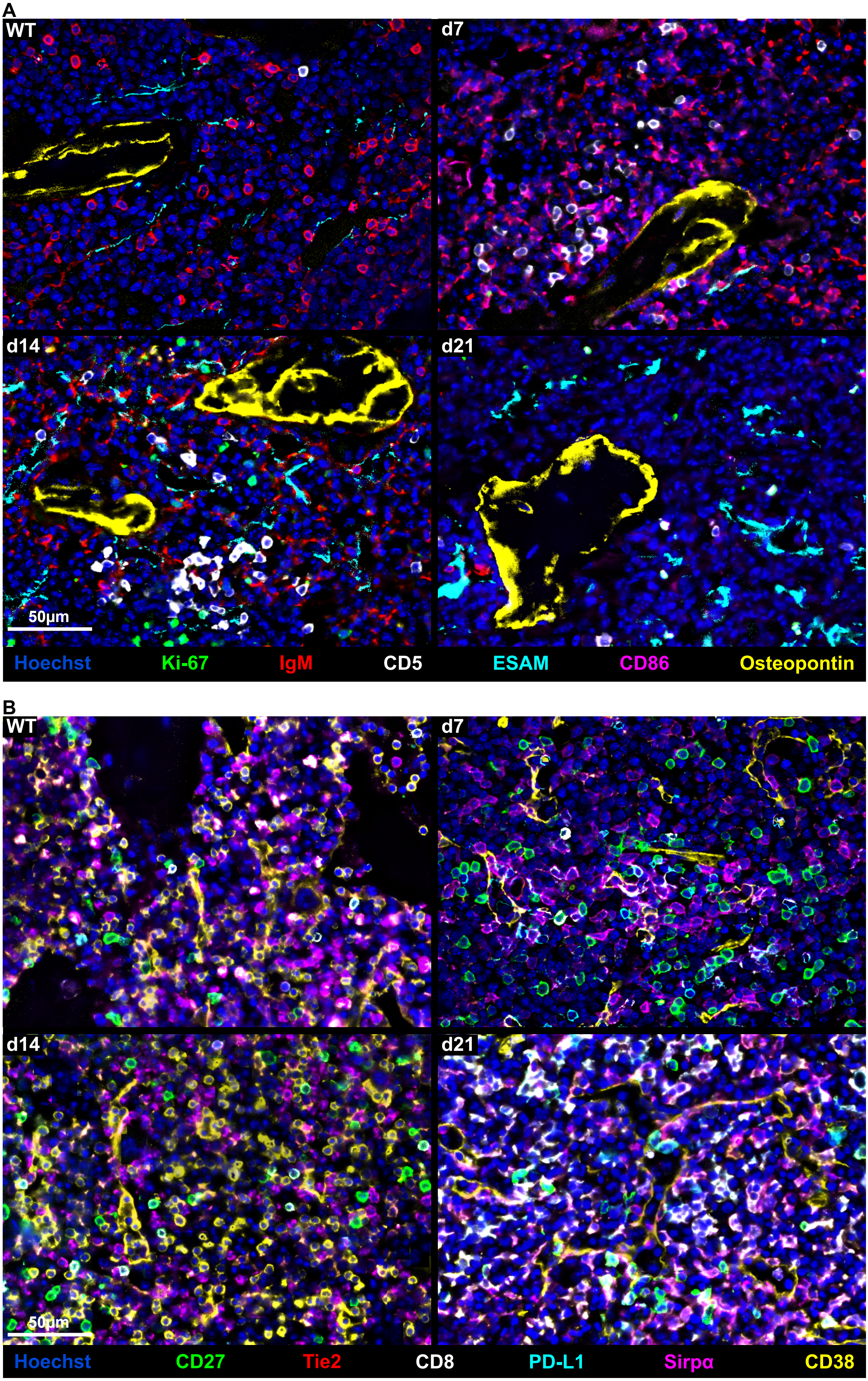

A
WT
d7
d14
d21
50µm
Hoechst
Ki-67
IgM
CD5
ESAM
CD86
Osteopontin
B
WT
d7
d14
d21
50µm
Hoechst
CD27
Tie2
CD8
PD-L1
Sirpα
CD38

**Figure S1C-D**

C

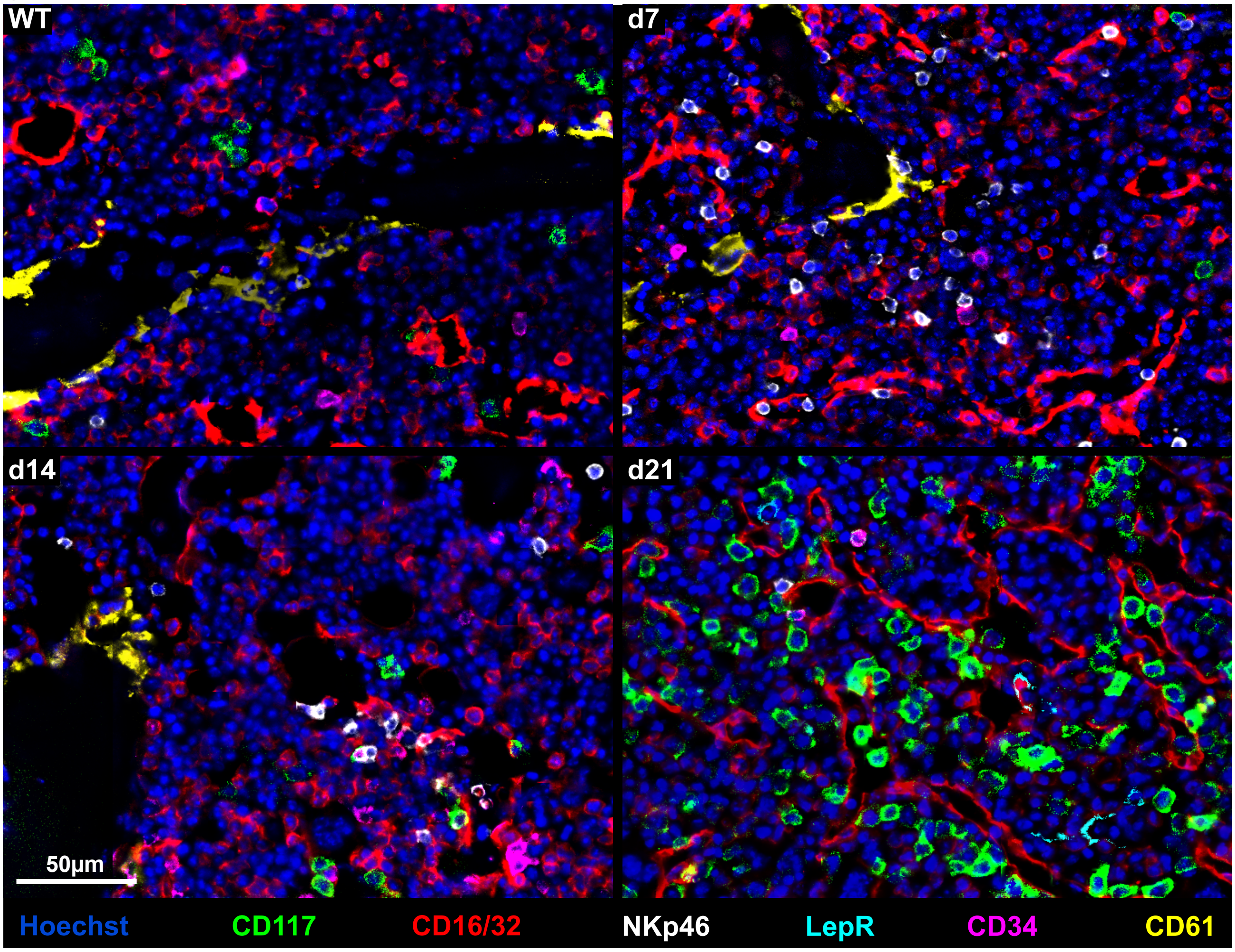


D

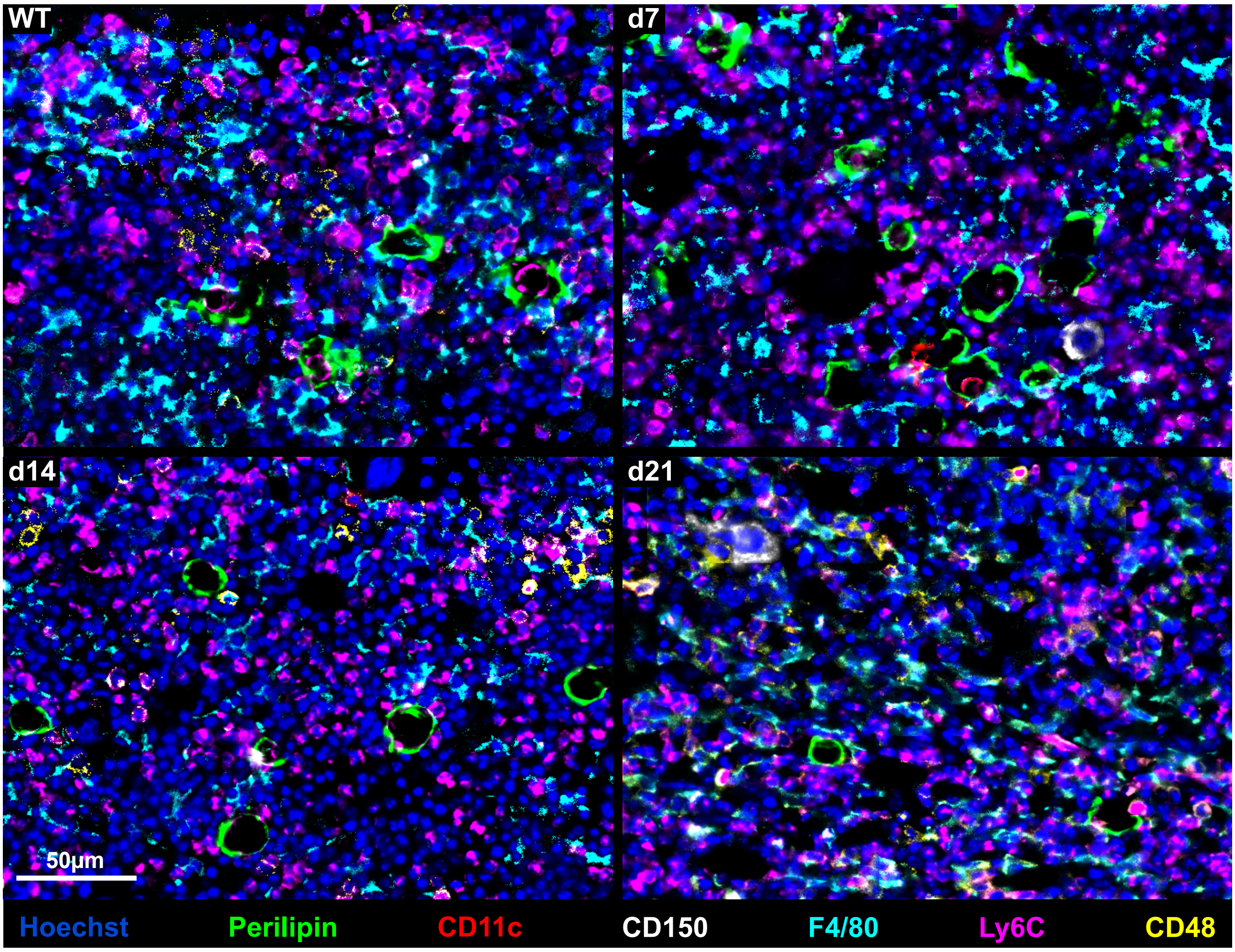

### Figure S1E-F

E

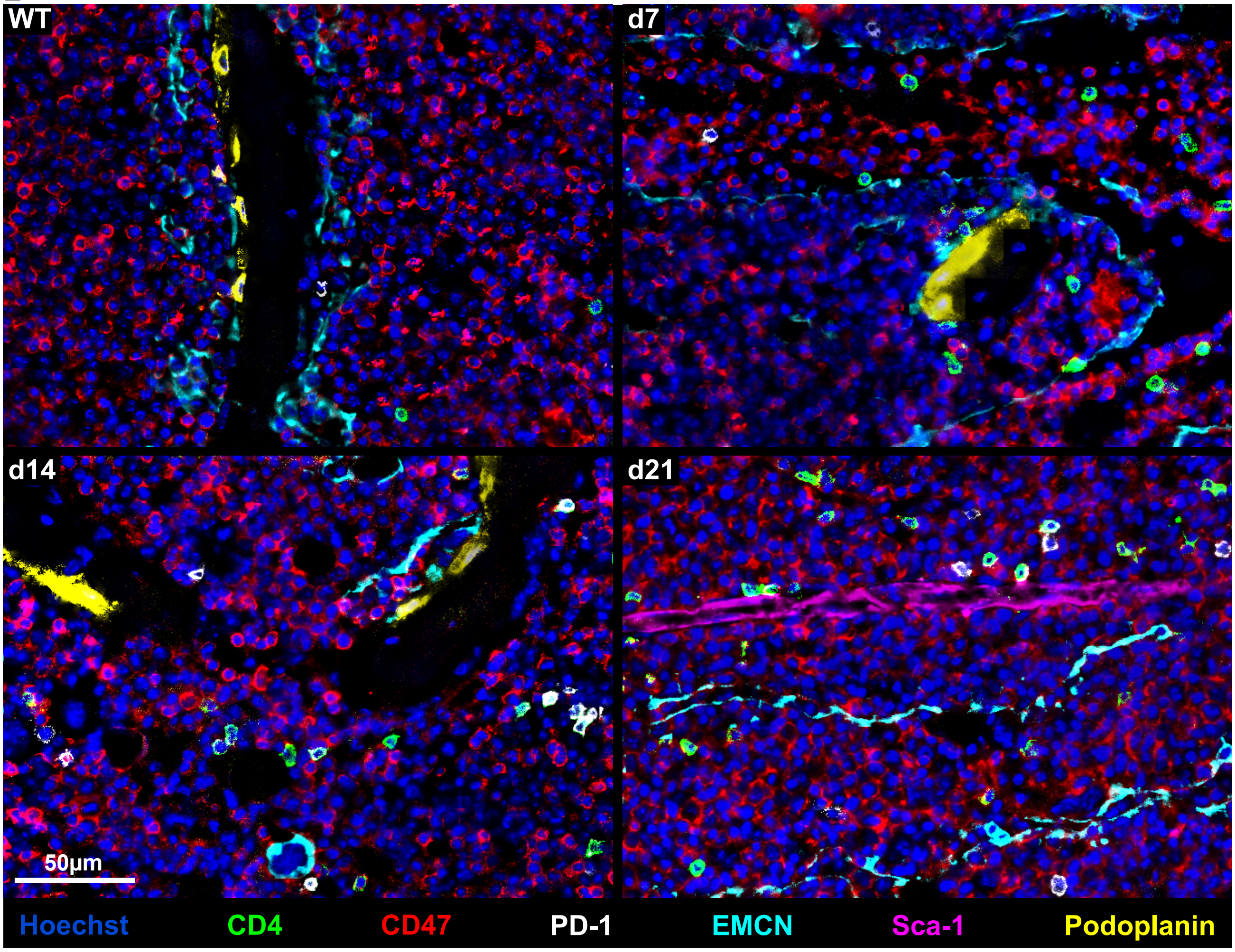


F

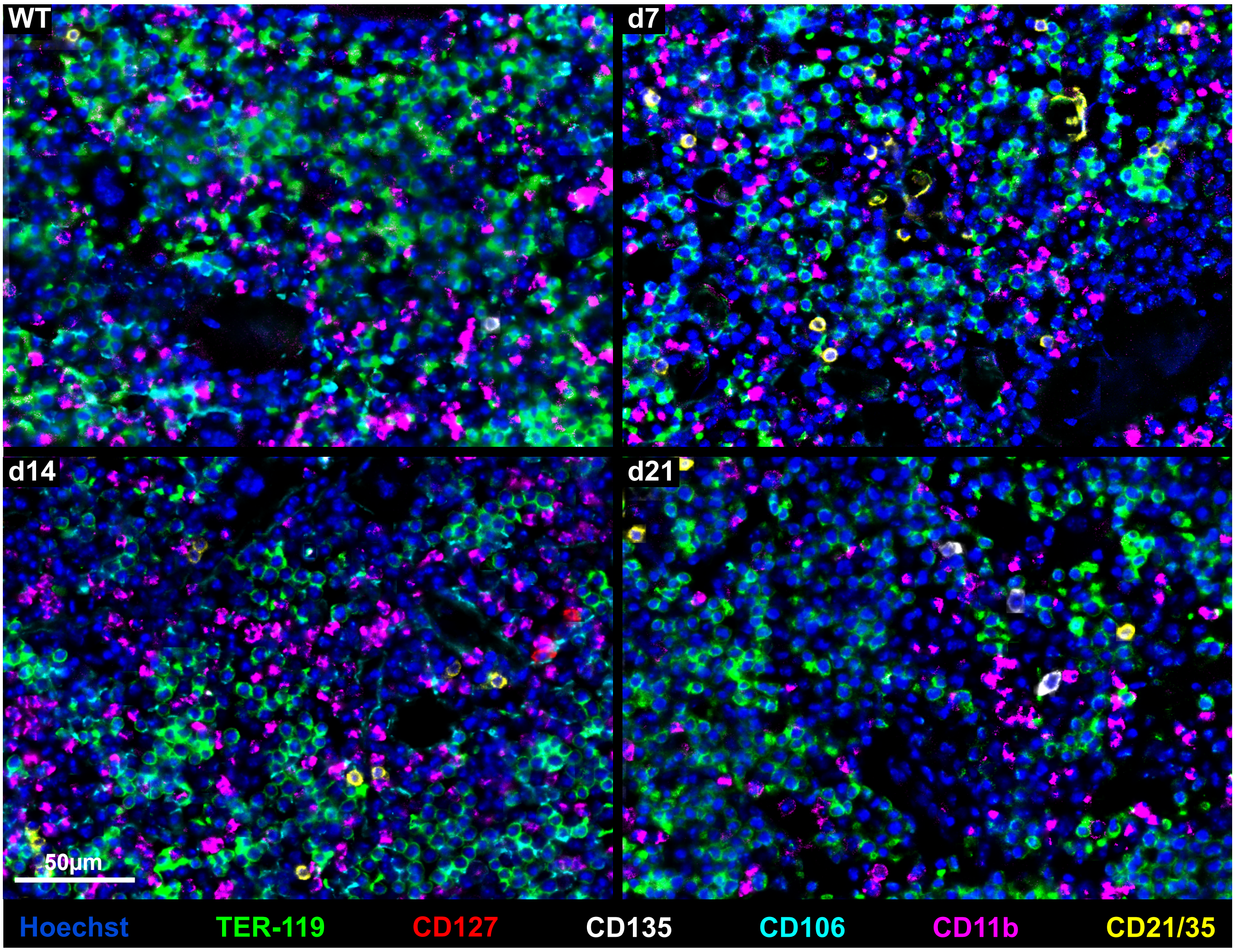

Figure S1G-H

G

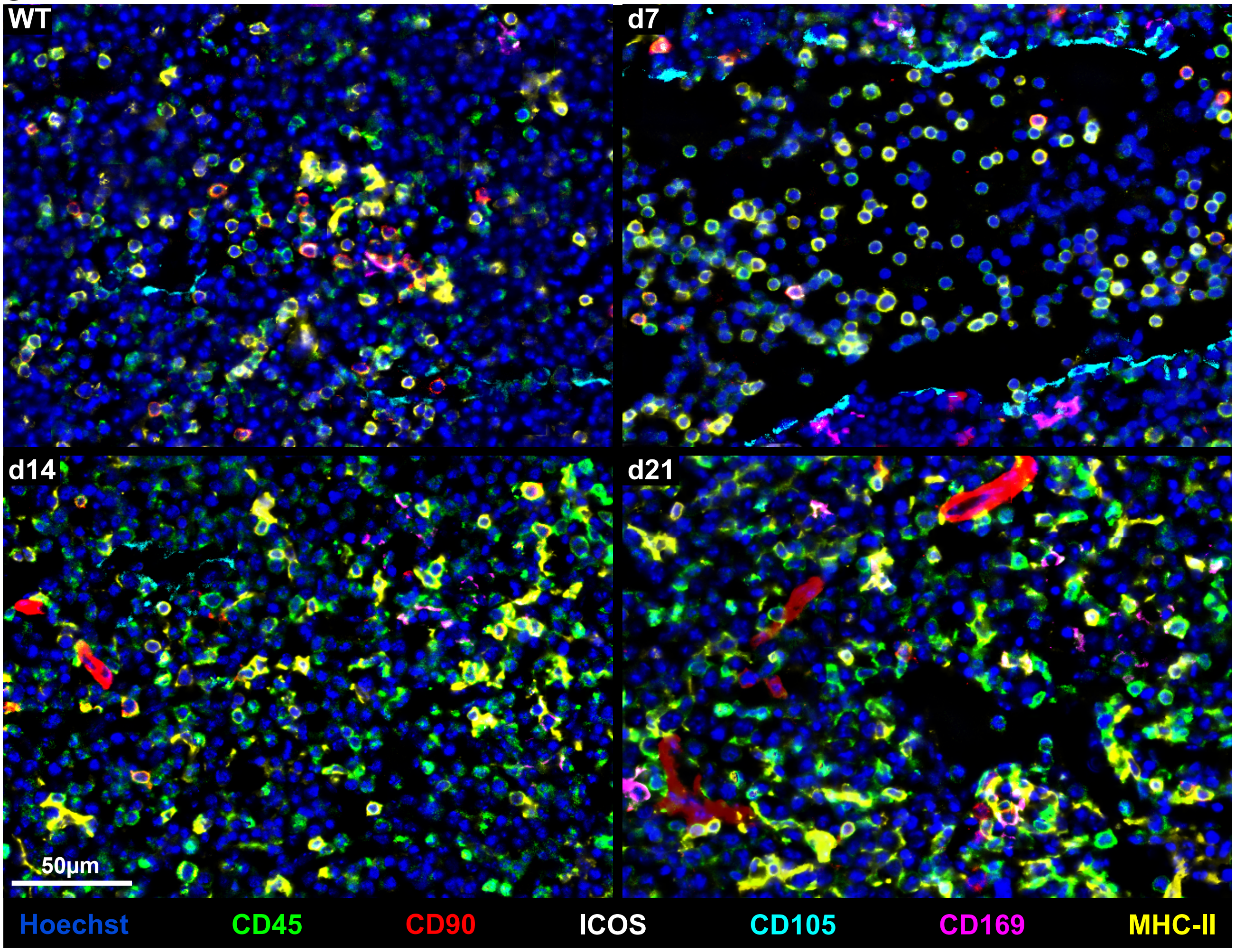


H

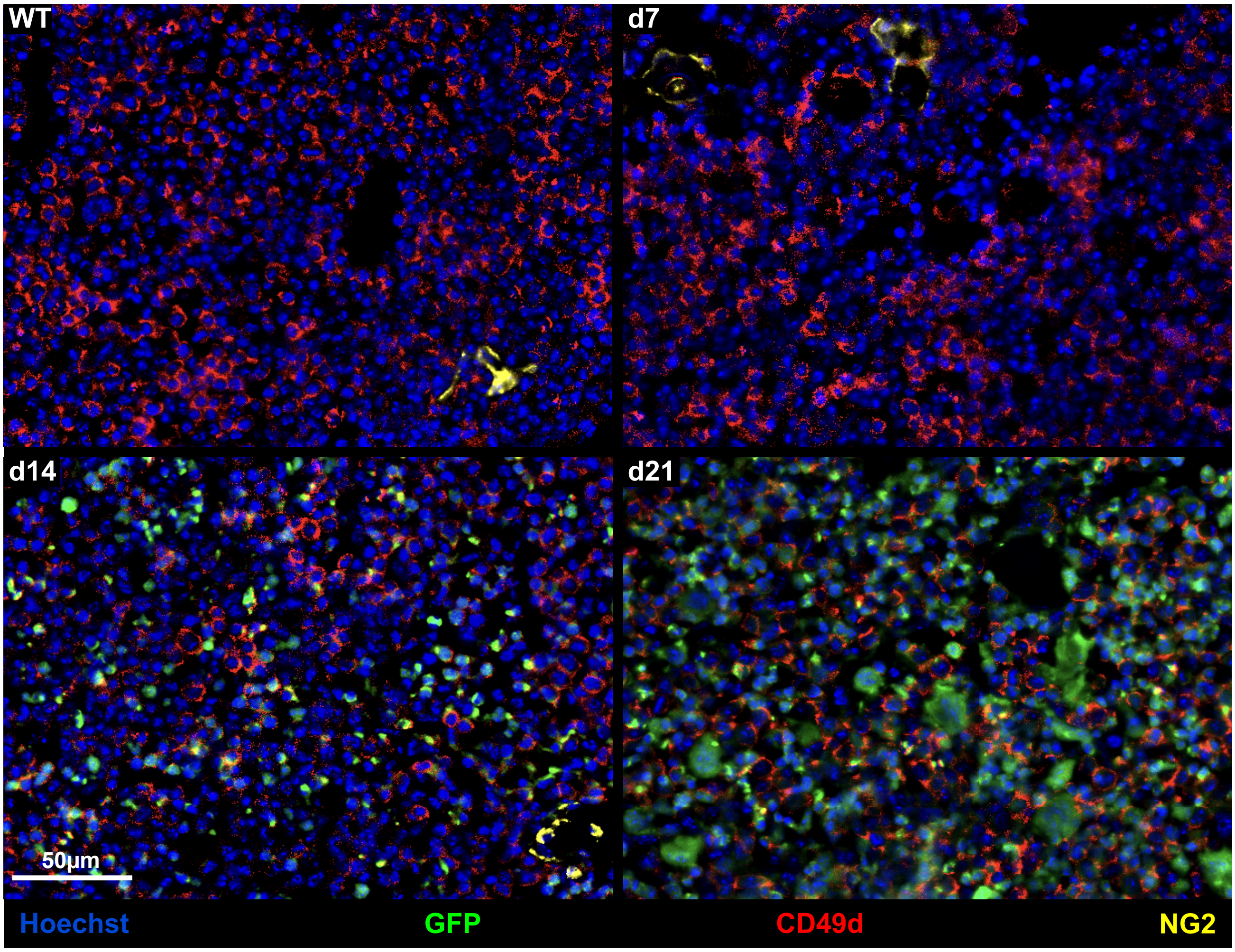

### Figure S2

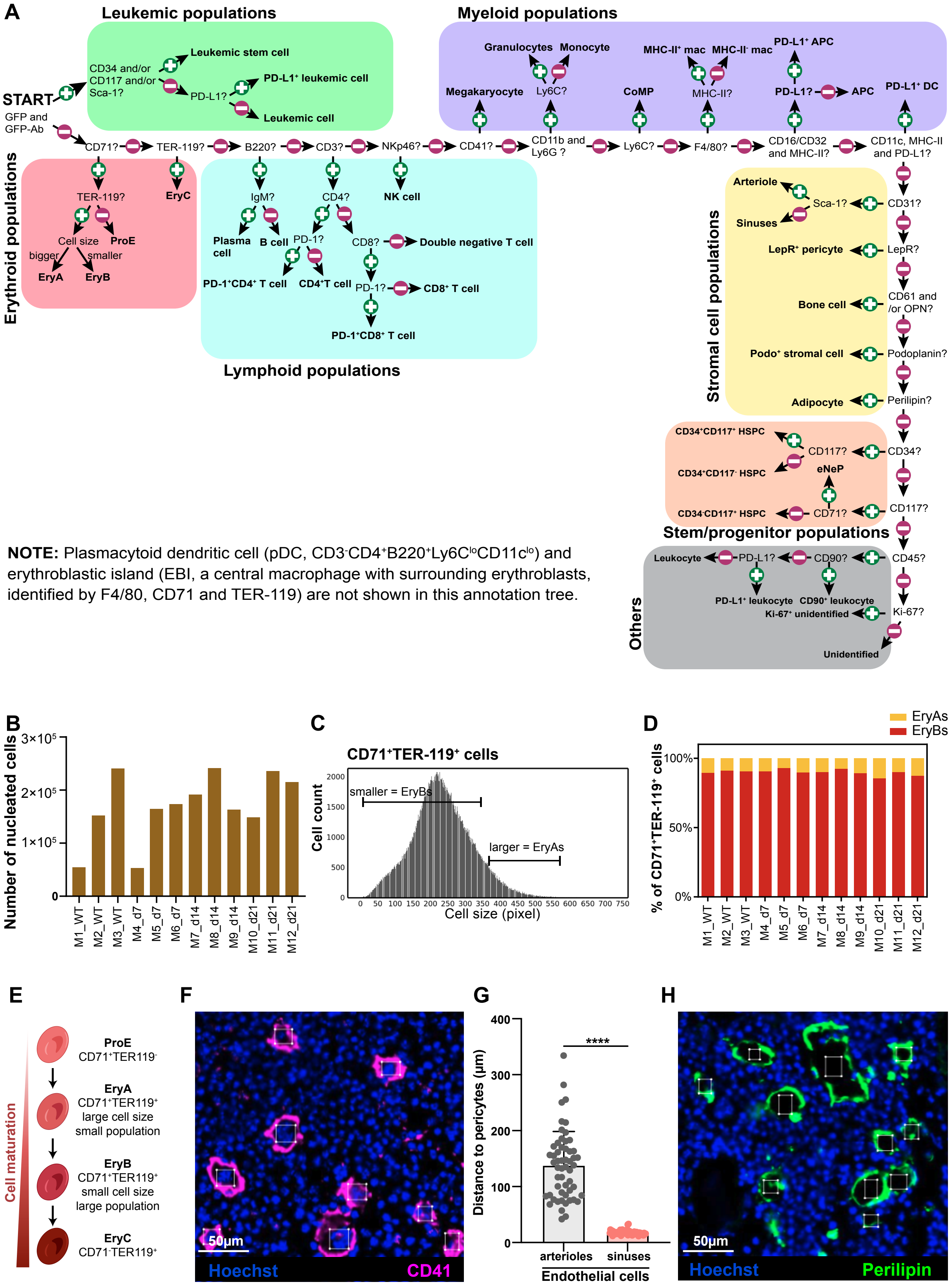
A
Leukemic populations
Myeloid populations
START
GFP and GFP-Ab
CD34 and/or CD117 and/or Sca-1?
Leukemic stem cell
PD-L1?
PD-L1⁺ leukemic cell
Leukemic cell
Granulocytes
Monocyte
Ly6C?
Megakaryocyte
CoMP
MHC-II⁺ mac
MHC-II⁻ mac
MHC-II?
PD-L1⁺ APC
PD-L1?
APC
PD-L1⁺ DC
CD71?
TER-119?
B220?
CD3?
NKp46?
CD41?
CD11b and Ly6G ?
Ly6C?
F4/80?
CD16/CD32 and MHC-II?
CD11c, MHC-II and PD-L1?
Erythroid populations
TER-119?
EryC
Cell size
ProE
bigger
smaller
EryA
EryB
IgM?
CD4?
NK cell
Plasma cell
B cell
PD-1?
CD8?
Double negative T cell
PD-1⁺CD4⁺ T cell
CD4⁺T cell
PD-1?
CD8⁺ T cell
PD-1⁺CD8⁺ T cell
Lymphoid populations
Stromal cell populations
Arteriole
Sca-1?
CD31?
Sinuses
LepR⁺ pericyte
LepR?
Bone cell
CD61 and /or OPN?
Podo⁺ stromal cell
Podoplanin?
Adipocyte
Perilipin?
CD34⁺CD117⁺ HSPC
CD117?
CD34?
CD34⁺CD117⁻ HSPC
eNeP
CD34⁻CD117⁺ HSPC
CD71?
CD117?
Stem/progenitor populations
NOTE: Plasmacytoid dendritic cell (pDC, CD3⁻CD4⁺B220⁺Ly6C^lo CD11c^lo) and erythroblastic island (EBI, a central macrophage with surrounding erythroblasts, identified by F4/80, CD71 and TER-119) are not shown in this annotation tree.
Leukocyte
PD-L1?
CD90?
CD45?
PD-L1⁺ leukocyte
CD90⁺ leukocyte
Ki-67⁺ unidentified
Ki-67?
Others
Unidentified
B
Number of nucleated cells
3×10⁵
2×10⁵
1×10⁵
0
M1_WT
M2_WT
M3_WT
M4_d7
M5_d7
M6_d7
M7_d14
M8_d14
M9_d14
M10_d21
M11_d21
M12_d21
C
CD71⁺TER-119⁺ cells
Cell count
smaller = EryBs
larger = EryAs
Cell size (pixel)
D
EryAs
EryBs
% of CD71⁺TER-119⁺ cells
100%
50%
0%
E
ProE
CD71⁺TER119⁻
EryA
CD71⁺TER119⁺
large cell size
small population
EryB
CD71⁺TER119⁺
small cell size
large population
EryC
CD71⁻TER119⁺
Cell maturation
F
50µm
Hoechst
CD41
G
Distance to pericytes (µm)
****
arterioles
sinuses
Endothelial cells
H
50µm
Hoechst
Perilipin

### Figure S3A

**A** Leukemic cell cluster (d7, GFP$^+$GFP-Ab$^+$)

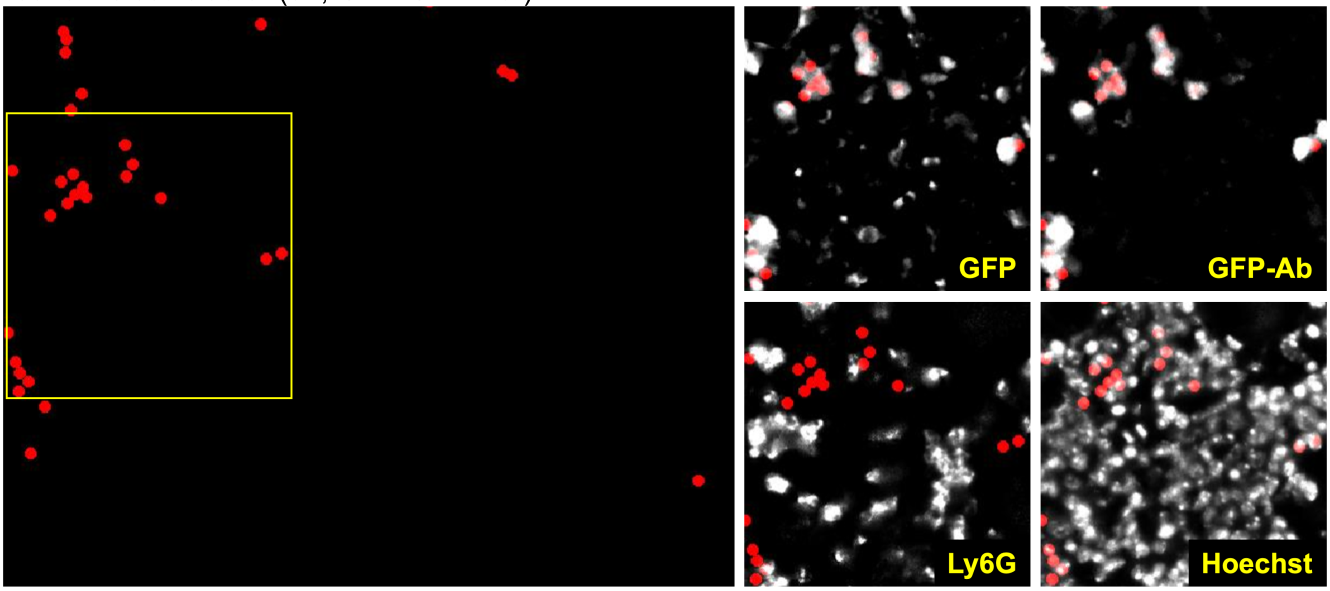


Leukemic cell cluster (d14, GFP$^+$GFP-Ab$^+$)

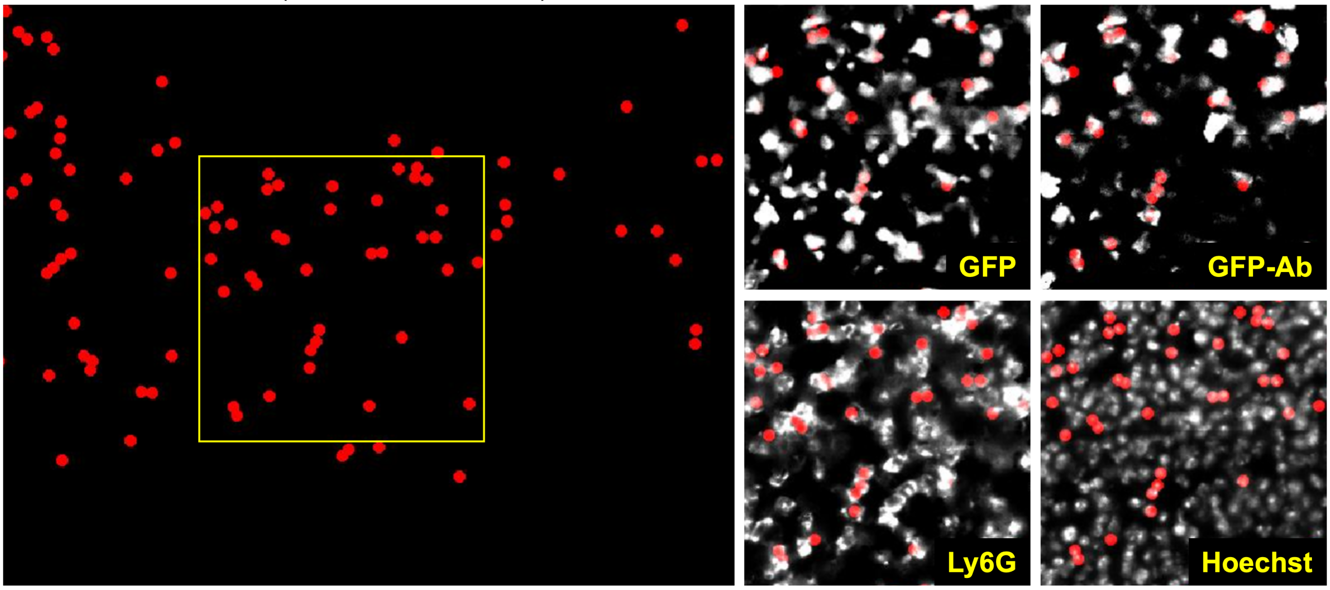


Leukemic cell cluster (d21, GFP$^+$GFP-Ab$^+$)

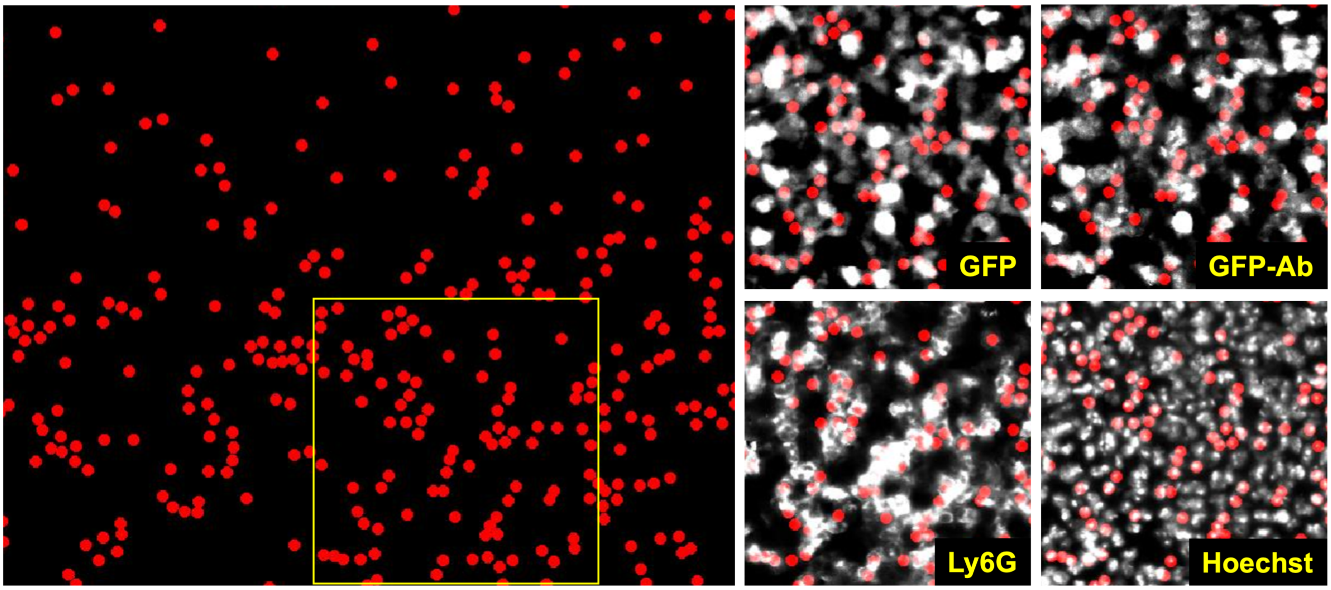

### Figure S3B

PD-L1+ leukemic cell cluster (PD-L1+GFP+GFP-Ab+)

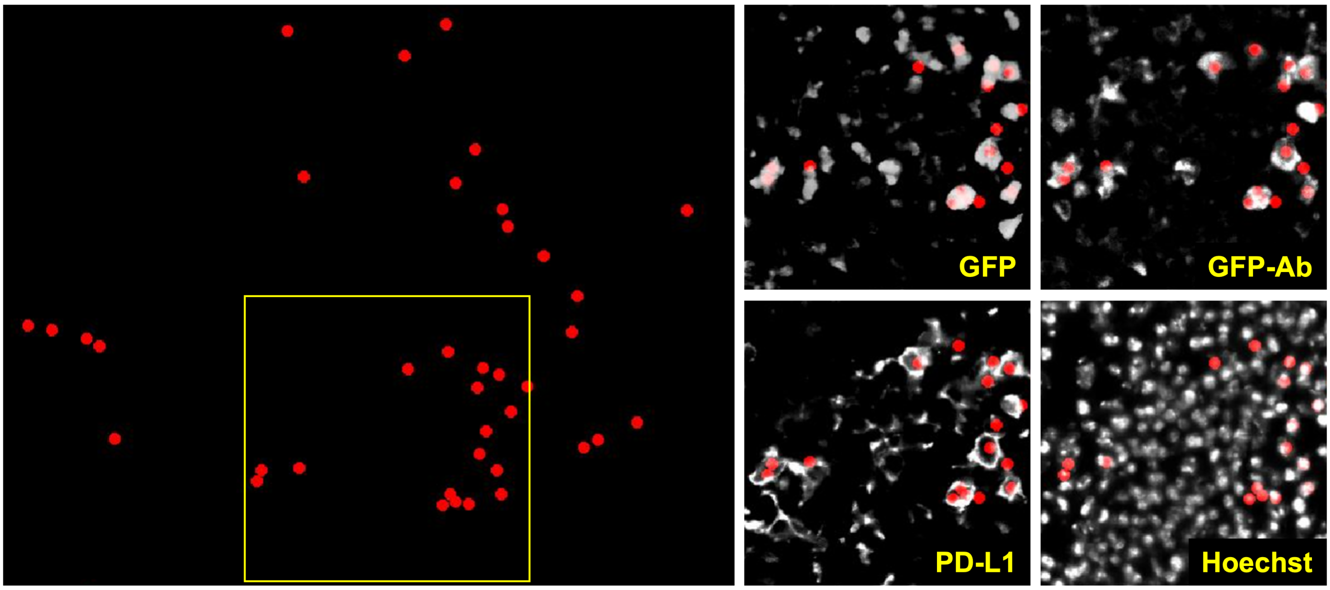


**B** LSCs cluster (d14, lin$^-$CD117+/CD34+GFP+GFP-Ab+)

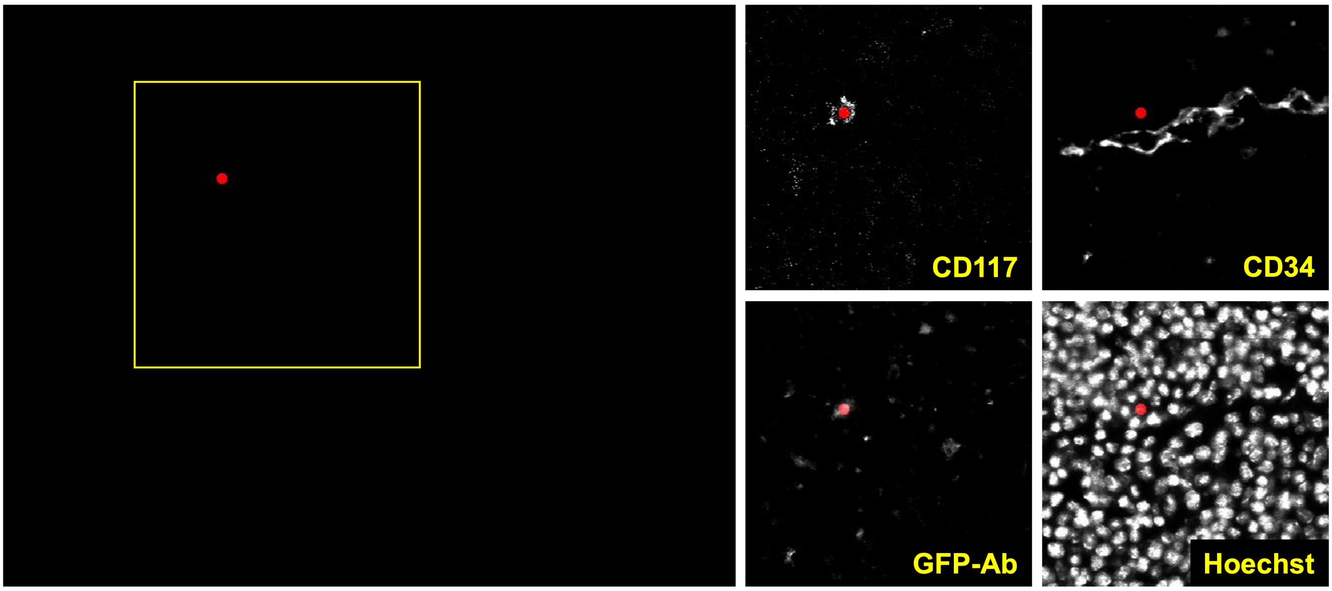


LSCs cluster (d21, lin$^-$CD117+/CD34+GFP+GFP-Ab+)

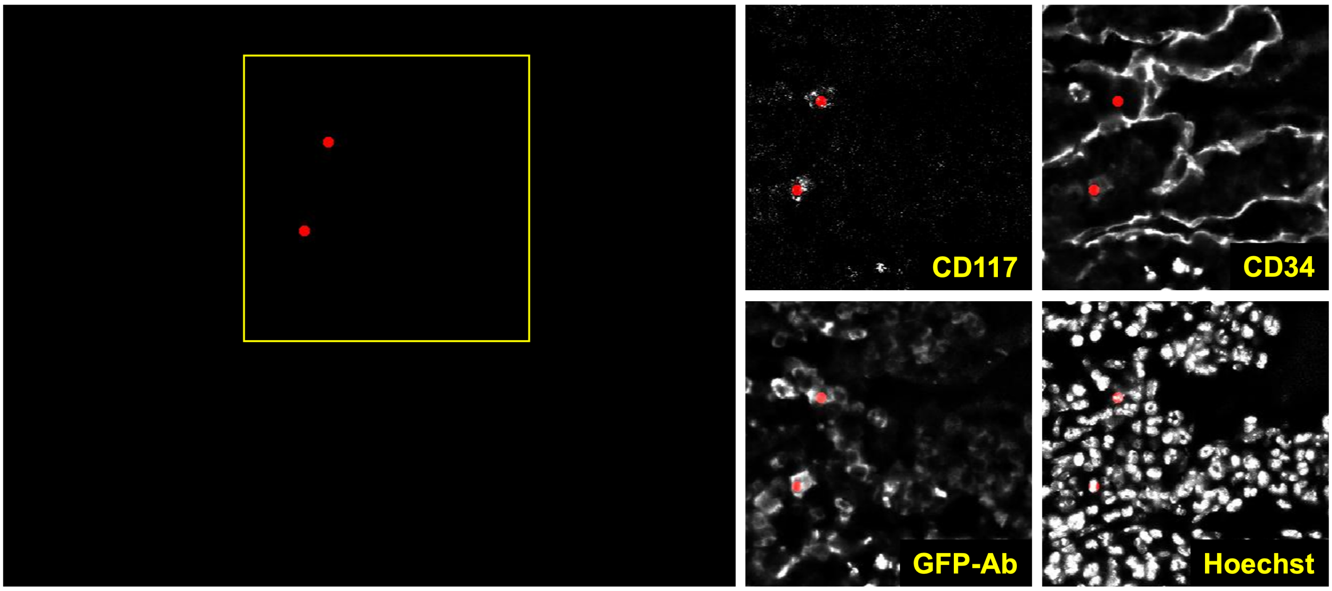

#### Figure S3C-D

**C** Granulocyte cluster (CD11b⁺Ly6C⁺Ly6G⁺)

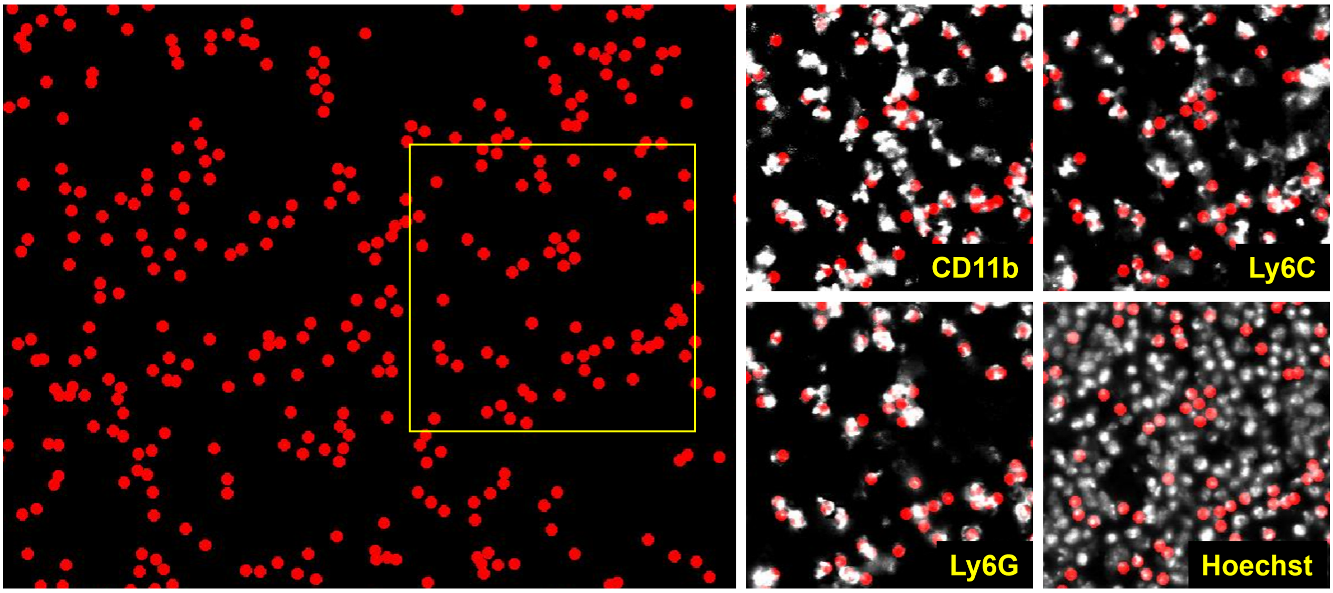


Monocyte cluster (CD11b⁺Ly6C⁺Ly6G⁻)

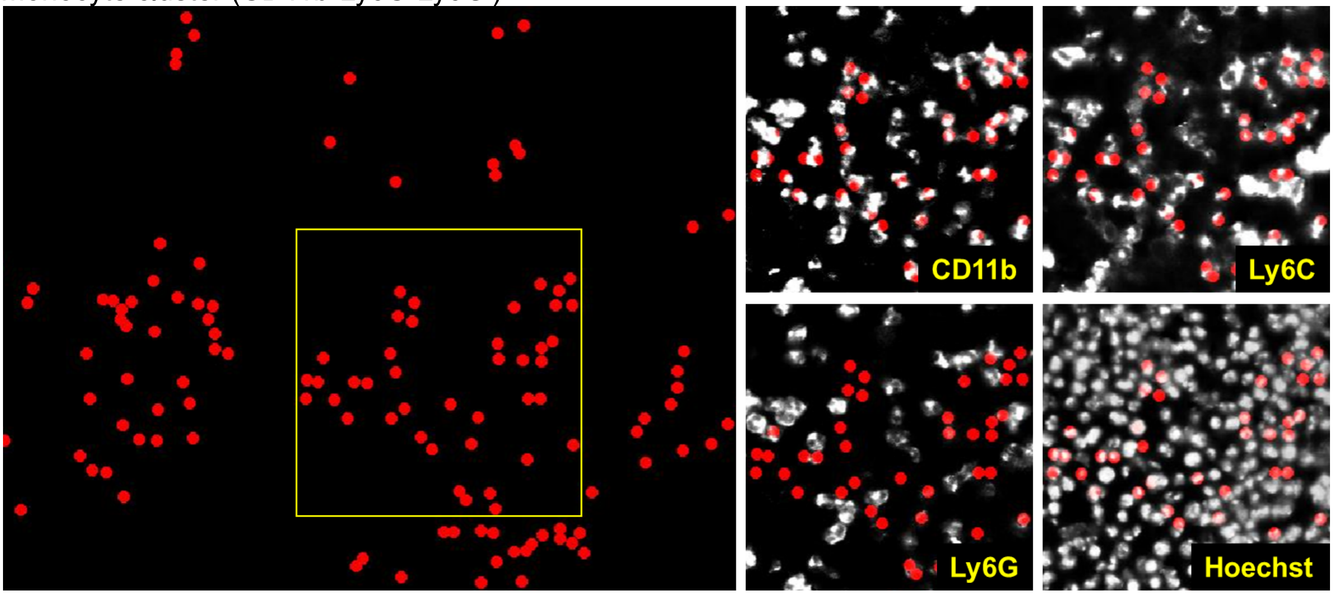


**D** PD-L1⁺ DC cluster (PD-L1⁺CD11c⁺MHC-II⁺)

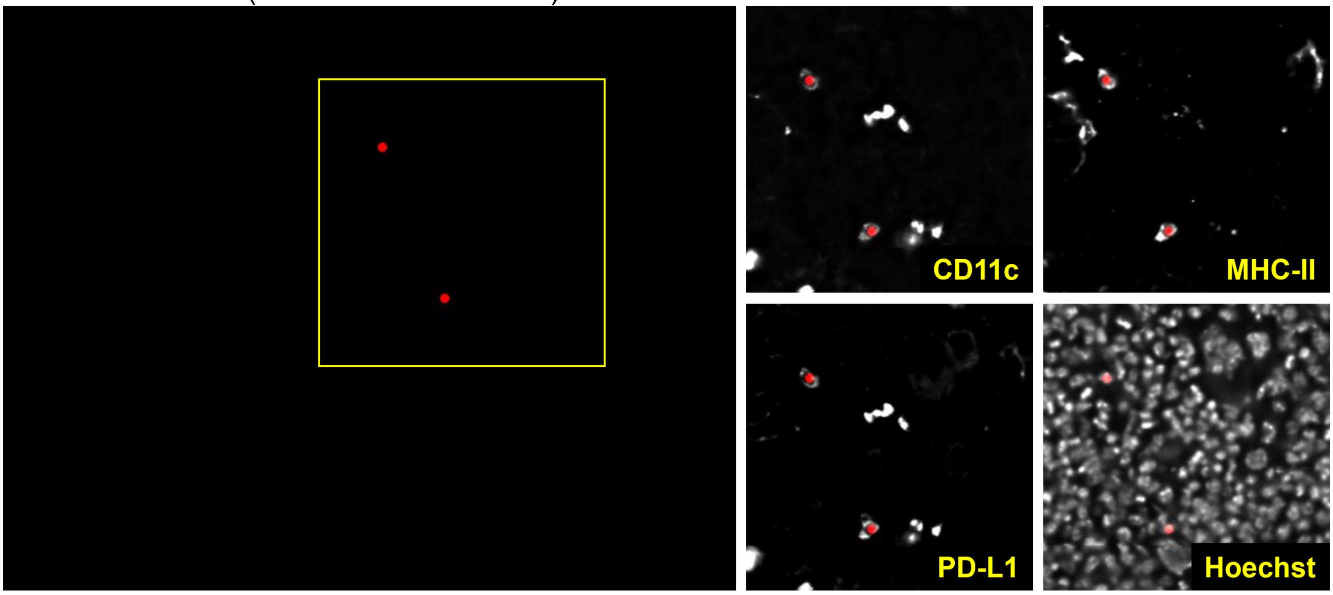

### Figure S3E-F

**E** pDC cluster (CD3$^{-}$CD4$^{+}$B220$^{+}$Ly6C$^{lo}$CD11c$^{lo}$)

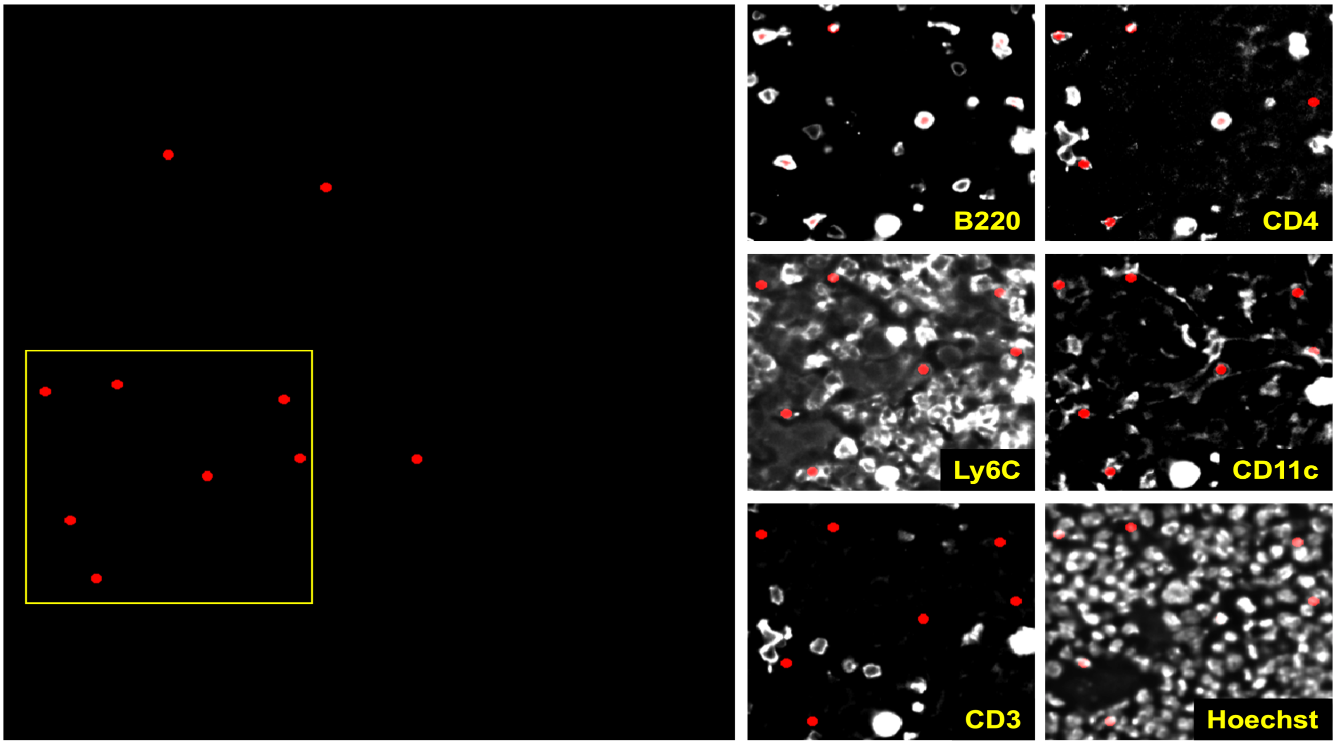


**F** MHC-II$^{+}$ macrophage cluster (MHC-II$^{+}$F4/80$^{+}$)

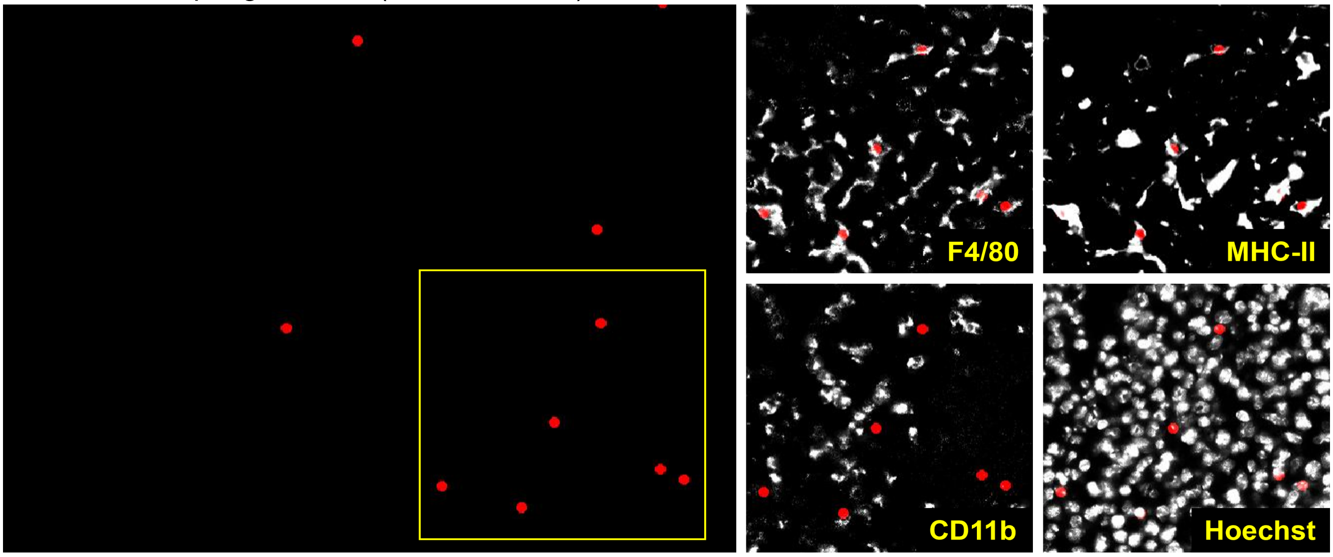


MHC-II$^{-}$ macrophage cluster (MHC-II$^{-}$F4/80$^{+}$)

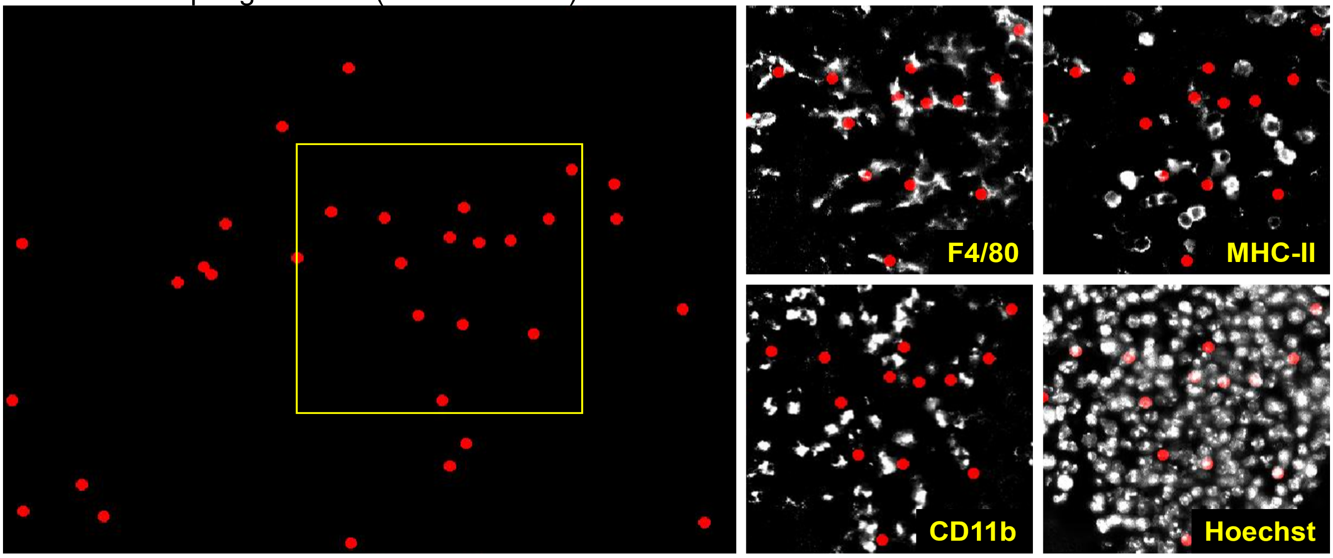

### Figure S3G-H

**G** APC cluster (CD16/32$^{+}$MHC-II$^{+}$)

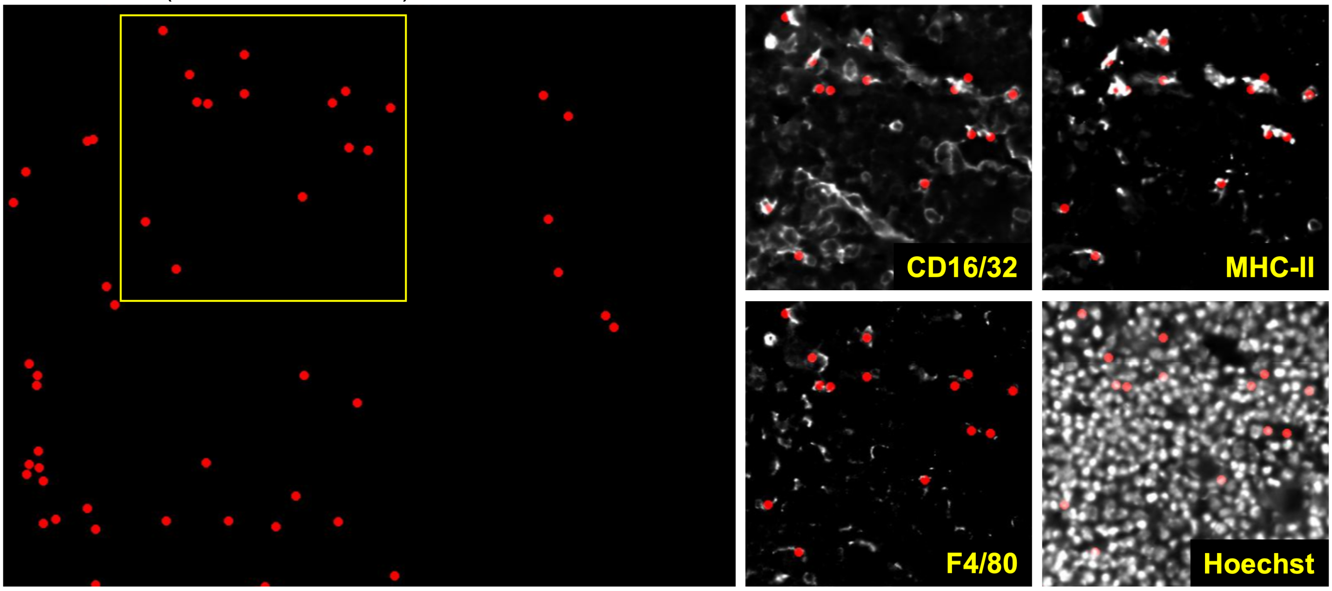


PD-L1$^{+}$ APC cluster (PD-L1$^{+}$CD16/32$^{+}$MHC-II$^{+}$)

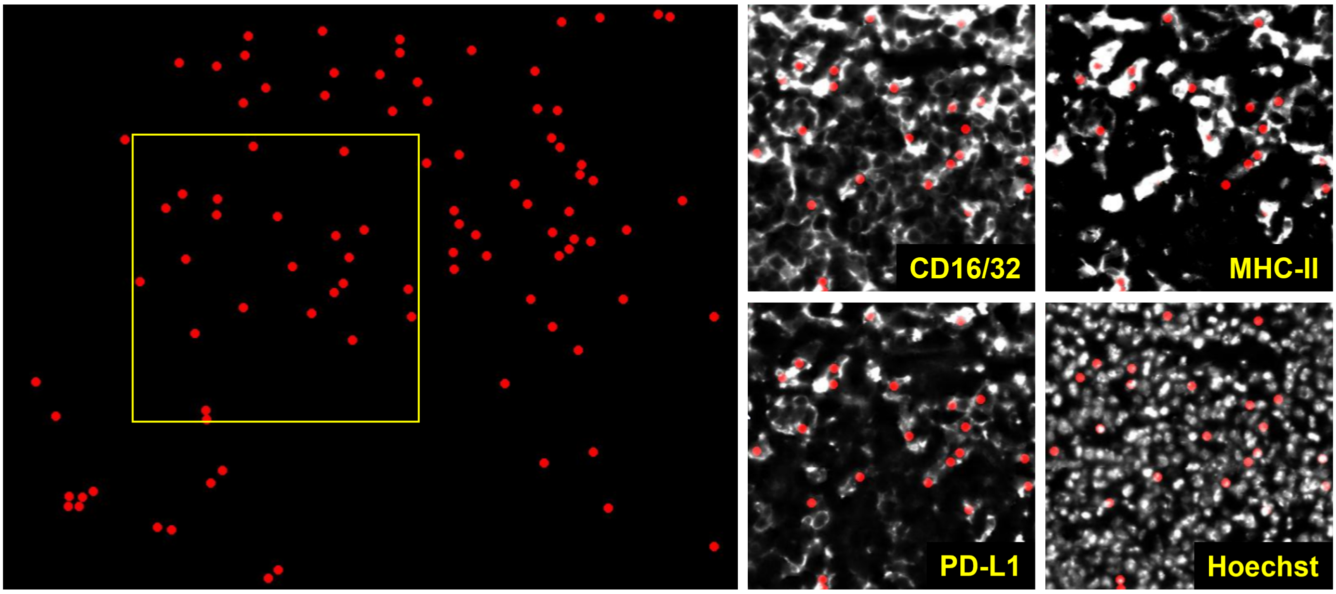


**H** CoMP cluster (lin$^{-}$CD11b$^{-}$Ly6C$^{+}$Ly6G$^{-}$)

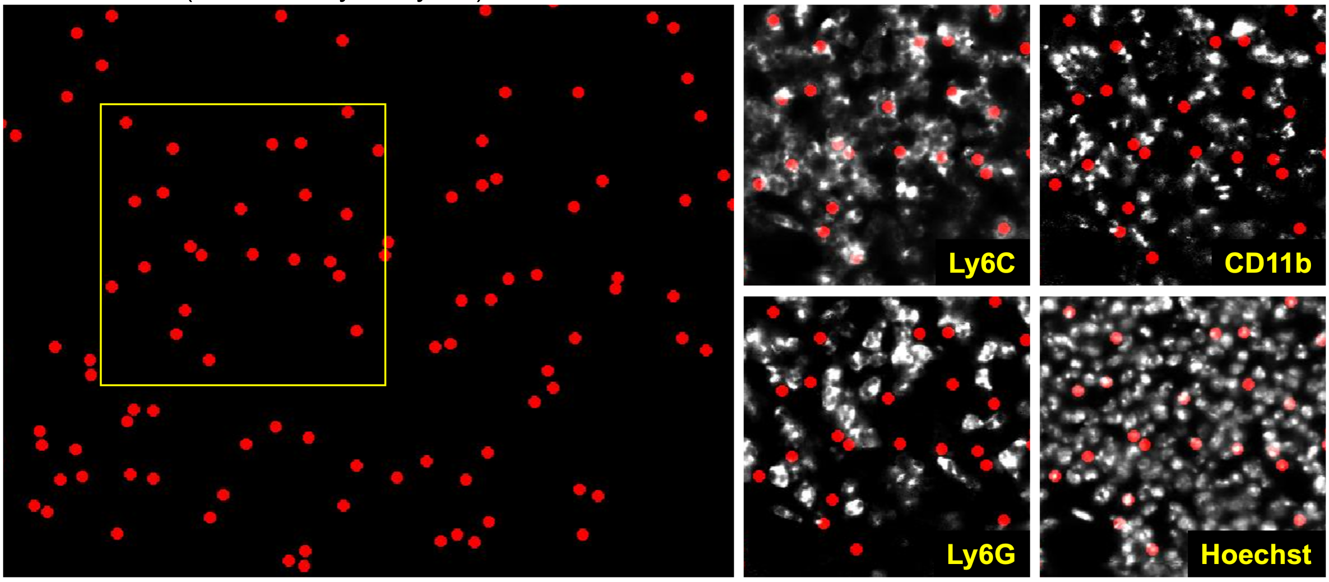

### Figure S3I-J

**I** eNeP cluster (lin$^-$CD117$^+$CD71$^+$CD49d$^+$CD38$^-$)

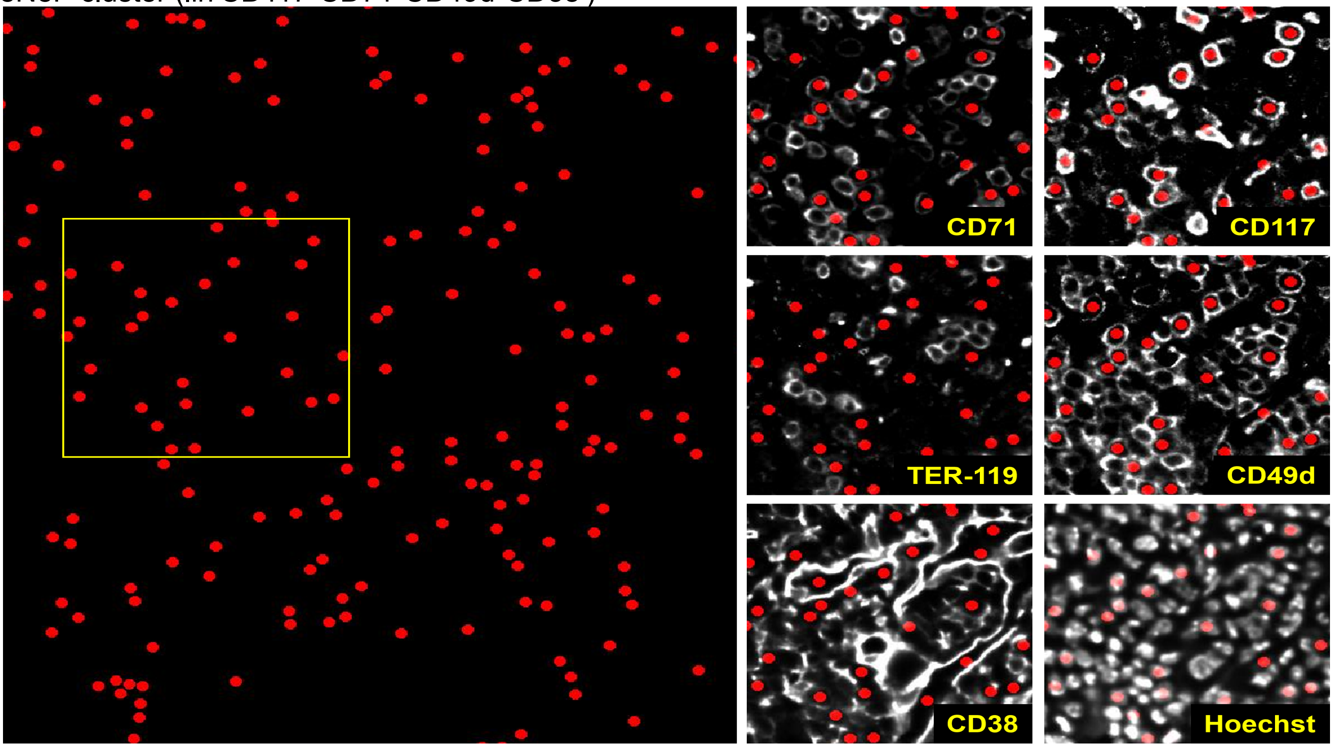


**J** CD34$^+$CD117$^-$ HSPC cluster (lin$^-$CD34$^+$CD117$^-$)

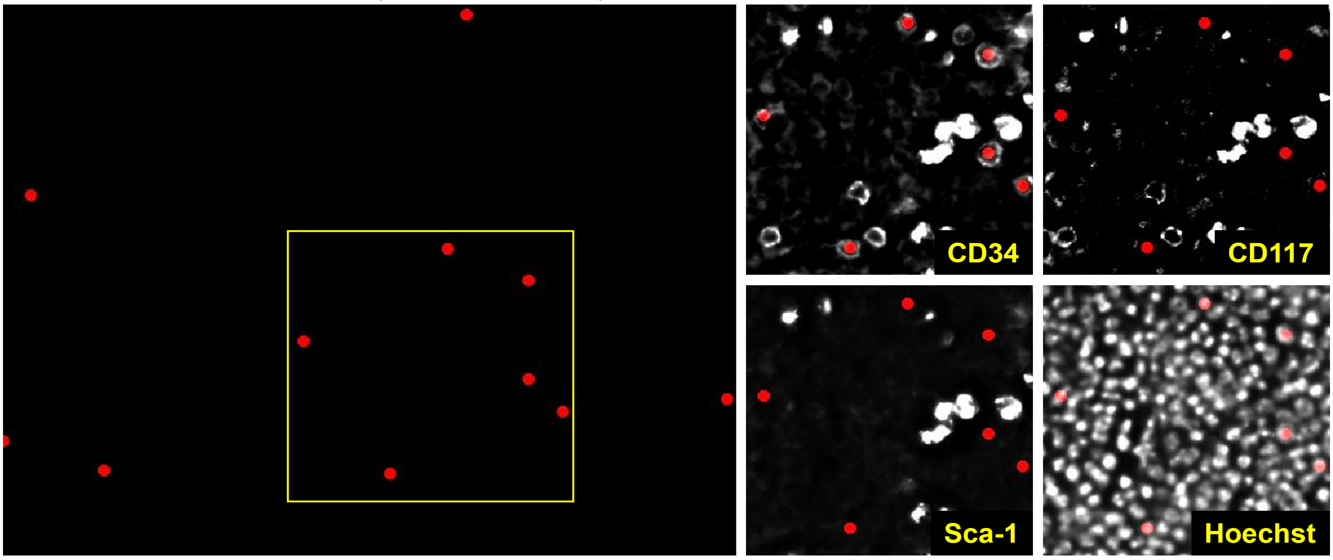


CD34$^+$CD117$^+$ HSPC cluster (lin$^-$CD34$^+$CD117$^+$)

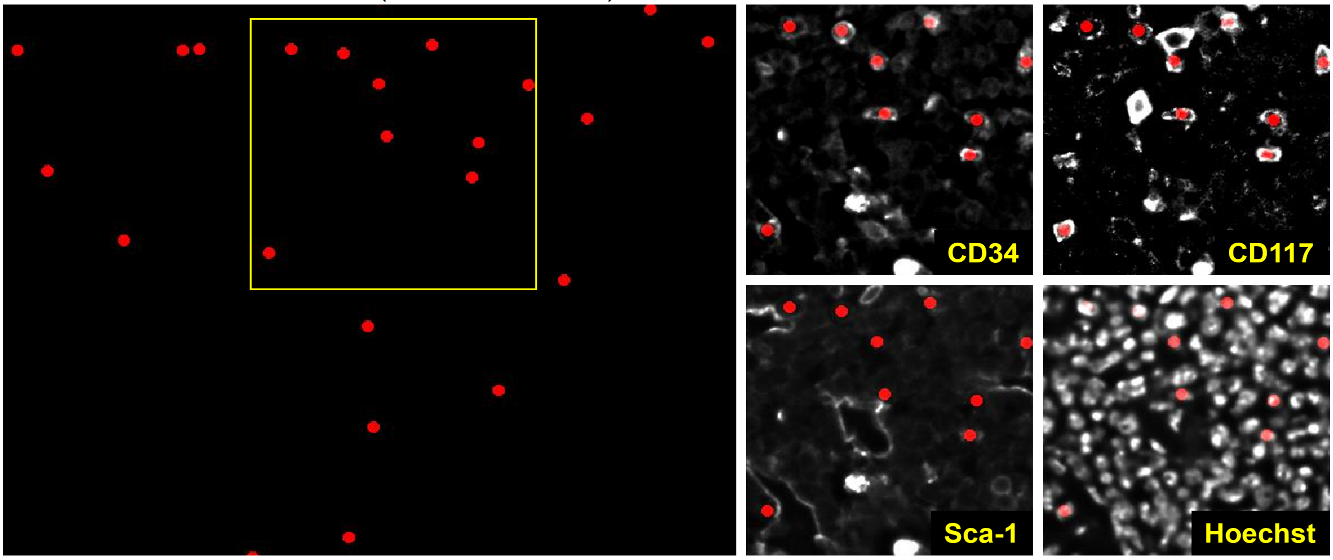

### Figure S3K

CD34$^{-}$CD117$^{+}$ HSPC cluster (lin$^{-}$CD34$^{-}$CD117$^{+}$)

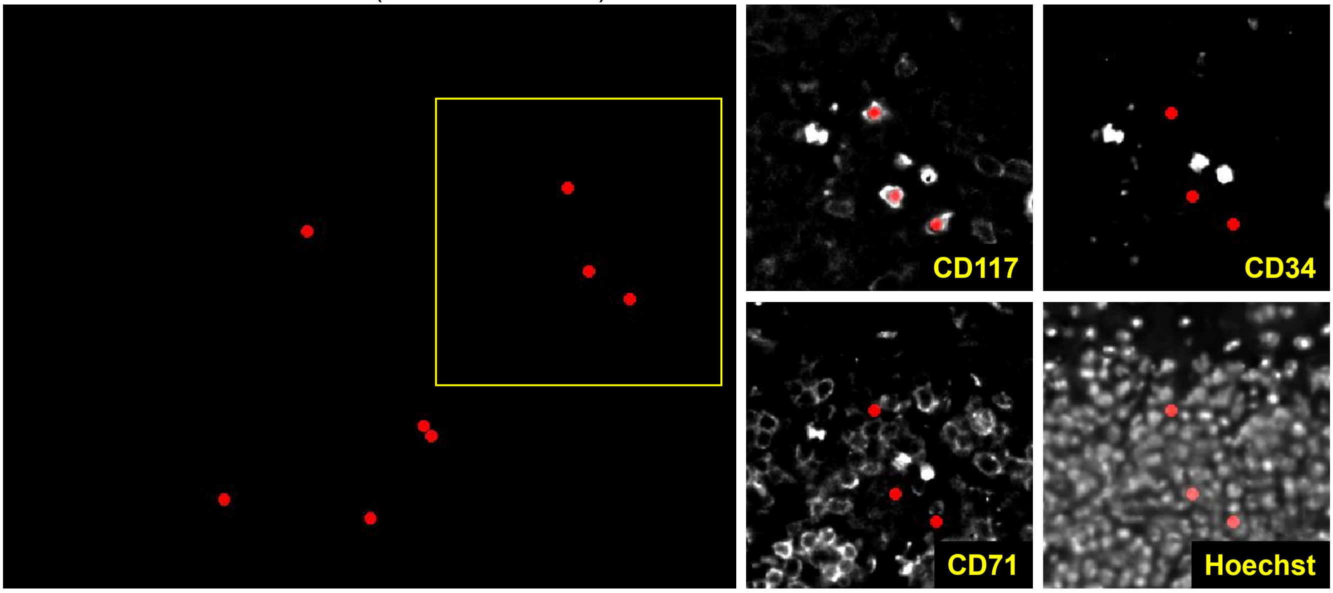


**K** EryA and EryB cell clusters (CD71$^{+}$TER119$^{+}$)

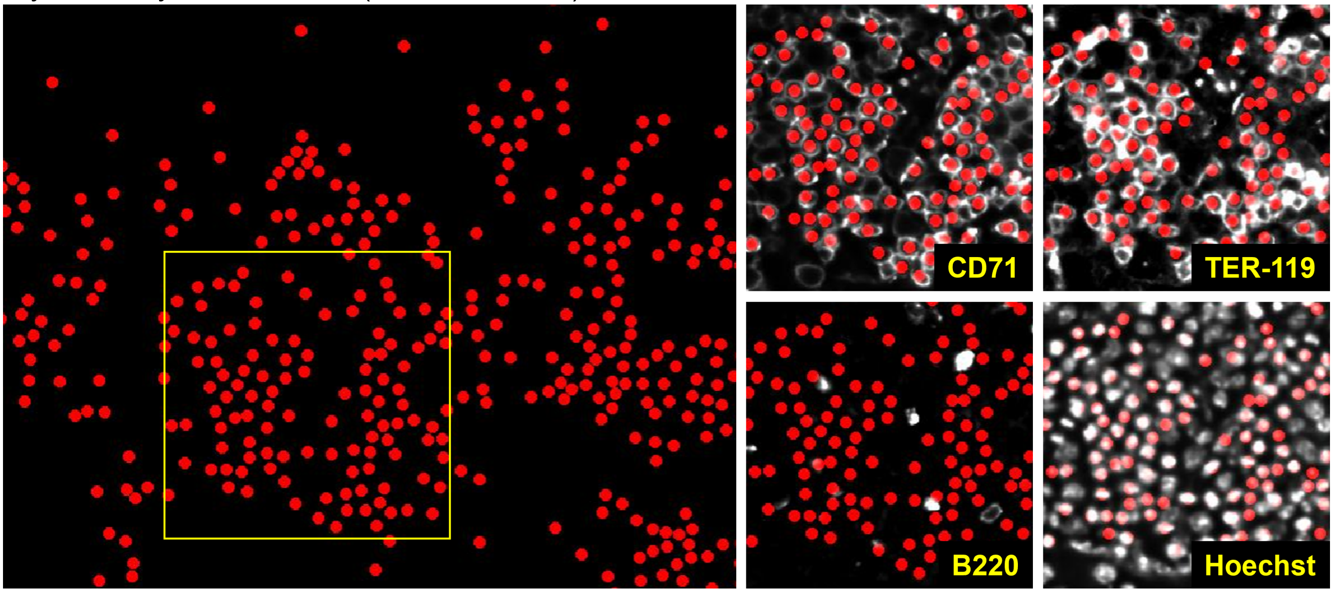


EryC cluster (CD71$^{-}$TER119$^{+}$)

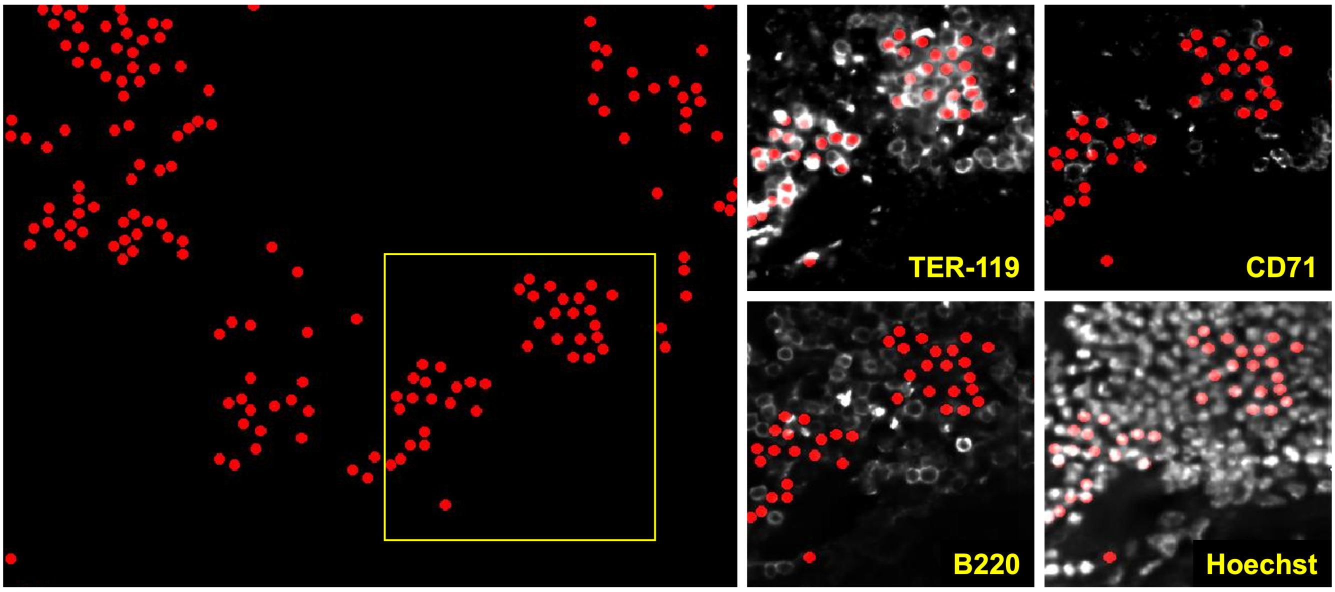

### Figure S3L-M

ProE cluster (CD71⁺TER119⁻)

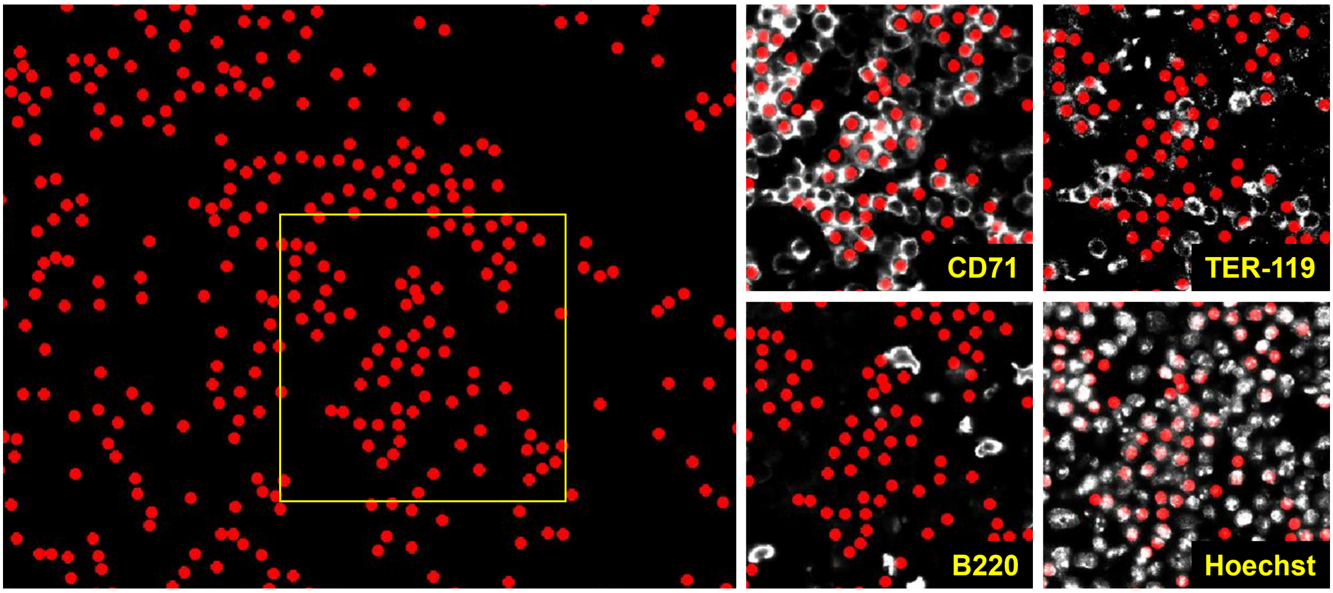


**L** EBI cluster (Macrophages surrounded by erythroblasts)

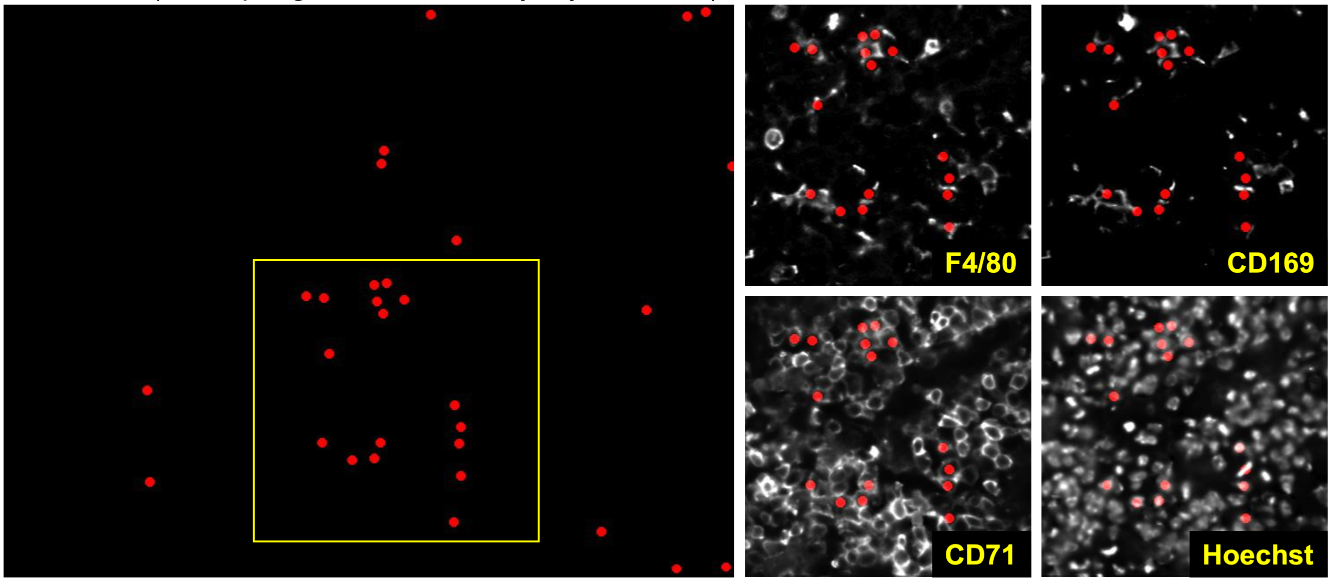


**M** Megakaryocyte cluster (CD41⁺)

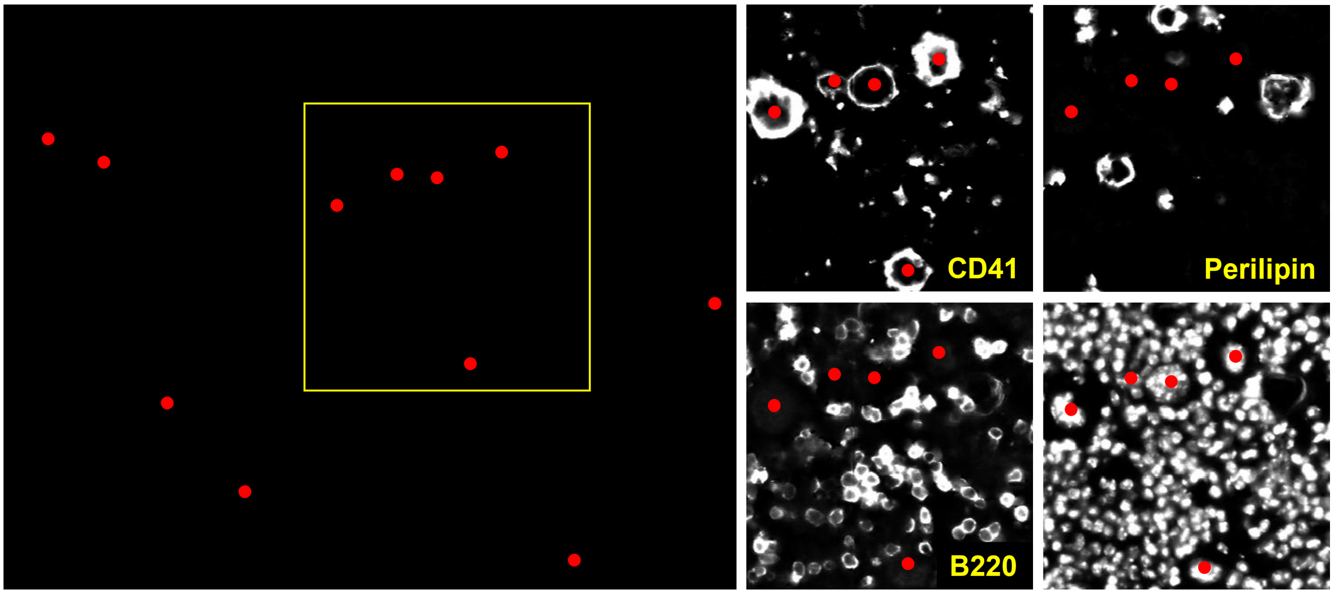

### Figure S3N-O

**N** B cell cluster (B220$^+$IgM$^-$)

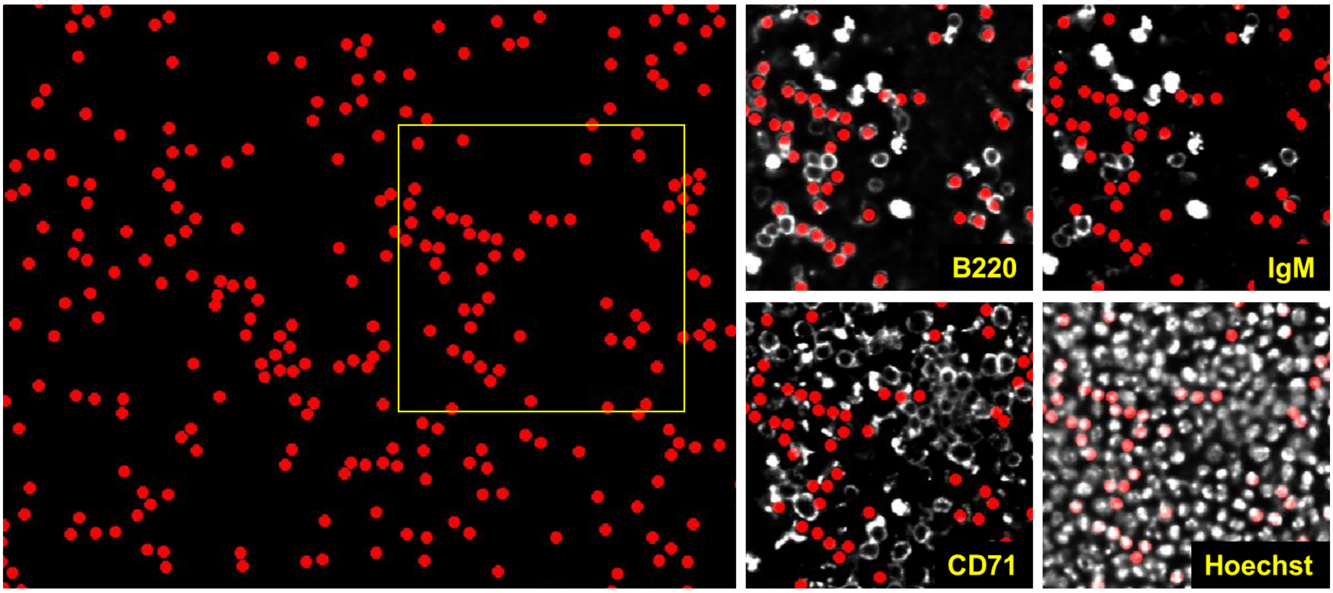


Plasma cell cluster (B220$^+$IgM$^+$)

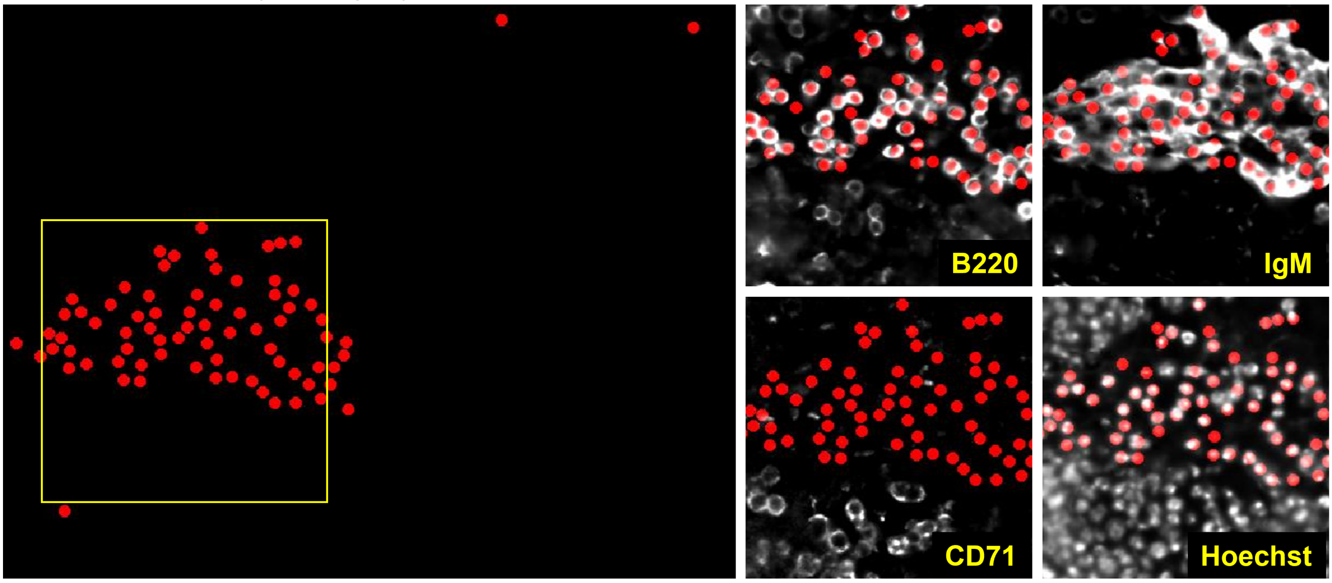


**O** NK cell cluster (CD3$^-$NKp46$^+$)

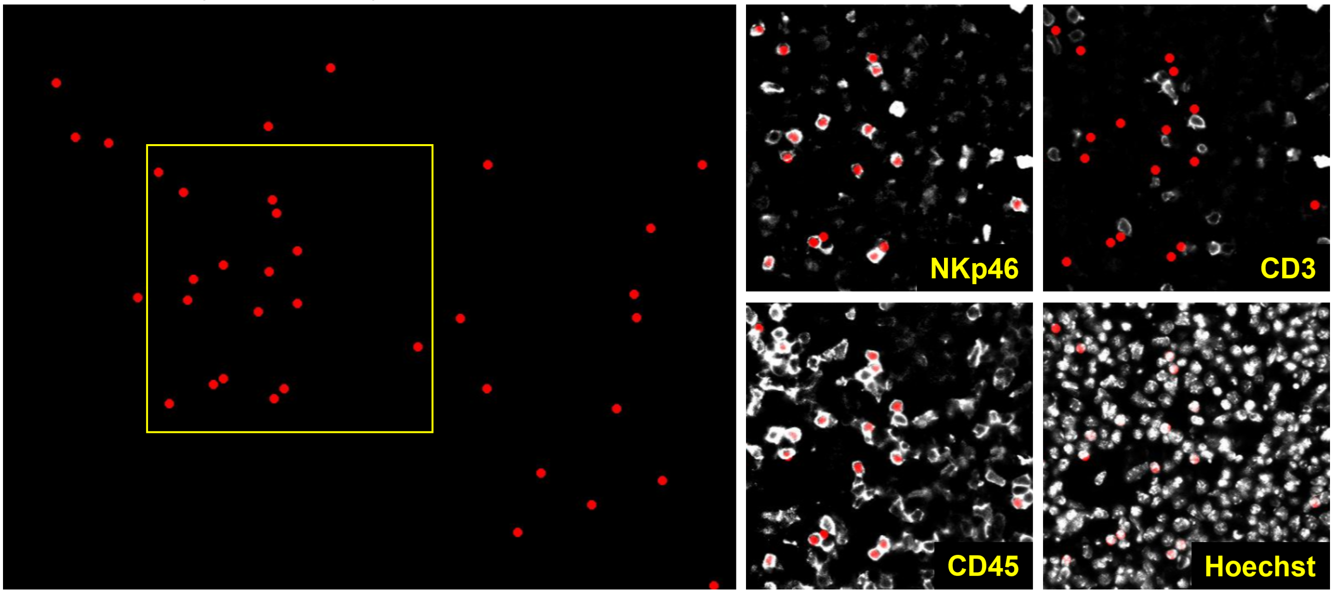

### Figure S3P

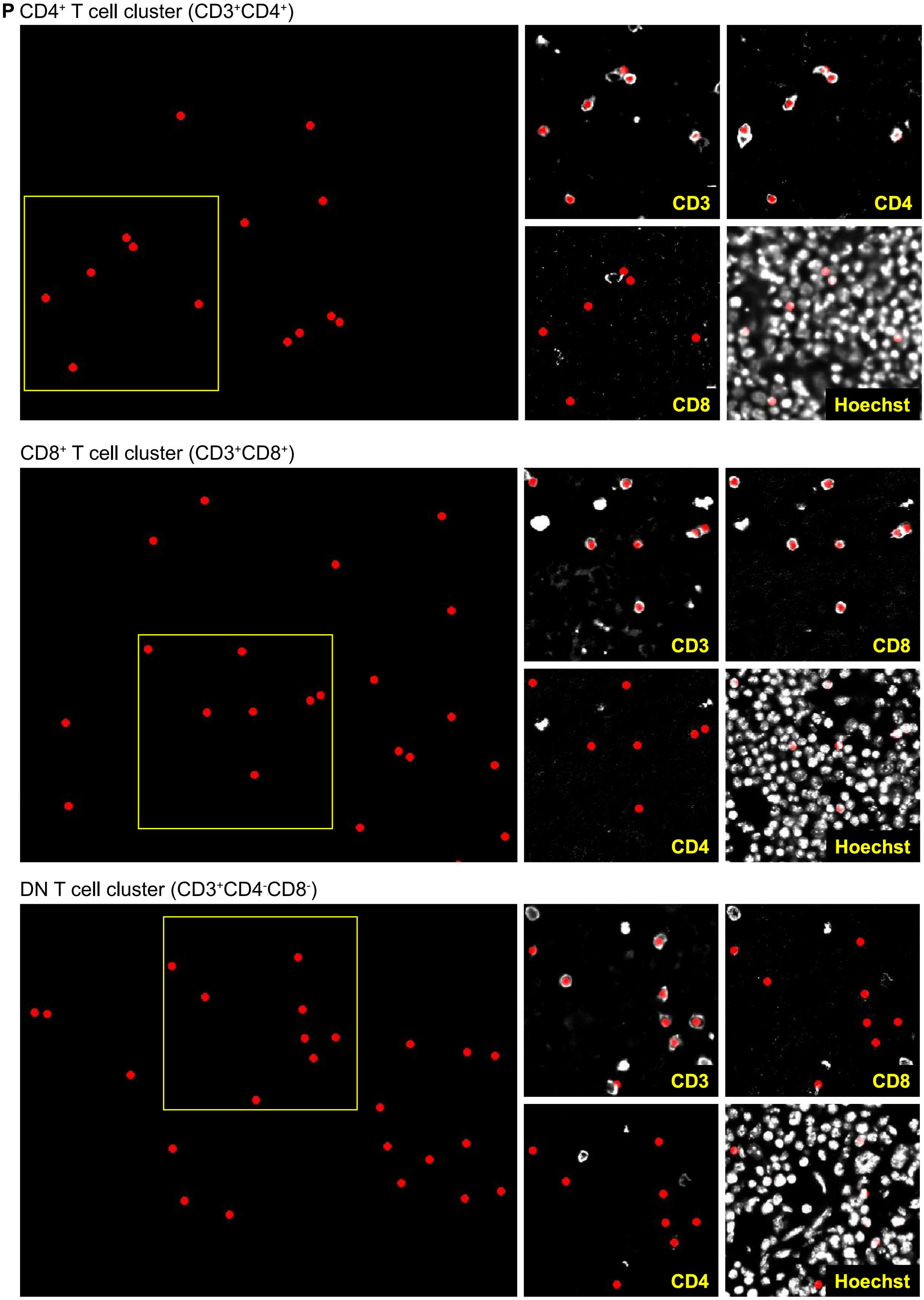
P CD4⁺ T cell cluster (CD3⁺CD4⁺)
CD3
CD4
CD8
Hoechst
CD8⁺ T cell cluster (CD3⁺CD8⁺)
CD3
CD8
CD4
Hoechst
DN T cell cluster (CD3⁺CD4⁻CD8⁻)
CD3
CD8
CD4
Hoechst

#### Figure S3Q-R

**Q** PD-1$^+$CD4$^+$ T cell cluster (PD-1$^+$CD3$^+$CD4$^+$)

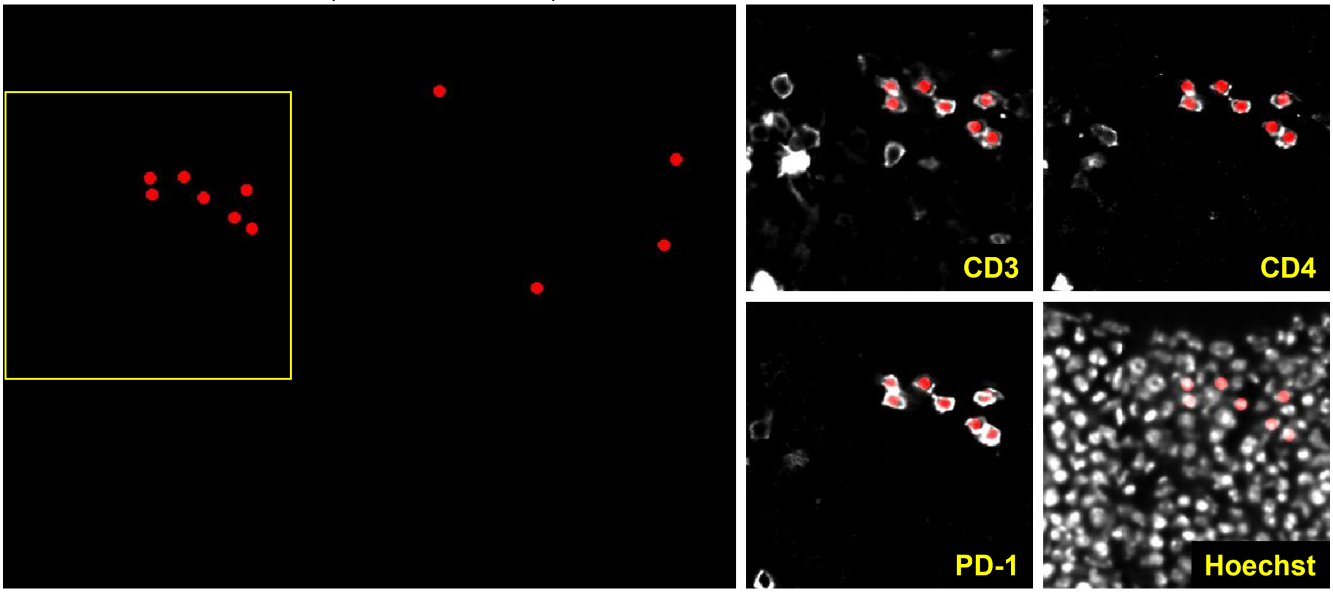


PD-1$^+$CD8$^+$ T cell cluster (PD-1$^+$CD3$^+$CD8$^+$)

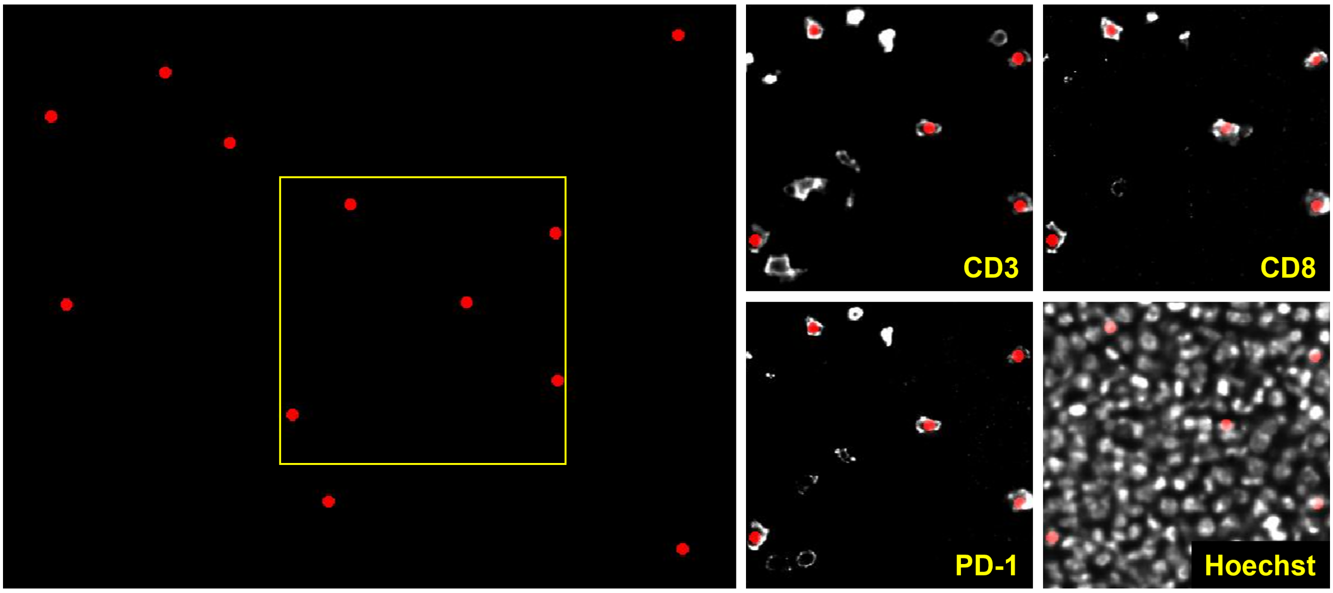


**R** Sinus cluster (CD31$^+$Sca-1$^-$)

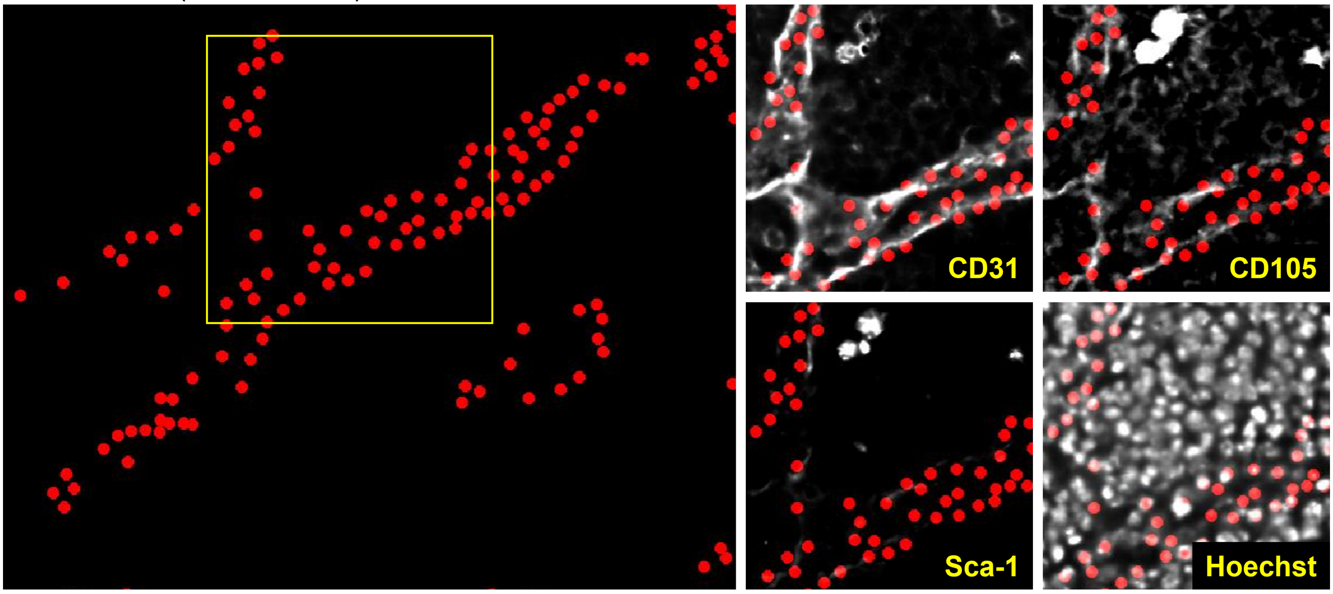

### Figure S3S-T

Arteriole cluster (CD31$^+$Sca-1$^+$)

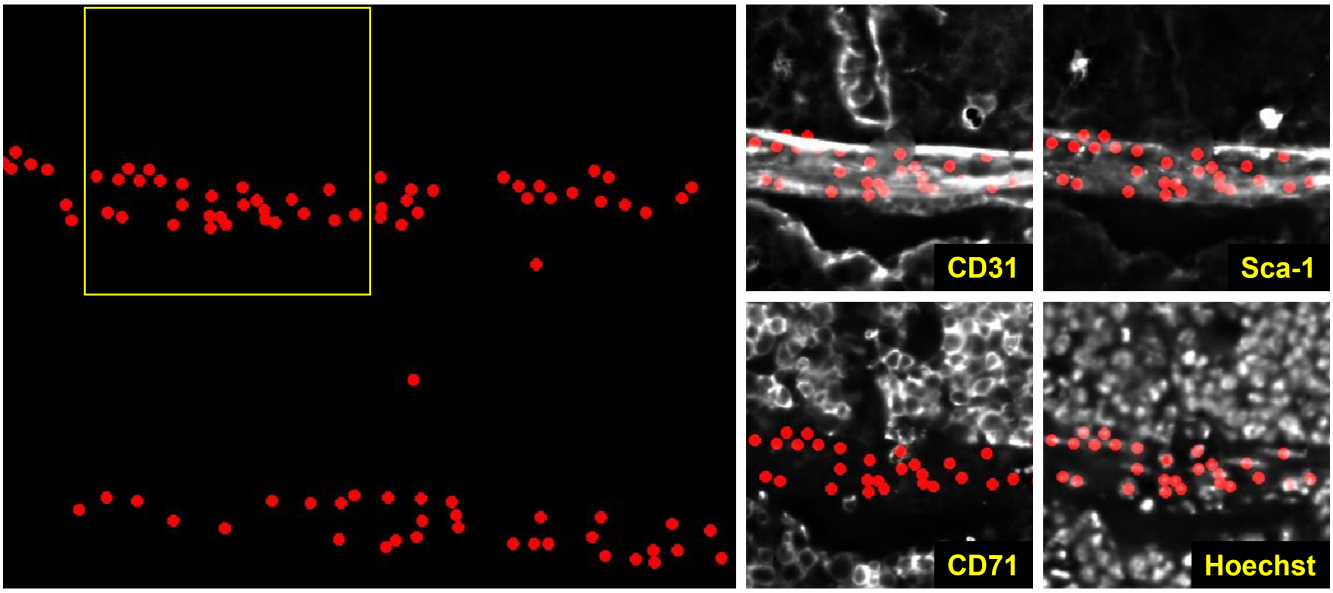


**S** LepR$^+$ pericyte cluster (LepR$^+$)

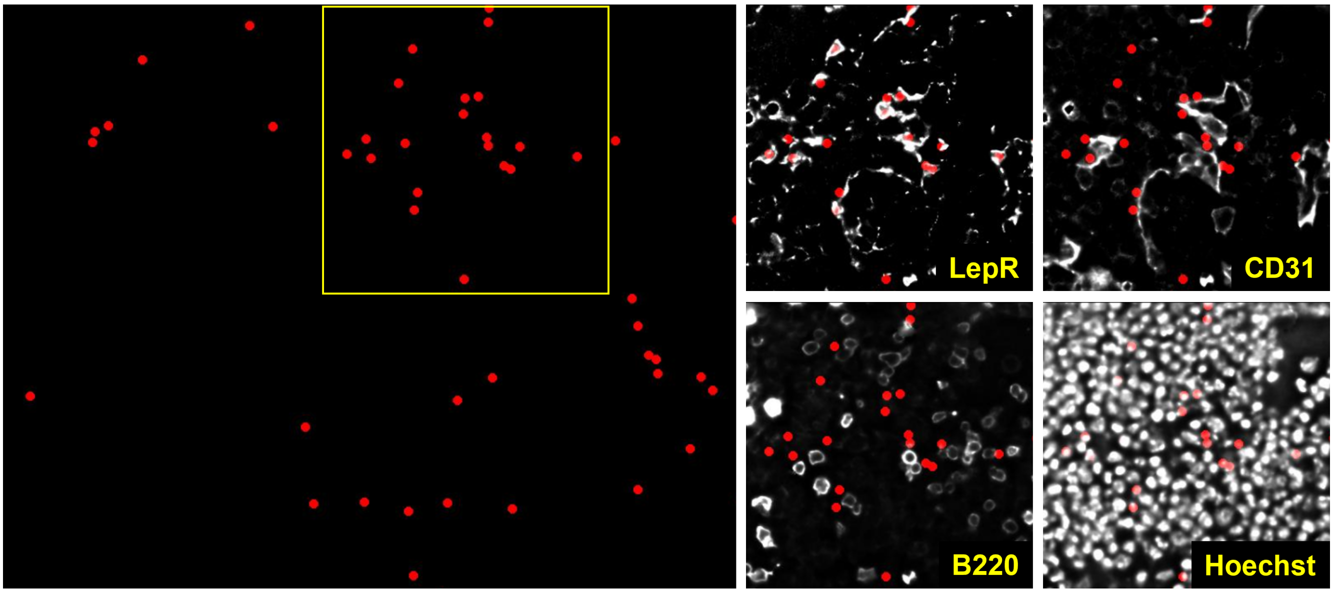


**T** Bone cell cluster (CD61$^+$ and/or osteopontin$^+$)

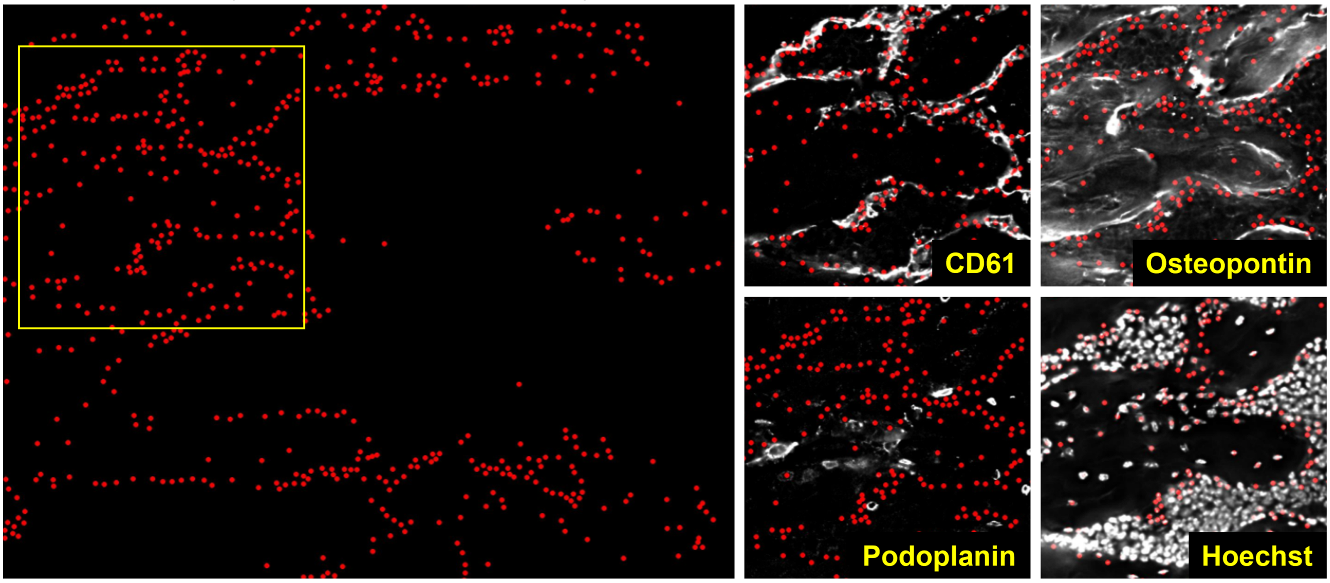

### Figure S3U-W

**U** Adipocyte cluster (Perilipin$^+$)

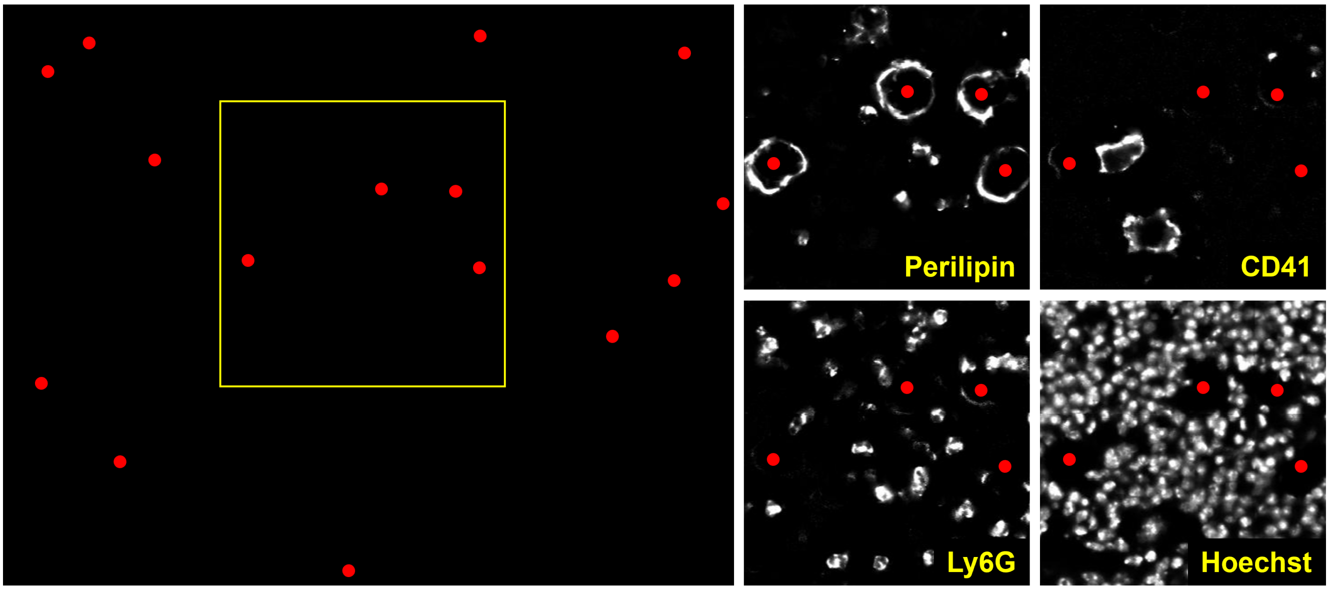


**V** Podo$^+$ stromal cell cluster (Podoplanin$^+$)

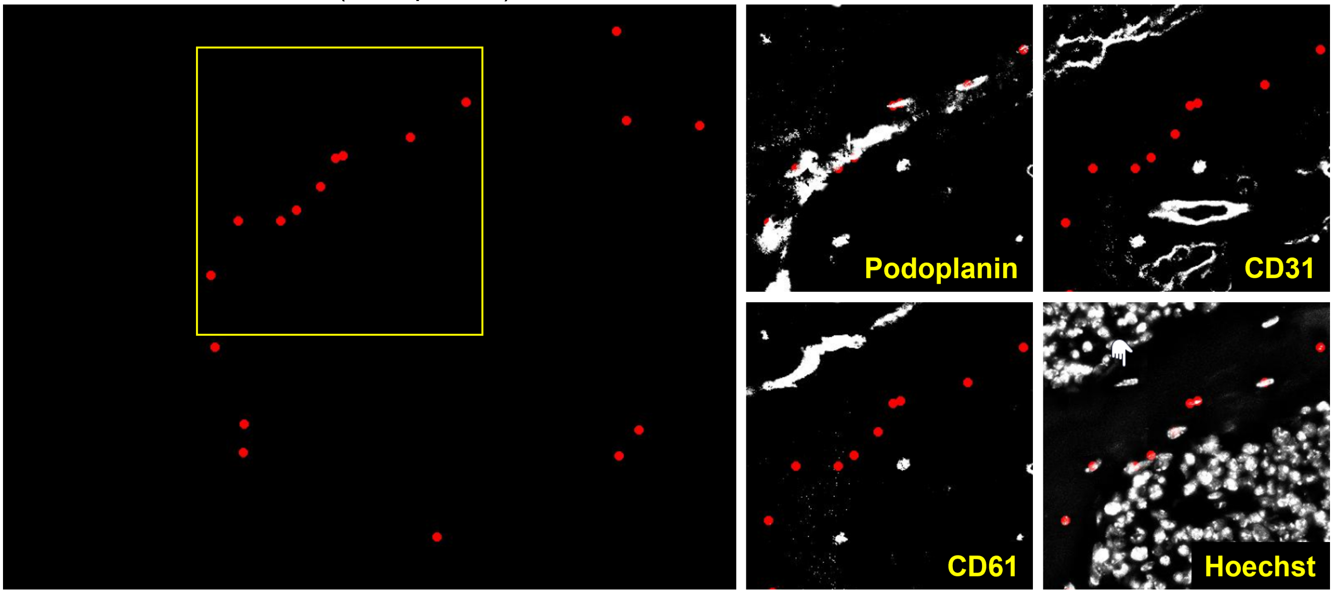


**W** Leukocyte cluster (CD45$^+$/negative for all other markers)

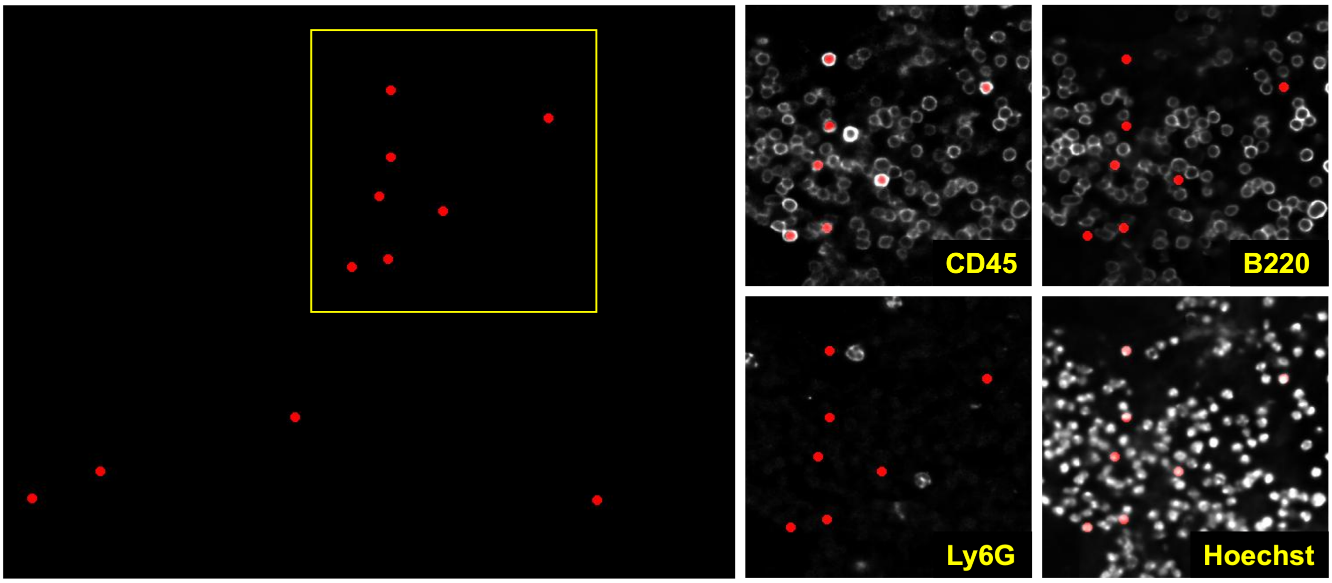

### Figure S3X

CD90⁺ leukocyte cluster (CD90⁺CD45⁺/negative for all other markers)

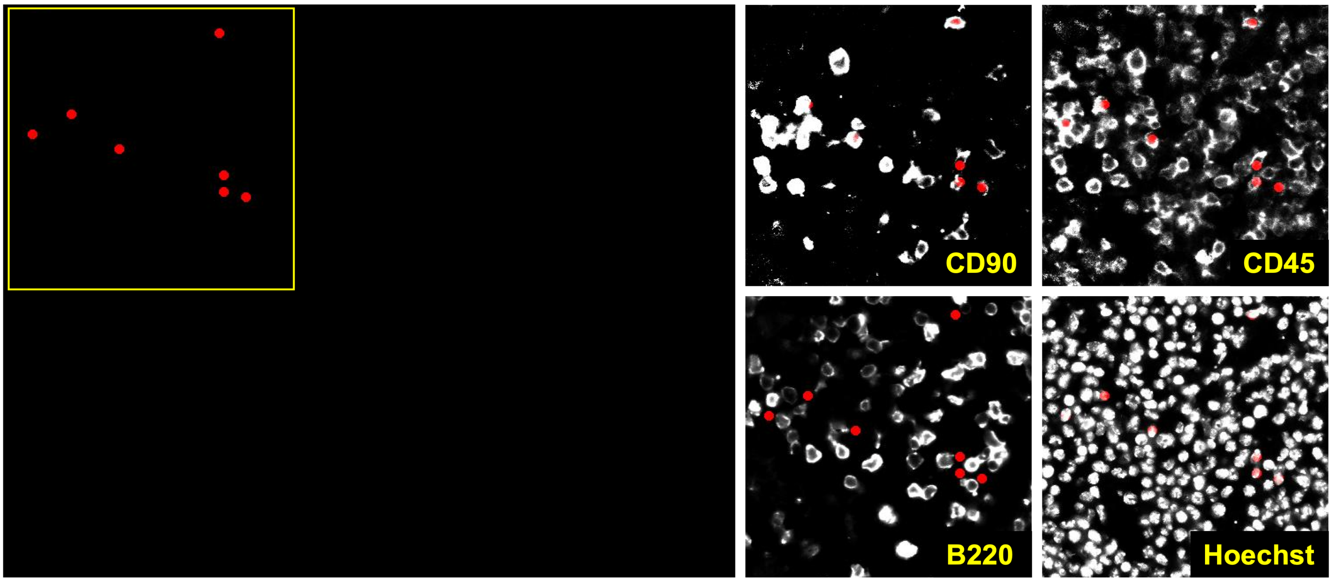


PD-L1⁺ leukocyte cluster (PD-L1⁺ CD45⁺/negative for all other markers)

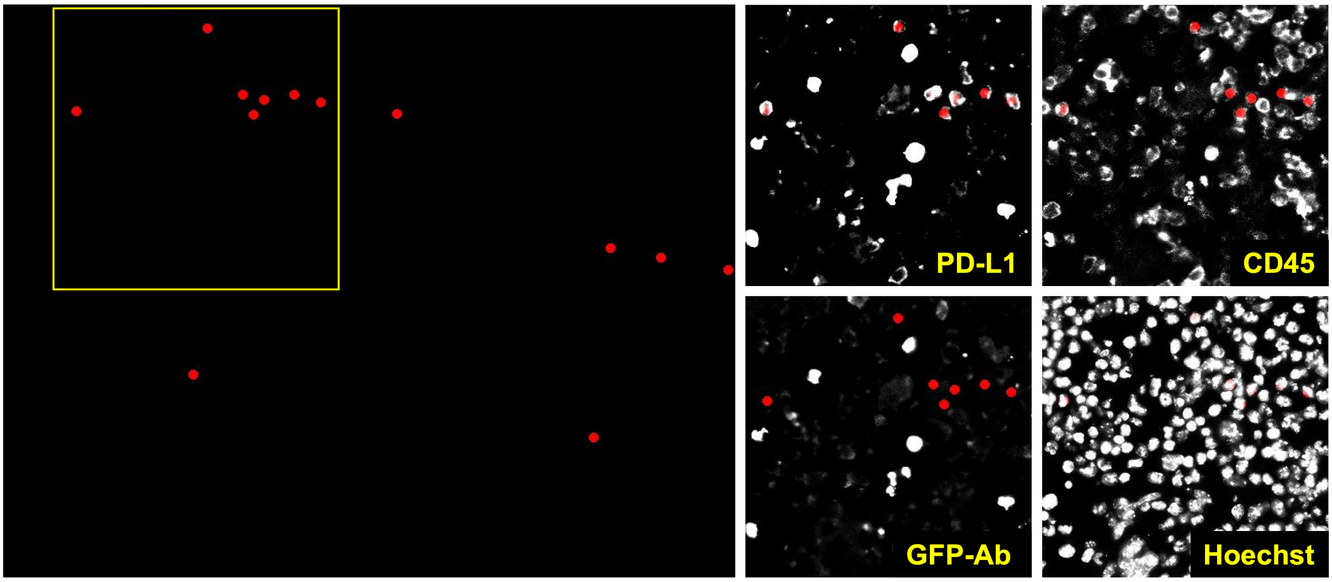


**X** Ki-67⁺ unidentified cluster (Ki-67⁺/negative for all other markers)

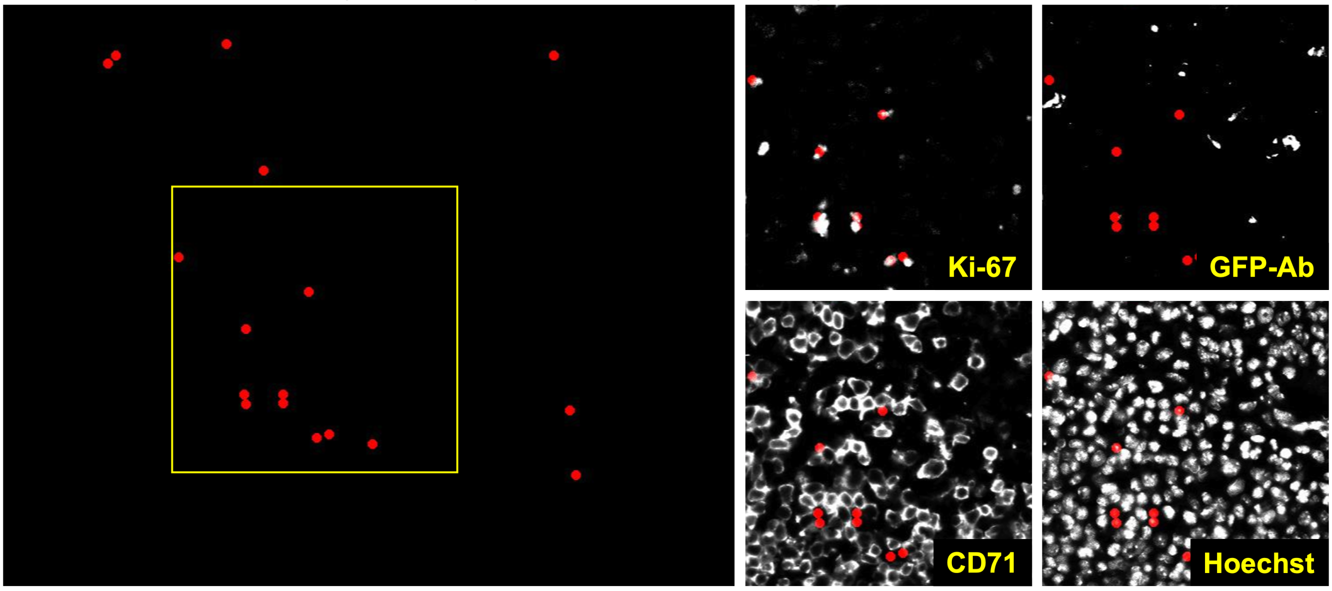

### Figure S4

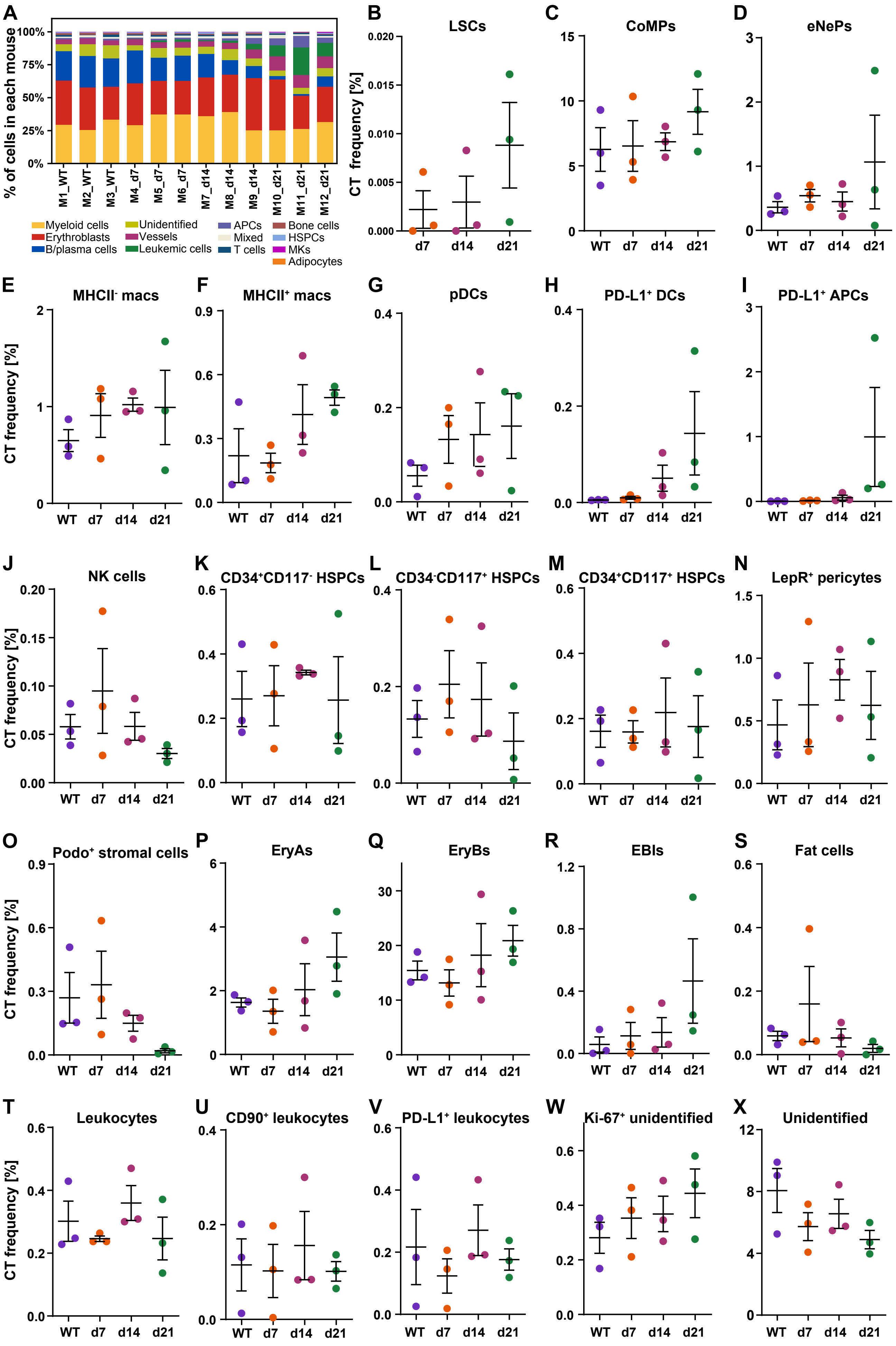

A
% of cells in each mouse
100%
75%
50%
25%
0%
M1_WT
M2_WT
M3_WT
M4_d7
M5_d7
M6_d7
M7_d14
M8_d14
M9_d14
M10_d21
M11_d21
M12_d21
Myeloid cells
Erythroblasts
B/plasma cells
Unidentified
Vessels
Leukemic cells
APCs
Mixed
T cells
Bone cells
HSPCs
MKs
Adipocytes
B LSCs
C CoMPs
D eNePs
E MHCII⁻ macs
F MHCII⁺ macs
G pDCs
H PD-L1⁺ DCs
I PD-L1⁺ APCs
J NK cells
K CD34⁺CD117⁻ HSPCs
L CD34⁻CD117⁺ HSPCs
M CD34⁺CD117⁺ HSPCs
N LepR⁺ pericytes
O Podo⁺ stromal cells
P EryAs
Q EryBs
R EBIs
S Fat cells
T Leukocytes
U CD90⁺ leukocytes
V PD-L1⁺ leukocytes
W Ki-67⁺ unidentified
X Unidentified
CT frequency [%]
WT
d7
d14
d21

### Figure S5

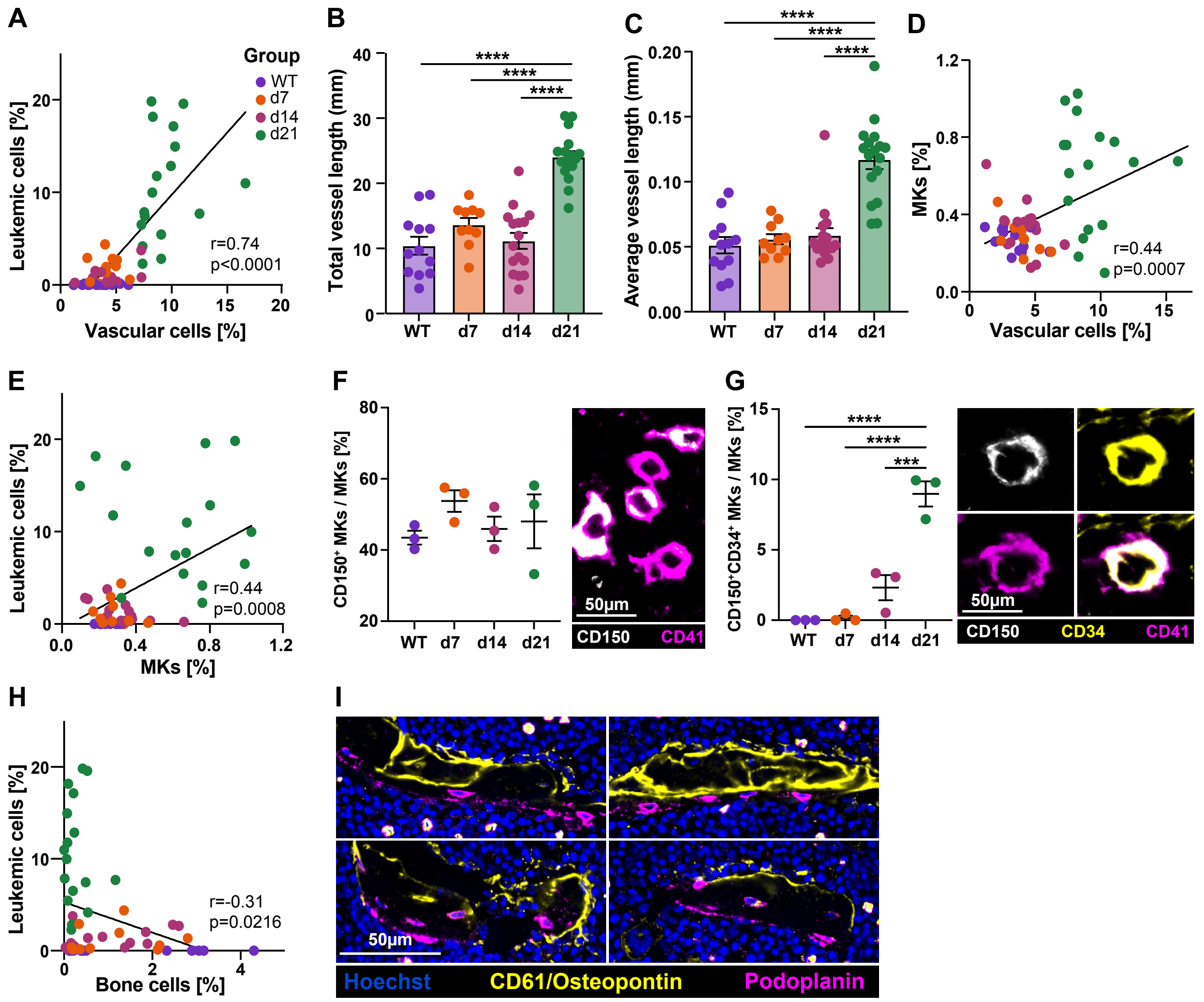
A
Group
WT
d7
d14
d21
Leukemic cells [%]
Vascular cells [%]
r=0.74
p<0.0001
B
Total vessel length (mm)
****
WT
d7
d14
d21
C
Average vessel length (mm)
D
MKs [%]
Vascular cells [%]
r=0.44
p=0.0007
E
Leukemic cells [%]
MKs [%]
r=0.44
p=0.0008
F
CD150⁺ MKs / MKs [%]
50µm
CD150
CD41
G
CD150⁺CD34⁺ MKs / MKs [%]
***
CD34
H
Leukemic cells [%]
Bone cells [%]
r=-0.31
p=0.0216
I
Hoechst
CD61/Osteopontin
Podoplanin

**Figure S6A-B**

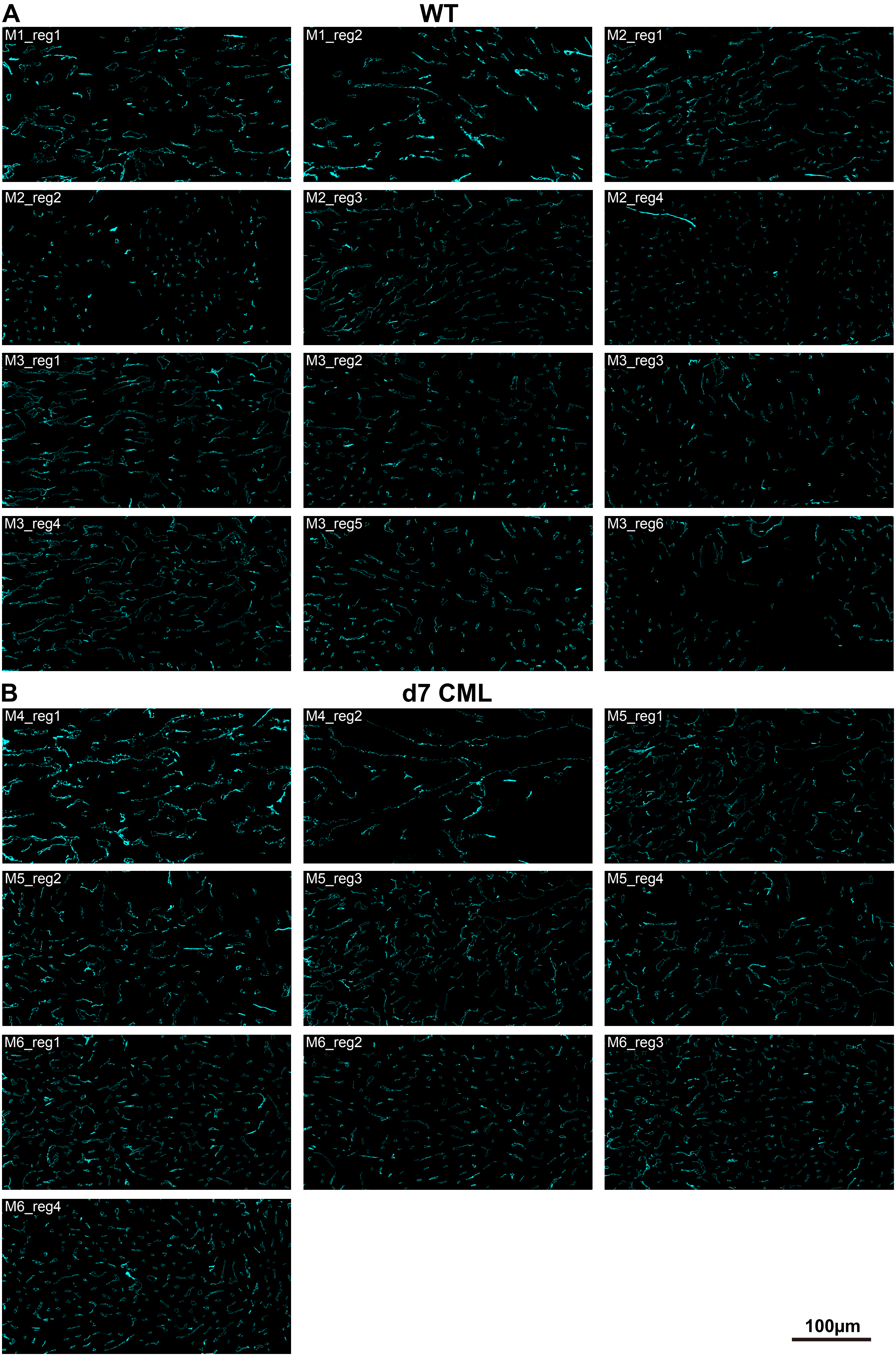
A
WT
M1_reg1
M1_reg2
M2_reg1
M2_reg2
M2_reg3
M2_reg4
M3_reg1
M3_reg2
M3_reg3
M3_reg4
M3_reg5
M3_reg6
B
d7 CML
M4_reg1
M4_reg2
M5_reg1
M5_reg2
M5_reg3
M5_reg4
M6_reg1
M6_reg2
M6_reg3
M6_reg4
100μm

### Figure S6C

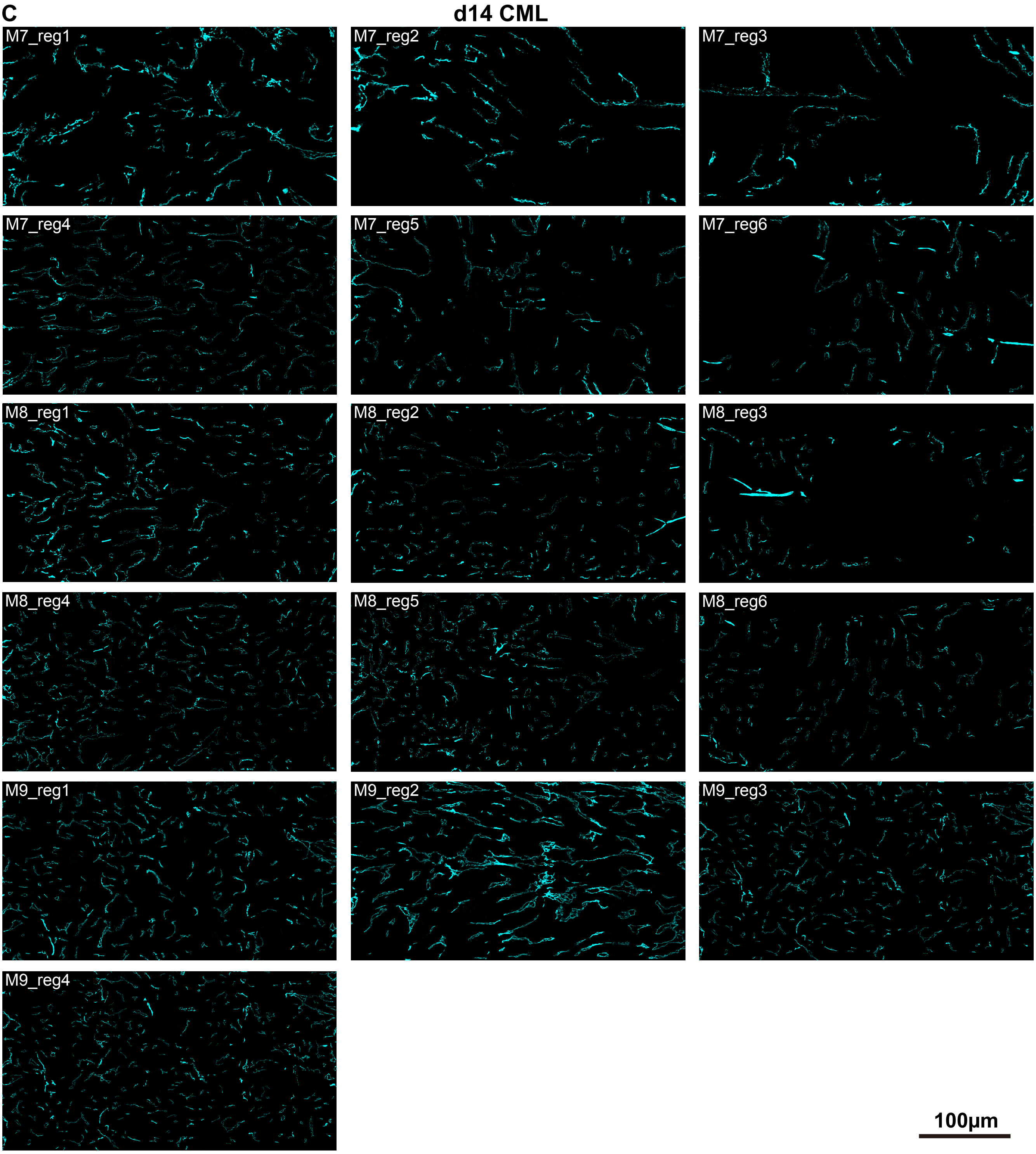
C
d14 CML
M7_reg1
M7_reg2
M7_reg3
M7_reg4
M7_reg5
M7_reg6
M8_reg1
M8_reg2
M8_reg3
M8_reg4
M8_reg5
M8_reg6
M9_reg1
M9_reg2
M9_reg3
M9_reg4
100µm

Figure S6D

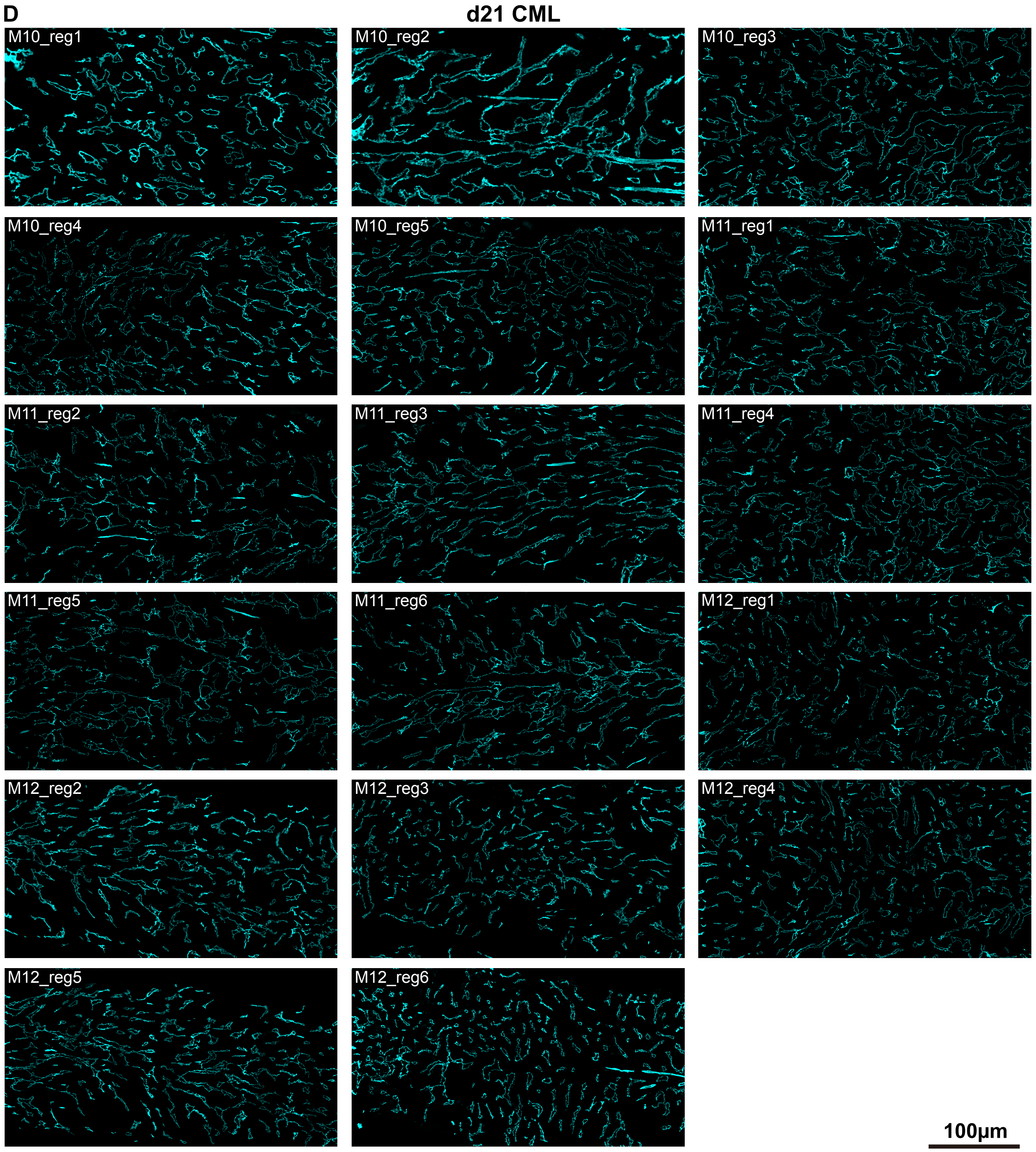
D
d21 CML
M10_reg1
M10_reg2
M10_reg3
M10_reg4
M10_reg5
M11_reg1
M11_reg2
M11_reg3
M11_reg4
M11_reg5
M11_reg6
M12_reg1
M12_reg2
M12_reg3
M12_reg4
M12_reg5
M12_reg6
100μm

### Figure S7

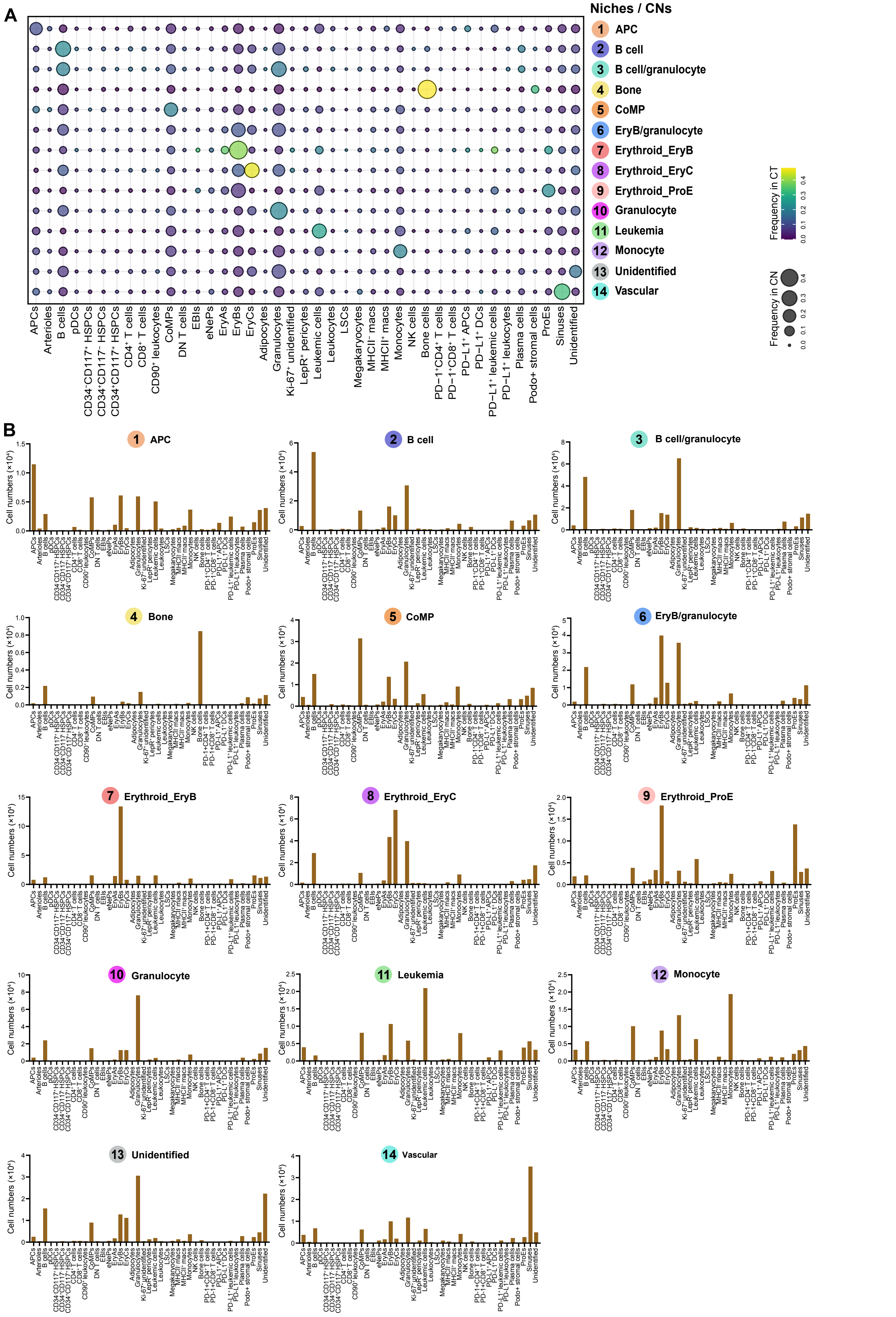
A
Niches / CNs
1 APC
2 B cell
3 B cell/granulocyte
4 Bone
5 CoMP
6 EryB/granulocyte
7 Erythroid_EryB
8 Erythroid_EryC
9 Erythroid_ProE
10 Granulocyte
11 Leukemia
12 Monocyte
13 Unidentified
14 Vascular
Frequency in CT
Frequency in CN
APCs
Arterioles
B cells
pDCs
CD34⁻CD117⁺ HSPCs
CD34⁺CD117⁻ HSPCs
CD34⁺CD117⁺ HSPCs
CD4⁺ T cells
CD8⁺ T cells
CD90⁺ leukocytes
CoMPs
DN T cells
EBIs
eNePs
EryAs
EryBs
EryCs
Adipocytes
Granulocytes
Ki-67⁺ unidentified
LepR⁺ pericytes
Leukemic cells
Leukocytes
LSCs
Megakaryocytes
MHCII⁻ macs
MHCII⁺ macs
Monocytes
NK cells
Bone cells
PD−1⁺CD4⁺ T cells
PD−1⁺CD8⁺ T cells
PD−L1⁺ APCs
PD−L1⁺ DCs
PD−L1⁺ leukemic cells
PD−L1⁺ leukocytes
Plasma cells
Podo+ stromal cells
ProEs
Sinuses
Unidentified
B
Cell numbers (×10⁴)

### Figure S8

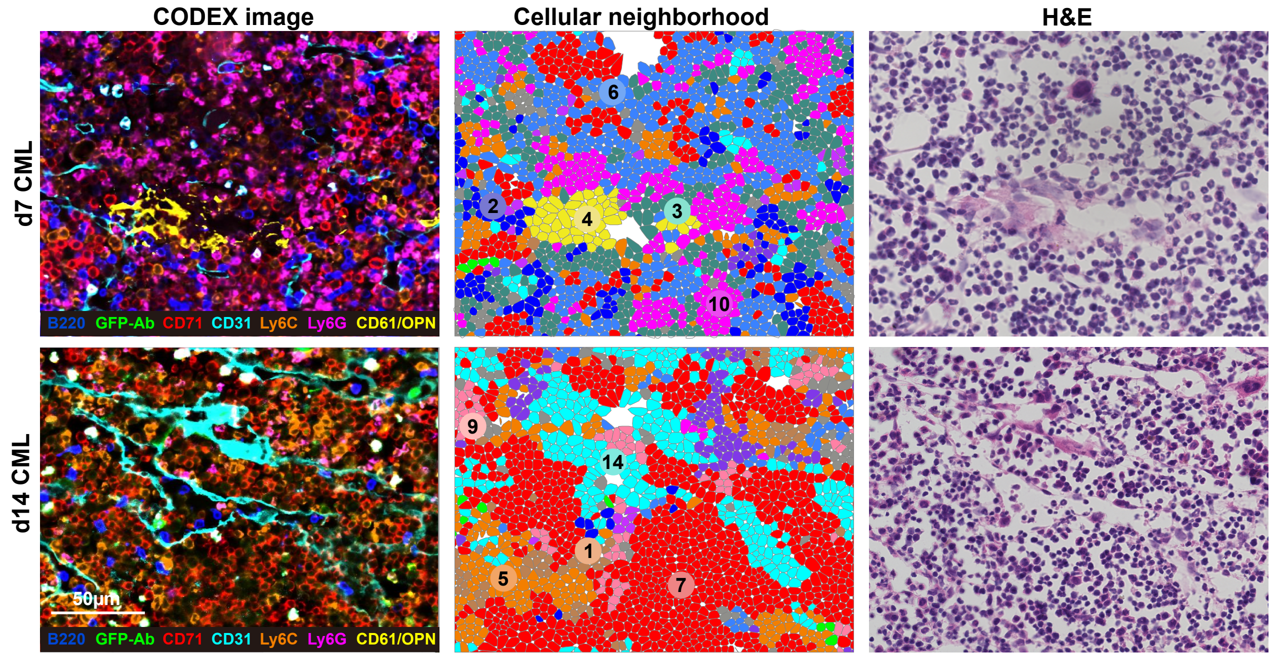
CODEX image
Cellular neighborhood
H&E
d7 CML
d14 CML
B220 GFP-Ab CD71 CD31 Ly6C Ly6G CD61/OPN
B220 GFP-Ab CD71 CD31 Ly6C Ly6G CD61/OPN
50μm
6
2
4
3
10
9
14
1
5
7

### Figure S9

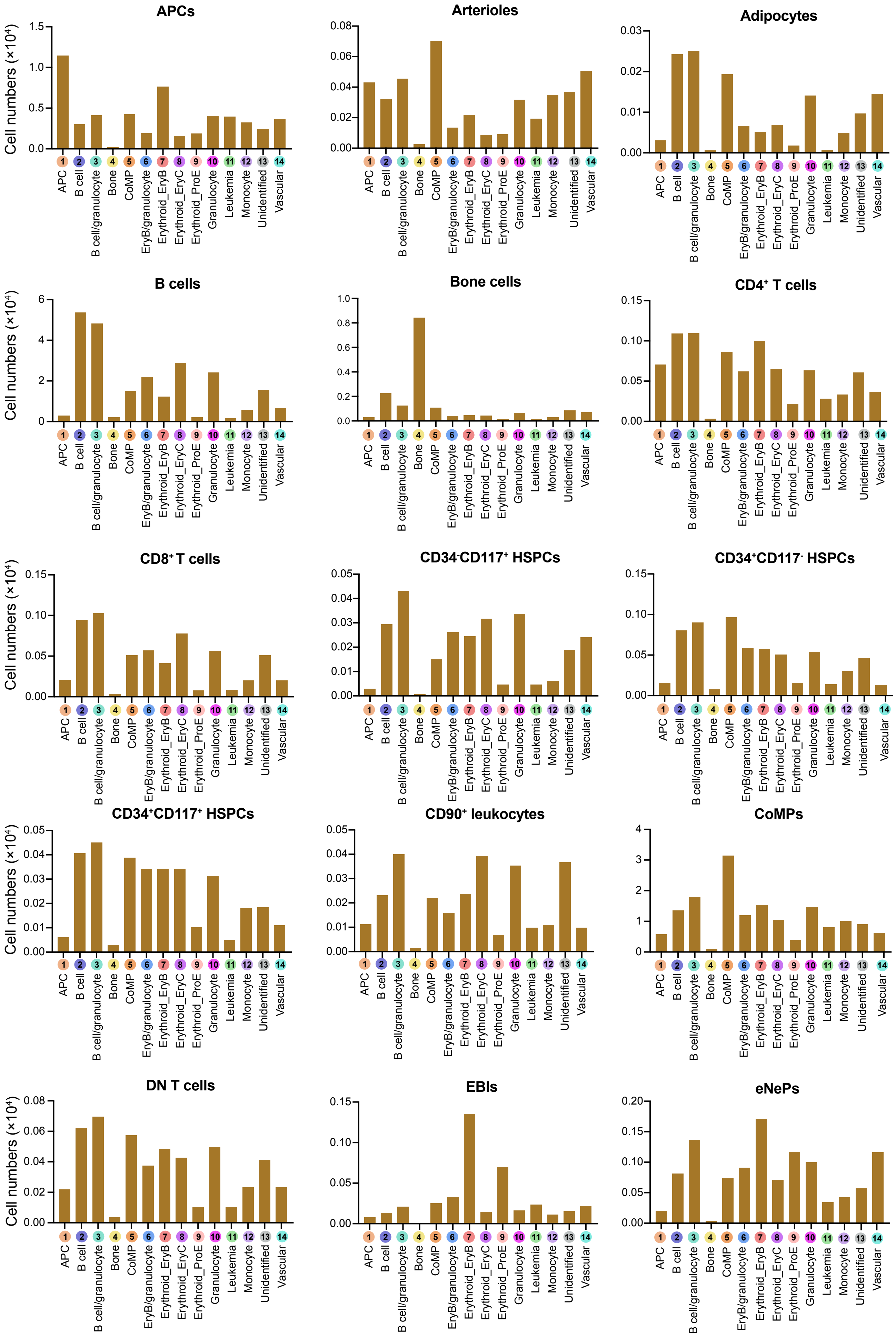
APCs
Arterioles
Adipocytes
B cells
Bone cells
CD4⁺ T cells
CD8⁺ T cells
CD34⁻CD117⁺ HSPCs
CD34⁺CD117⁻ HSPCs
CD34⁺CD117⁺ HSPCs
CD90⁺ leukocytes
CoMPs
DN T cells
EBIs
eNePs
Cell numbers (×10⁴)
APC
B cell
B cell/granulocyte
Bone
CoMP
EryB/granulocyte
Erythroid_EryB
Erythroid_EryC
Erythroid_ProE
Granulocyte
Leukemia
Monocyte
Unidentified
Vascular

### Figure S9

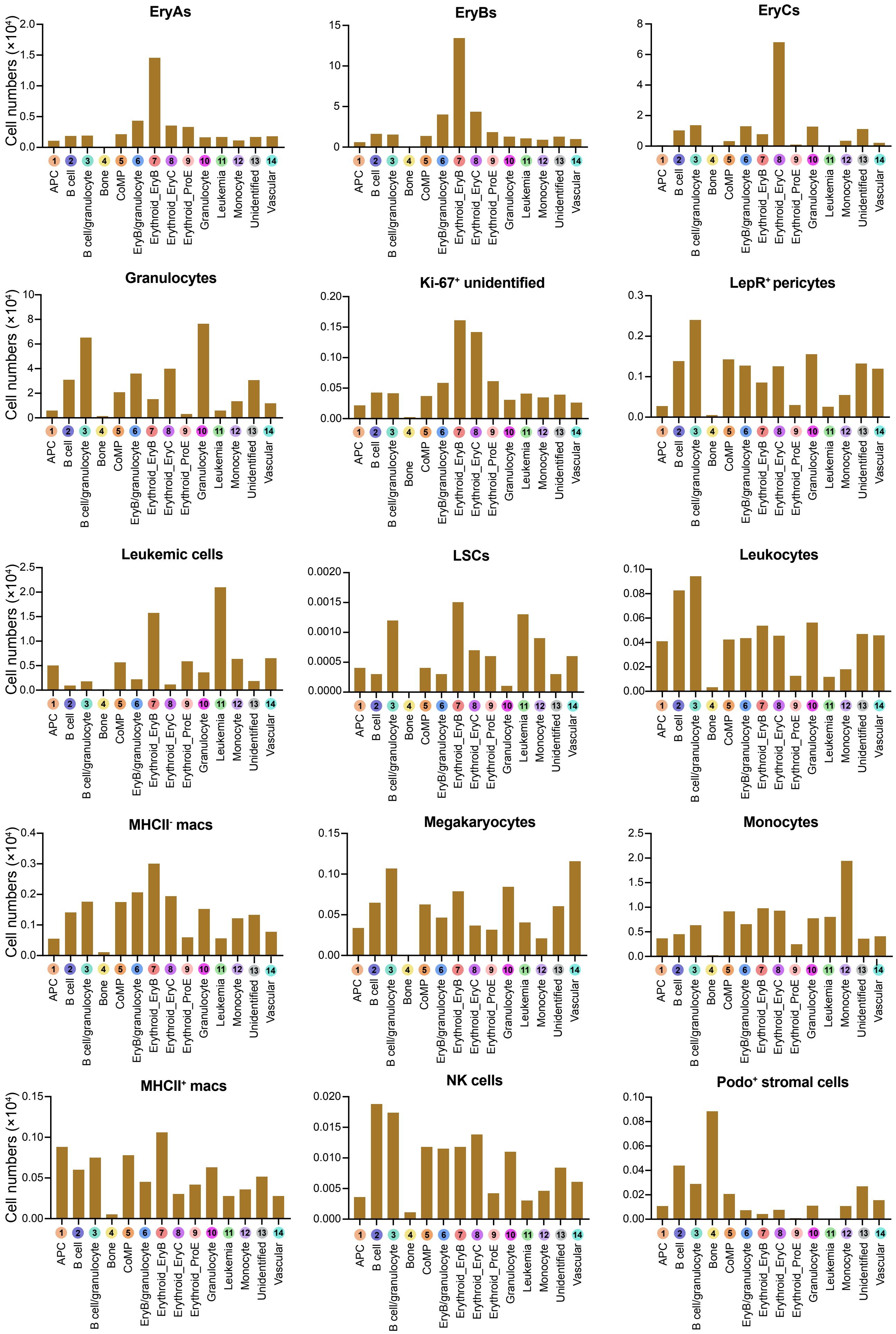

EryAs
EryBs
EryCs
Granulocytes
Ki-67⁺ unidentified
LepR⁺ pericytes
Leukemic cells
LSCs
Leukocytes
MHCII⁻ macs
Megakaryocytes
Monocytes
MHCII⁺ macs
NK cells
Podo⁺ stromal cells
Cell numbers (×10⁴)
APC
B cell
B cell/granulocyte
Bone
CoMP
EryB/granulocyte
Erythroid_EryB
Erythroid_EryC
Erythroid_ProE
Granulocyte
Leukemia
Monocyte
Unidentified
Vascular

### Figure S9

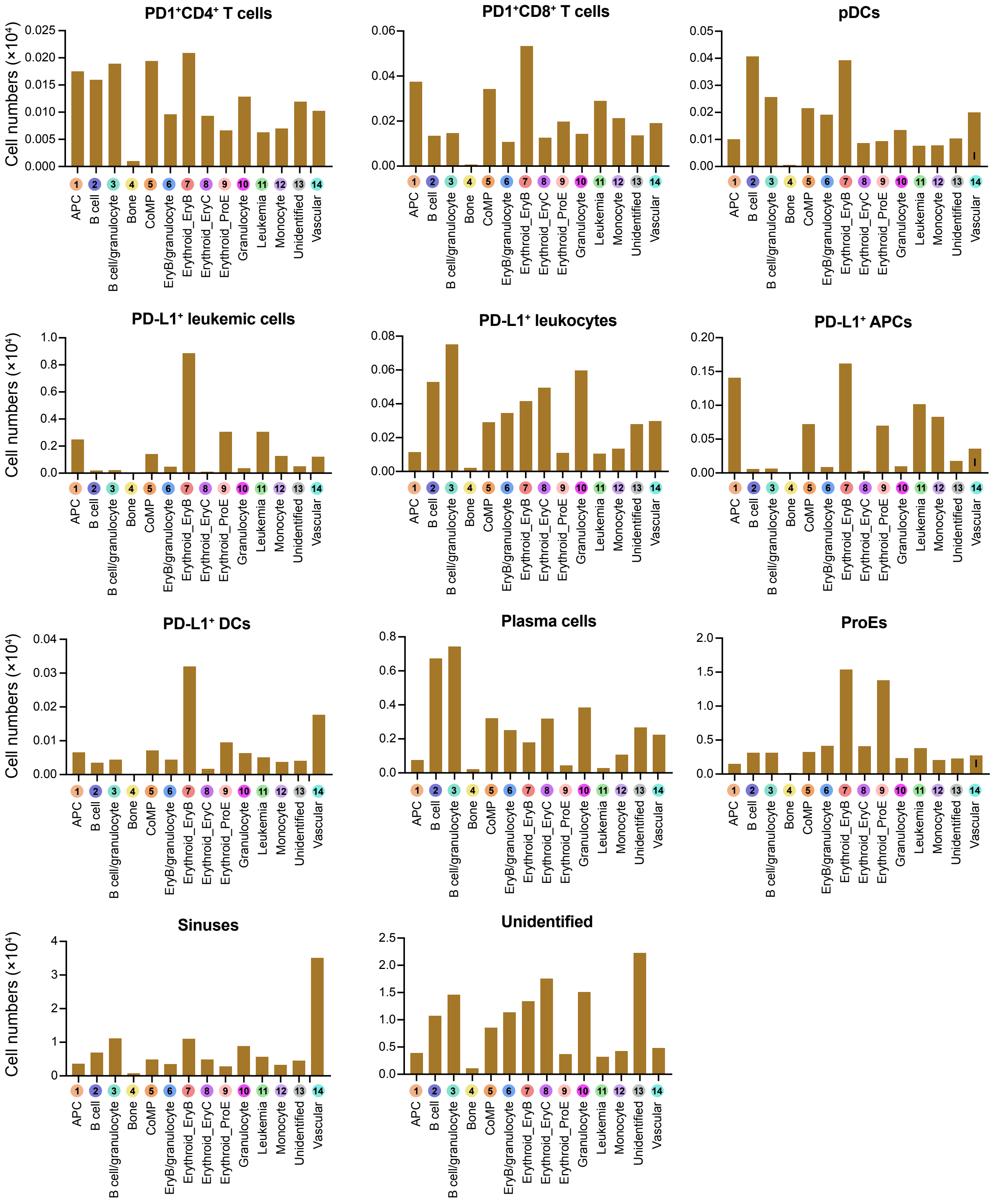

PD1⁺CD4⁺ T cells
PD1⁺CD8⁺ T cells
pDCs
PD-L1⁺ leukemic cells
PD-L1⁺ leukocytes
PD-L1⁺ APCs
PD-L1⁺ DCs
Plasma cells
ProEs
Sinuses
Unidentified
Cell numbers (×10⁴)
APC
B cell
B cell/granulocyte
Bone
CoMP
EryB/granulocyte
Erythroid_EryB
Erythroid_EryC
Erythroid_ProE
Granulocyte
Leukemia
Monocyte
Unidentified
Vascular

### Figure S10A-B

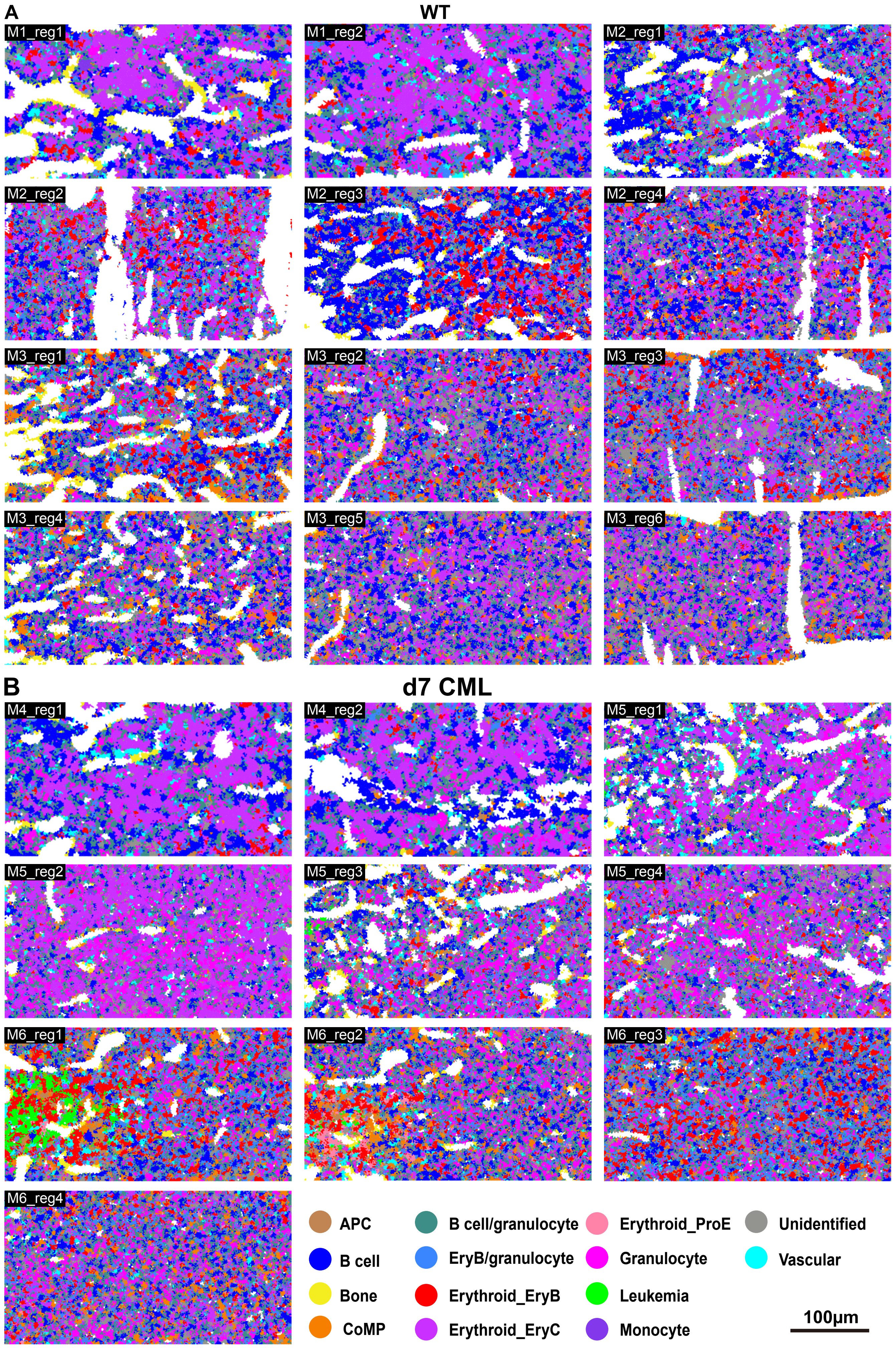

### Figure S10C

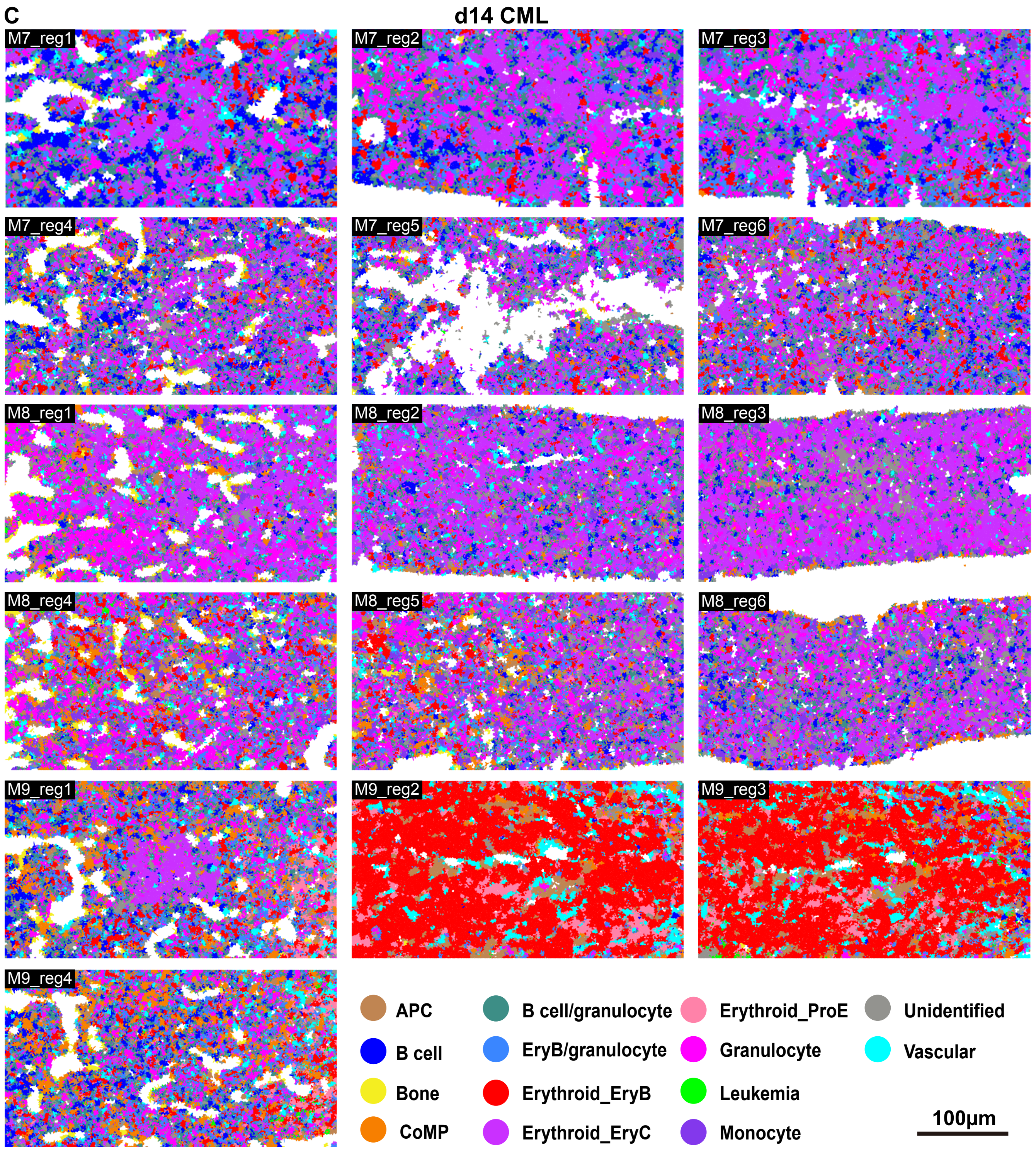
C
d14 CML
M7_reg1
M7_reg2
M7_reg3
M7_reg4
M7_reg5
M7_reg6
M8_reg1
M8_reg2
M8_reg3
M8_reg4
M8_reg5
M8_reg6
M9_reg1
M9_reg2
M9_reg3
M9_reg4
APC
B cell/granulocyte
Erythroid_ProE
Unidentified
B cell
EryB/granulocyte
Granulocyte
Vascular
Bone
Erythroid_EryB
Leukemia
CoMP
Erythroid_EryC
Monocyte
100μm

### Figure S10D

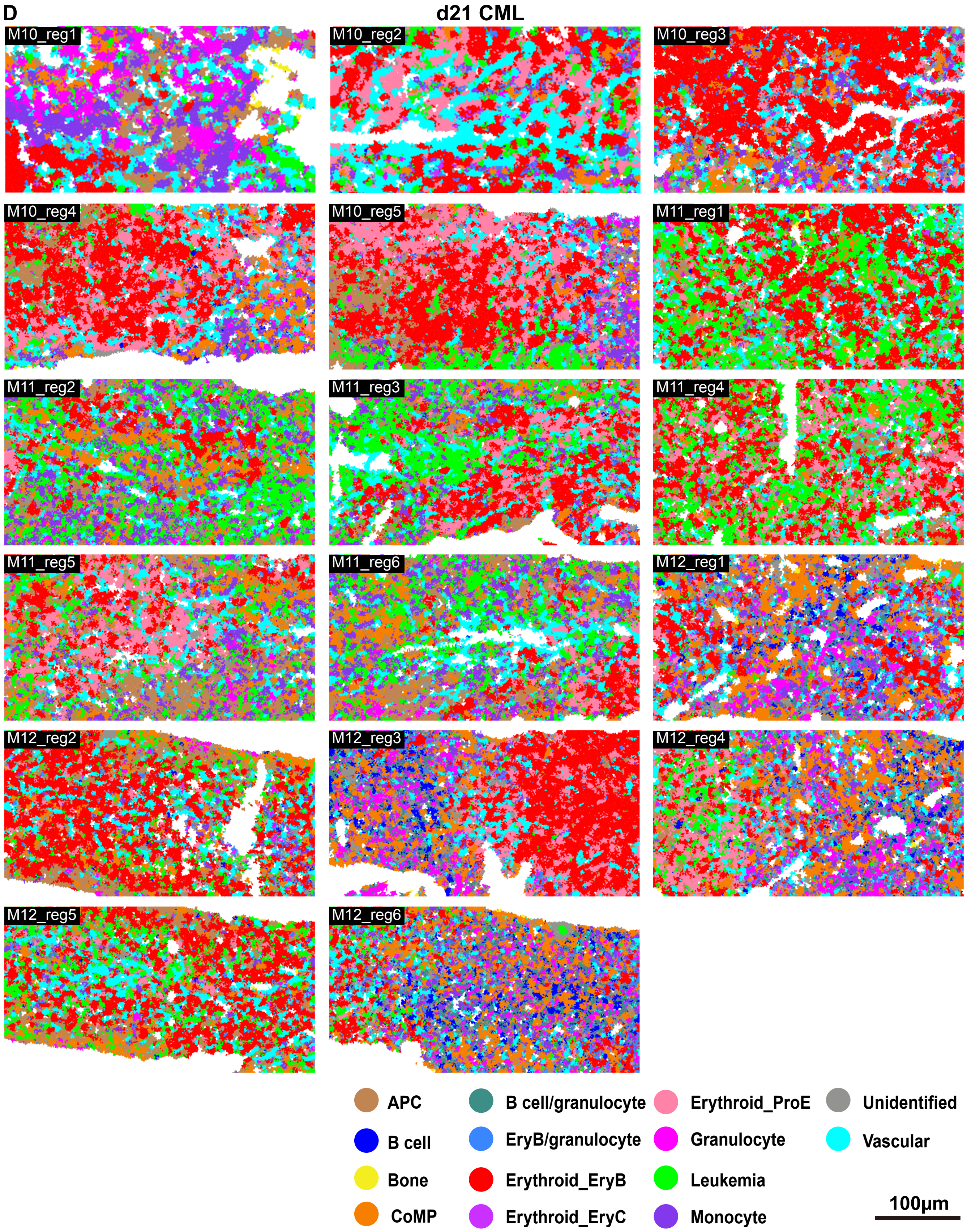
D
d21 CML
M10_reg1
M10_reg2
M10_reg3
M10_reg4
M10_reg5
M11_reg1
M11_reg2
M11_reg3
M11_reg4
M11_reg5
M11_reg6
M12_reg1
M12_reg2
M12_reg3
M12_reg4
M12_reg5
M12_reg6
APC
B cell
Bone
CoMP
B cell/granulocyte
EryB/granulocyte
Erythroid_EryB
Erythroid_EryC
Erythroid_ProE
Granulocyte
Leukemia
Monocyte
Unidentified
Vascular
100μm

### Figure S11

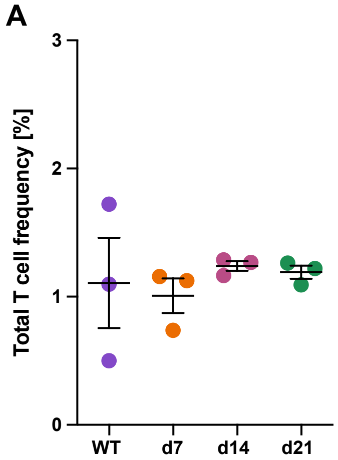

A
Total T cell frequency [%]
0
1
2
3
WT
d7
d14
d21


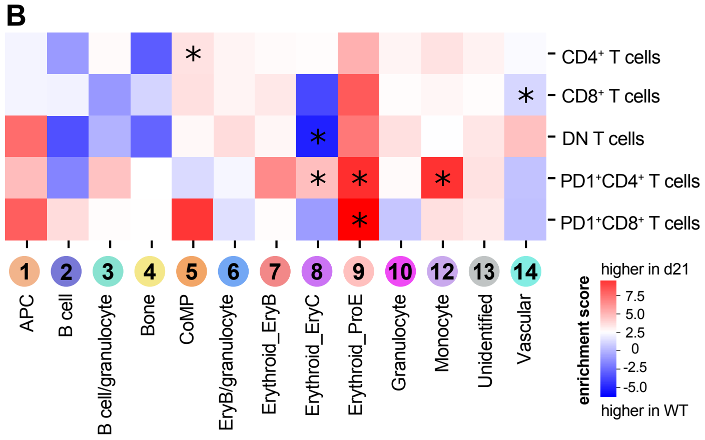

B
CD4⁺ T cells
CD8⁺ T cells
DN T cells
PD1⁺CD4⁺ T cells
PD1⁺CD8⁺ T cells
1 APC
2 B cell
3 B cell/granulocyte
4 Bone
5 CoMP
6 EryB/granulocyte
7 Erythroid_EryB
8 Erythroid_EryC
9 Erythroid_ProE
10 Granulocyte
12 Monocyte
13 Unidentified
14 Vascular
higher in d21
7.5
5.0
2.5
0
-2.5
-5.0
higher in WT
enrichment score

#### Figure S12

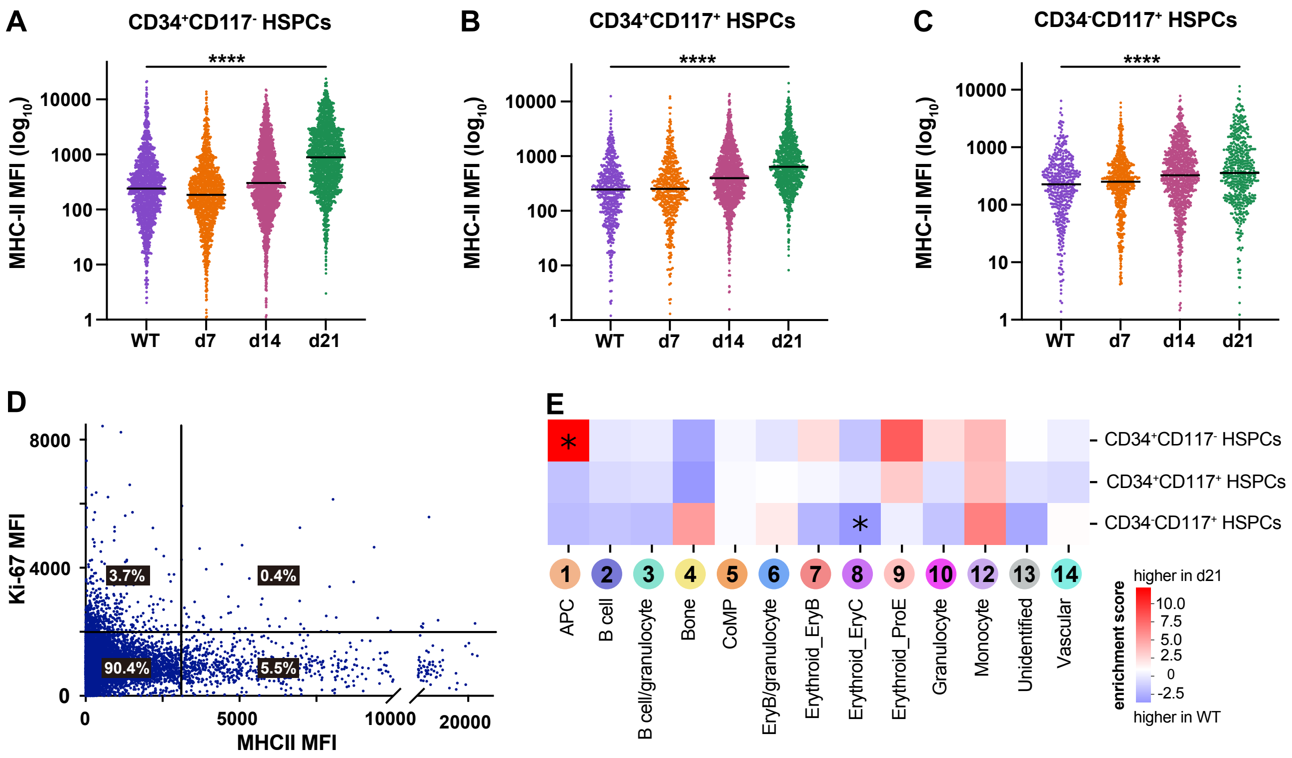
A
CD34⁺CD117⁻ HSPCs
B
CD34⁺CD117⁺ HSPCs
C
CD34⁻CD117⁺ HSPCs
****
MHC-II MFI (log10)
10000
1000
100
10
1
WT
d7
d14
d21
D
Ki-67 MFI
8000
4000
0
3.7%
0.4%
90.4%
5.5%
0
5000
10000
20000
MHCII MFI
E
CD34⁺CD117⁻ HSPCs
CD34⁺CD117⁺ HSPCs
CD34⁻CD117⁺ HSPCs
1 APC
2 B cell
3 B cell/granulocyte
4 Bone
5 CoMP
6 EryB/granulocyte
7 Erythroid_EryB
8 Erythroid_EryC
9 Erythroid_ProE
10 Granulocyte
12 Monocyte
13 Unidentified
14 Vascular
higher in d21
enrichment score
10.0
7.5
5.0
2.5
0
-2.5
higher in WT

### Figure S13

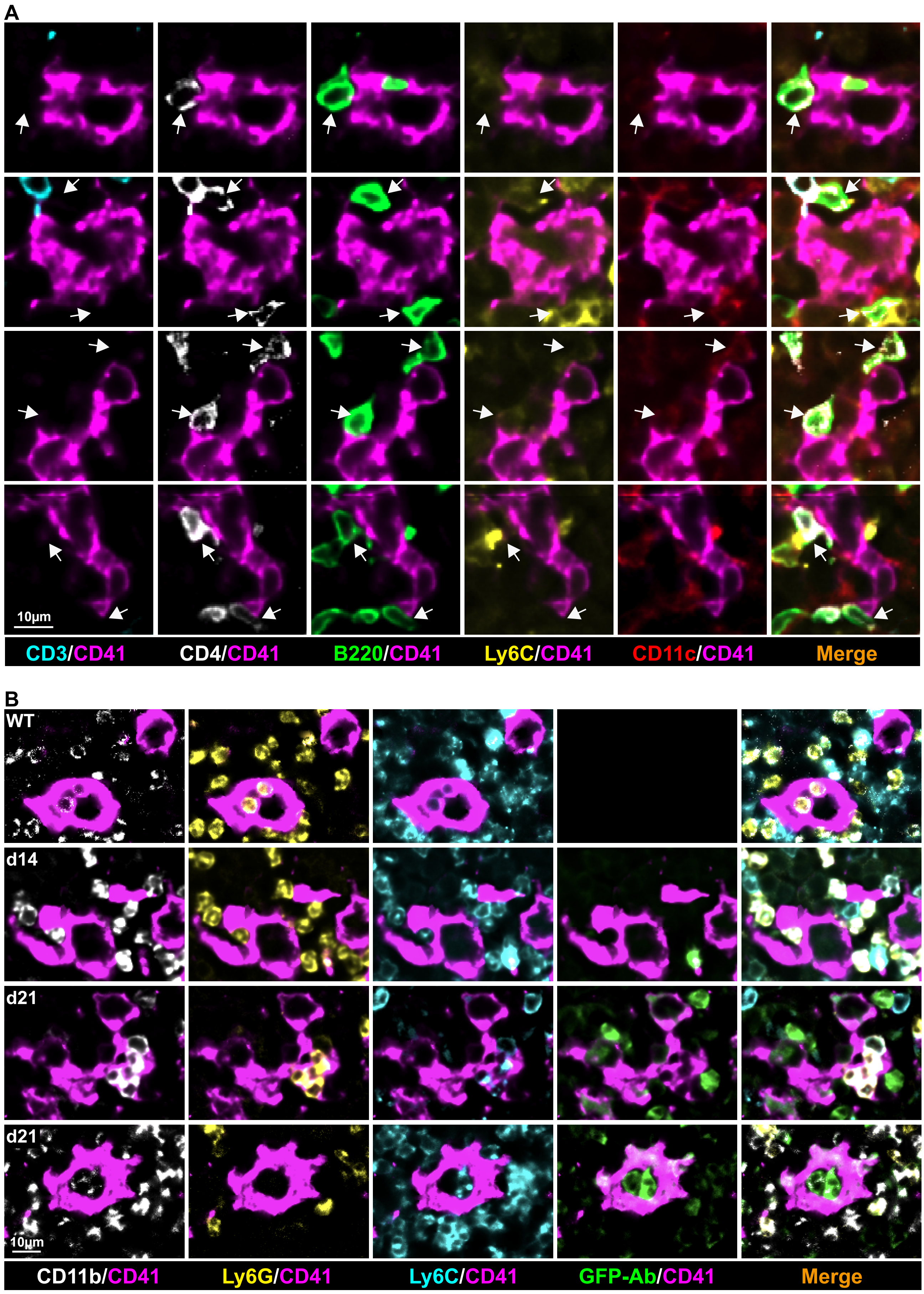
A
10µm
CD3/CD41
CD4/CD41
B220/CD41
Ly6C/CD41
CD11c/CD41
Merge
B
WT
d14
d21
d21
10µm
CD11b/CD41
Ly6G/CD41
Ly6C/CD41
GFP-Ab/CD41
Merge

### Figure S14

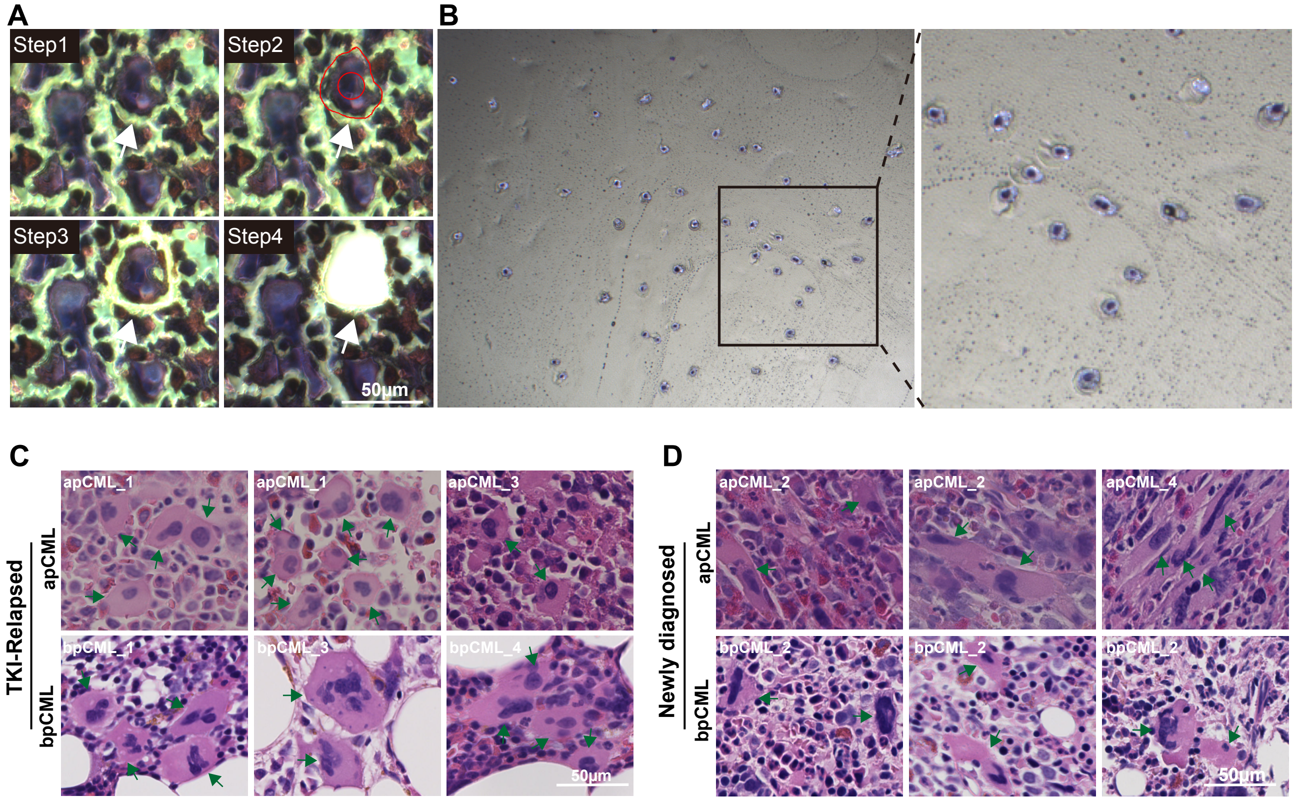
A
Step1
Step2
Step3
Step4
50μm
B
C
TKI-Relapsed
apCML
bpCML
apCML_1
apCML_1
apCML_3
bpCML_1
bpCML_3
bpCML_4
50μm
D
Newly diagnosed
apCML
bpCML
apCML_2
apCML_2
apCML_4
bpCML_2
bpCML_2
bpCML_2
50μm

### Table S1

| Antigen | Clone | Manufacture | Catalog No. | CODEX oligo | Working dilution | Exposure time | Comment |
|---|---|---|---|---|---|---|---|
| B220 | RA3.3/6.1 | BioXcell | BE0067 | 3 | 1:100 | 1/15s | |
| CD105 | MJ7/18 | Biolegend | 120402 | 69 | 1:20 | 1/15s | |
| CD106 | 429 | BD Biosciences | 553330 | 76 | 1:100 | 1/25s | |
| CD117 | 2B8 | Biolegend | 105802 | 30 | 1:200 | 1/50s | |
| CD11b | M1/70 | Biolegend | 101202 | 5 | 1:100 | 1/50s | |
| CD11c | HL3 | BD Biosciences | 553799 | 26 | 1:50 | 1/15s | |
| CD127 | polyclonal | Thermo Fisher Scientific | PA5-47927 | 79 | 1:20 | 1/8.5s | |
| CD135 | 113308 | Novus Biologicals | MAB7681-SP | 61 | 1:50 | 1/6s | |
| CD150 | mShad150 | eBioscience | CUST03314 | 24 | 1:100 | 1/60s | |
| CD115 | 460615 | Novus Biologicals | MAB388 | 29 | 1:100 | 1/30s | not working |
| CD16/32 | 2.4G2 | BD Biosciences | 553142 | 28 | 1:400 | 1/30s | |
| CD169 | polyclonal | Bio-Rad | Not available | 46 | 1:50 | 1/25s | |
| CD21/35 | 7G8 | BD Biosciences | Not available | 72 | 1:50 | 1/15s | |
| CD27 | LG.3A10 | BD Biosciences | 553777 | 32 | 1:50 | 1/20s | |
| CD3 | 17A2 | BD Biosciences | 55273 | 6 | 1:200 | 1/15s | |
| CD31 | MEC13.3 | Biolegend | 102502 | 66 | 1:200 | 1/30s | |
| CD34 | RAM34 | eBioscience | 14-0341-82 | 59 | 1:20 | 1/12s | |
| CD38 | 90 | Biolegend | 102702 | 44 | 1:200 | 1/50s | |
| CD4 | RM4-5 | BD Biosciences | 553043 | 7 | 1:400 | 1/60s | |
| CD41 | MWREg30 | Biolegend | 133902 | 75 | 1:500 | 1/300s | |
| CD45 | 30-F11 | BD Biosciences | 553076 | 62 | 1:200 | 1/70s | |
| CD47 | miap410 | Biolegend | 127501 | 65 | 1:300 | 1/40s | |
| CD48 | HM48-1 | Biolegend | 103402 | 15 | 1:200 | 1/30s | |
| CD49d | R1-2 | Biolegend | 103610 | 68 | 1:50 | 1/70s | |
| CD5 | 53-7.3 | BD Biosciences | 553017 | 77 | 1:50 | 1/15s | |
| CD61 | 2C9.G2 | Biolegend | 104302 | 49 | 1:200 | 1/30s | |
| CD71 | C2F2 | BD Biosciences | 553264 | 21 | 1:400 | 1/175s | |
| CD86 | GL-1 | Biolegend | 105002 | 51 | 1:100 | 1/12s | |
| CD8 | 53-6.7 | BD Biosciences | 553027 | 11 | 1:400 | 1/70s | |
| CD90 | G7 | BD Biosciences | 553016 | 63 | 1:300 | 1/60s | |
| EMCN (Endomucin) | polyclonal | R&D Systems | AF4666 | 80 | 1:100 | 1/15s | |
| ESAM (Endothelial Cell Adhesion Molecule) | 1G8 | Biolegend | 93892 | 58 | 1:100 | 1/30s | |
| F4/80 | T45-2342 | BD Biosciences | 565409 | 48 | 1:200 | 1/50s | |
| GFP | N/A | N/A | N/A | N/A | N/A | 1/6s | |
| GFP-Ab | FM264G | Biolegend | 338002 | 67 | 1:20 | 1/15s | |
| ICOS | C398.4A | Biolegend | 313502 | 60 | 1:20 | 1/8.5s | |
| IgM | II/41 | BD Biosciences | 553435 | 33 | 1:50 | 1/25s | |
| Ki-67 | B56 | BD Biosciences | 556003 | 52 | 1:50 | 1/15s | |
| LepR | polyclonal | R&D Systems | AF497 | 38 | 1:300 | 1/15s | |
| Ly6C | HK1.4 | BioXcell | Not available | 41 | 1:100 | 1/20s | |
| Ly6G | 1A8 | BioXcell | BE0075-1 | 55 | 1:400 | 1/100s | |
| MHC-II | M5/114.15.2 | BioXcell | BE0108 | 36 | 1:800 | 1/300s | |
| NG2 (Neural/glial antigen 2) | polyclonal | Millipore | AB5320 | 23 | 1:50 | 1/15s | |
| NKp46 | 29A1.4 | BD Biosciences | 560754 | 45 | 1:20 | 1/8.5 | |
| OPN (Osteopontin) | polyclonal | R&D Systems | AF808 | 2 | 1:400 | 1/300s | |
| PD-1 | 29F.1A12 | Biolegend | 135202 | 74 | 1:200 | 1/20s | |
| PD-L1 | 10F.9G2 | Biolegend | 124302 | 81 | 1:100 | 1/8.5s | |
| Perilipin | polyclonal | Sigma Aldrich | P1873 | 70 | 1:100 | 1/25s | |
| Podoplanin | 8.1.1 | Biolegend | 127402 | 8 | 1:100 | 1/15s | |
| Runx2 | 232902 | R&D Systems | MAB2006 | 14 | 1:200 | 1/100s | not working |
| Sca-1 | D7 | Biolegend | 108102 | 56 | 1:100 | 1/12s | |
| Siglec H | 551 | Biolegend | 129602 | 20 | 1:100 | 1/6s | not working |
| Sirpα | P84 | Biolegend | 144001 | 42 | 1:50 | 1/15s | |
| TCRgd | UC7-13D5 | Biolegend | 107502 | 71 | 1:50 | 1/30s | not working |
| TER-119 | TER-119 | BD Biosciences | 553670 | 43 | 1:400 | 1/120s | |
| Tie2 | TEK4 | Biolegend | 124002 | 57 | 1:50 | 1/30s | |

**Table S2**

| Categories | Cell Type | Antigen expression pattern |
|---|---|---|
| **Myeloid cells** | Common monocyte progenitors (CoMPs) | lin$^{-}$CD11b-Ly6C+Ly6G- |
| | Early neutrophil progenitors (eNePs) | lin$^{-}$CD117+CD71+CD49d+CD38- |
| | Granulocytes | CD11b+Ly6C+Ly6G+ |
| | MHC-II$^{-}$ macrophages (MHC-II$^{-}$ macs) | F4/80+MHC-II- |
| | MHC-II$^{+}$ macrophages (MHC-II$^{+}$ macs) | F4/80+MHC-II+ |
| | Monocytes | CD11b+Ly6C+Ly6G- |
| | CD8$^{+}$ T cells | PD-L1+CD11c+MHC-II+ |
| | plasmacytoid dendritic cells (pDCs) | CD3-CD4+B220+Ly6C$^{lo}$CD11c$^{lo}$ |
| **Erythroblasts** | EryAs | CD71+TER119+ and larger cell size, smaller population |
| | EryBs | CD71+TER119+ and smaller cell size, larger population |
| | EryCs | CD71-TER119+ |
| | ProEs | CD71+TER119- |
| **B cells/plasma cells** | B cells | B220+IgM- |
| | Plasma cells | B220+IgM+ |
| **Unidentified** | Ki-67$^{+}$ unidentified | Ki67+ and negative for all the lineage markers |
| | Unidentified | Negative for all the lineage markers |
| **Vessels** | Arterioles | CD31+Sca-1+ |
| | LepR$^{+}$ pericytes | LepR+CD31- |
| | Sinuses | CD31+Sca-1- |
| **Leukemic cells** | Leukemic cells | GFP+GFP-Ab+ |
| | Leukemic stem cells (LSCs) | lin$^{-}$CD117+/CD34+GFP+GFP-Ab+ |
| | PD-L1$^{+}$ leukemic cells | PD-L1+GFP+GFP-Ab+ |
| **Antigen presenting cells(APCs)** | APCs | CD16/CD32+MHCII+ |
| | PD-L1$^{+}$ APCs | PD-L1+CD16/CD32+MHCII+ |
| **Mixed cell types** | CD90$^{+}$ leukocytes | CD90+CD45+ and negative for all the lineage markers |
| | Erythroblastic islands (EBIs) | Macrophage surrounded by erythroblasts |
| | Leukocytes | CD45+ and negative for all the linegae markers |
| | Natural killer cells (NK cells) | CD3-NKp46+ |
| | PD-L1$^{+}$ leukocytes | PD-L1+CD45+ and negative for all the lineage markers |
| | Podo$^{+}$ stromal cells | Podoplanin+ |
| **T cells** | CD4$^{+}$ T cells | CD3+CD4+ |
| | CD8$^{+}$ T cells | CD3+CD8+ |
| | Double negative T cells (DN T cells) | CD3+CD4-CD8- |
| | PD-1$^{+}$CD4$^{+}$ T cells | PD-1+CD3+CD4+ |
| | PD-1$^{+}$CD8$^{+}$ T cells | PD-1+CD3+CD8+ |
| **Bone cells** | Bone cells | CD61+ and/or osteopontin+ |
| **hematopoietic stem/progenitor cells (HSPCs)** | CD34$^{-}$CD117$^{+}$ HSPCs | lin$^{-}$CD34-CD117+ |
| | CD34$^{+}$CD117$^{-}$ HSPCs | lin$^{-}$CD34+CD117 |
| | CD34$^{+}$CD117$^{+}$ HSPCs | lin$^{-}$CD34+CD117+ |
| **Megakaryocytes** | Megakaryocytes | CD41+ (manually identified) |
| **Adipocytes** | Adipocytes | Perilipin+ (manually identified) |

### Table S3

#### CML Patient Metadata

| Sample ID | Gender | Age (year) | Diagnosis_histology | Diagnosis_clinical | Newly_Diagnosed/Relapsed | Prior_TKIs_treatment | mutations_(VAF) | BCR_ABL_mut_type | Blasts_BM | Blasts_PB |
|---|---|---|---|---|---|---|---|---|---|---|
| apCML1 | male | 72 | CML accelerated phase | CML, BCR ABL pos. | relapsed | Yes | BCR-ABL (47%) | b2a2 | 15% | NA |
| apCML2 | male | 27 | CML accelerated phase | CML, BCR ABL pos. | newly diagnosed | none | BCR-ABL (69.2%), NF1 | NA | 2.5% | 5% |
| apCML3 | male | 63 | CML accelerated phase | CML, BCR ABL pos. | relapsed | Yes | BCR-ABL (42.3%) | b2a2 | <5% | 6% |
| apCML4 | female | 60 | CML accelerated phase | CML, BCR ABL pos. | newly diagnosed | none | NA | NA | 5%-10% | 10%-15% |
| bpCML1 | male | 53 | CML blast phase | CML, BCR ABL pos. | relapsed | Yes | BCR-ABL (36.2%) | e14e13a2 | 40% | 22% |
| bpCML2 | male | 55 | CML blast phase | CML, BCR ABL pos. | newly diagnosed | none | BCR-ABL (51.9%) | NA | 50-60% | 45% |
| bpCML3 | male | 62 | CML blast phase | CML, BCR ABL pos. | relapsed | Yes | BCR-ABL (85.3%) | e14a2 | 90% | 16% |
| bpCML5 | male | 47 | CML blast phase | CML, BCR ABL pos. | relapsed | Yes | BCR-ABL (19.54%) | b2a2/b3a2 | 39% | 44% |
| bpCML7 | female | 62 | CML blast phase | CML, BCR ABL pos. | relapsed | Yes | NA | NA | >30% | NA |
| bpCML8 | female | 57 | CML blast phase | CML, BCR ABL pos. | relapsed | Yes | BCR-ABL (12%) | NA | 39% | 25% |
| bpCML9 | female | 62 | CML blast phase | CML, BCR ABL pos. | relapsed | Yes | NA | NA | >80% | NA |
| bpCML10 | male | 57 | CML blast phase | CML, BCR ABL pos. | relapsed | Yes | BCR-ABL (103%) | b2a2 | 95% | NA |
| CML1 | male | 30 | CML chronic phase | CML, BCR ABL pos. | newly diagnosed | none | BCR-ABL (36%) | e13a2 (b2a2) | <5% | 6% |
| CML2 | female | 50 | CML chronic phase | CML, BCR ABL pos. | newly diagnosed | none | BCR-ABL | NA | 2% | 0 |
| CML3 | female | 55 | CML chronic phase | CML, BCR ABL pos. | newly diagnosed | none | BCR-ABL (41%) | e14a2 (b3a2) | 2% | 1% |
| CML4 | male | 66 | CML chronic phase | CML, BCR ABL pos. | newly diagnosed | none | BCR-ABL (53%) | b2a2, b3a2 | <5% | 9% |
| CML5 | female | 58 | CML chronic phase | CML, BCR ABL pos. | newly diagnosed | none | BCR-ABL (28%) | b3a2 | <5% | 2% |
| CML6 | female | 67 | CML chronic phase | CML, BCR ABL pos. | newly diagnosed | none | BCR-ABL | b3a2, e1a2 | 2% | 1% |
| CML7 | female | 57 | CML chronic phase | CML, BCR ABL pos. | newly diagnosed | none | BCR-ABL (33%) | b2a2, b3a2 | 1.5% | 3% |
| CML8 | female | 57 | CML chronic phase | CML, BCR ABL pos. | newly diagnosed | none | BCR-ABL (33%) | b3a2 | 2.5% | 3% |
| CML9 | male | 31 | CML chronic phase | CML, BCR ABL pos. | newly diagnosed | none | BCR-ABL (260%) | b2a2 | <5% | 12% |
| CML10 | male | 77 | CML chronic phase | CML, BCR ABL pos. | newly diagnosed | none | BCR-ABL (50%) | b2a2, b3a2 | 7.5% | 0 |
| CML11 | male | 27 | CML chronic phase | CML, BCR ABL pos. | newly diagnosed | none | BCR-ABL | b3a2 | <5% | 2% |
| CML12 | male | 73 | CML chronic phase | CML, BCR ABL pos. | newly diagnosed | none | BCR-ABL (28%) | b3a2 | 2.5% | 0 |
| CML13 | female | 48 | CML chronic phase | CML, BCR ABL pos. | newly diagnosed | none | BCR-ABL (38%) | b2a2/b3a2 | <5% | 0 |
| CML14 | male | 37 | CML chronic phase | CML, BCR ABL pos. | newly diagnosed | none | BCR-ABL (56%) | e14a2 (b3a2) | 1.5% | 0 |
| CML15 | male | 56 | CML chronic phase | CML, BCR ABL pos. | newly diagnosed | none | BCR-ABL | b2a2 | <5% | 1% |
| CML16 | male | 32 | CML chronic phase | CML, BCR ABL pos. | newly diagnosed | none | BCR-ABL (76%) | e13a2 (b2a2) | <5% | 1% |

#### "Healthy" Bone Marrow

| Sample ID | Gender | Age (year) | Prior Chemotherapy | Diagnosis_PAS | Results |
|---|---|---|---|---|---|
| HBM_1 | female | 59 | no | AIHA | no significant abnormalities detected |
| HBM_2 | male | 42 | no | Suspicion of paraaortitis in M. Ormond | no significant abnormalities detected |
| HBM_3 | male | 61 | no | ED FL | no signs of lymphoma inflitration |
| HBM_4 | female | 40 | no | Suspicion of Lymphoma | no signs of lymphoma inflitration |
| HBM_5 | male | 59 | no | ED Vitreoretinal lymphoma | no signs of lymphoma inflitration |
| HBM_6 | female | 56 | no | Suspicion of Myeloma or Lymphoma | no signs of lymphoma or myeloma inflitration |
| HBM_7 | male | 50 | no | MALT lymphoma of the stomach | no signs of lymphoma inflitration |
| HBM_8 | female | 74 | no | MALT lymphoma of the parotid gland | no signs of lymphoma inflitration |
| HBM_9 | female | 38 | no | ED PMBL | no signs of lymphoma inflitration |
| HBM_10 | male | 65 | no | ED FL | no signs of lymphoma inflitration |
| HBM_11 | male | 48 | no | Suspicion of MDS | no significant maturation abnormalities or increase in blast cells |
| HBM_12 | male | 21 | no | Suspicion of Burkitt lymphoma | no signs of lymphoma inflitration |
| HBM_13 | male | 41 | no | NLPHL | no signs of lymphoma inflitration |
| HBM_14 | female | 81 | no | ED nodal MZL | no signs of lymphoma inflitration |
| HBM_15 | female | 59 | no | MZL Parotid Gland | no signs of lymphoma inflitration |
| HBM_16 | male | 59 | no | DLBCL | no signs of lymphoma inflitration |
| HBM_17 | female | 76 | no | DLBCL | no signs of lymphoma inflitration |

AIHA, autoimmune hemolytic anemia; ED, early-stage disease; FL, follicular lymphoma; MALT, mucosa-associated lymphoid tissue; PMBL, primary mediastinal (thymic) large B-cell lymphoma;
PMBL, primary mediastinal (thymic) large B-cell lymphoma; MDS, myelodysplastic syndromes; NLPHL, nodular lymphocyte-predominant hodgkin lymphoma;
MZL, marginal zone lymphoma; DLBCL, diffuse large B-cell lymphoma.

### Table S4

| Oligo No. | Antibody oligo sequence (5'-3') | Fluorescent oligonucleotide sequences (5'-3') | Fluorescent oligonucleotide sequences (5'-3') | Blocking component 4 (BC4) oligos (5'-3') |
|---|---|---|---|---|
| 2 | /mal/ATGGTTTAGGACTAC | /5ATTO550N/GTAGTCCTAAACCAT | | GTAGTCCTAAACCAT |
| 3 | /mal/TACTCCTCGCCG | | /5Alex647N/CGGCGAGGAGTA | CGGCGAGGAGTA |
| 5 | /mal/TCTCCCATTAGTCGG | /5ATTO550N/CCGACTAATGGGAGA | | CCGACTAATGGGAGA |
| 6 | /mal/TGGATGTGTTACGAT | | /5Alex647N/ATCGTAACACATCCA | ATCGTAACACATCCA |
| 7 | /mal/CGCTAAGATATTCTAAG | /5ATTO550N/CTTAGAATATCTTAGCG | | CTTAGAATATCTTAGCG |
| 8 | /mal/CGCAGATGAATATTC | /5ATTO550N/GAATATTCATCTGCG | | GAATATTCATCTGCG |
| 11 | /mal/GGGTTATACACTCGT | /5ATTO550N/ACGAGTGTATAACCC | | ACGAGTGTATAACCC |
| 14 | /mal/AGATTCAGTCTCTCG | /5ATTO550N/CGAGAGACTGAATCT | | CGAGAGACTGAATCT |
| 15 | /mal/CTGTAATAGGCACTA | /5ATTO550N/TAGTGCCTATTACAG | | TAGTGCCTATTACAG |
| 20 | /mal/ATGTAGGATGGTCTC | /5ATTO550N/GAGACCATCCTACAT | | GAGACCATCCTACAT |
| 21 | /mal/CGTGCCGTTTCAC | | /5Alex647N/GTGAAACGGCACG | GTGAAACGGCACG |
| 23 | /mal/GGTTTCCTCAGACAC | | /5Alex647N/GTGTCTGAGGAAACC | GTGTCTGAGGAAACC |
| 24 | /mal/ATAAGGGCTCATTGT | /5ATTO550N/ACAATGAGCCCTTAT | | ACAATGAGCCCTTAT |
| 26 | /mal/CACTTGTCTAACCAA | | /5Alex647N/TTGGTTAGACAAGTG | TTGGTTAGACAAGTG |
| 28 | /mal/TGGTCCACTAACGTA | | /5Alex647N/TACGTTAGTGGACCA | TACGTTAGTGGACCA |
| 29 | /mal/ATAGGGCATTTGAAG | | /5Alex647N/CTTCAAATGCCCTAT | CTTCAAATGCCCTAT |
| 30 | /mal/CACATGAGCGAATCA | /5ATTO550N/TGATTCGCTCATGTG | | TGATTCGCTCATGTG |
| 32 | /mal/TACCAAATCCTGATG | | /5Alex647N/CATCAGGATTTGGTA | CATCAGGATTTGGTA |
| 33 | /mal/TTATCATGAGGAGCG | | /5Alex647N/CGCTCCTCATGATAA | CGCTCCTCATGATAA |
| 36 | /mal/ACCTACACAATGCTA | /5ATTO550N/TAGCATTGTGTAGGT | | TAGCATTGTGTAGGT |
| 38 | /mal/GCGTCTACTTATAAG | /5ATTO550N/CTTATAAGTAGACGC | | CTTATAAGTAGACGC |
| 41 | /mal/TGTATGAGTAGTAATCT | | /5Alex647N/AGATTACTACTCATACA | AGATTACTACTCATACA |
| 42 | /mal/TCTAAGTCAGAGAGC | | /5Alex647N/GCTCTCTGACTTAGA | GCTCTCTGACTTAGA |
| 43 | /mal/GACATTATCCGTGAT | /5ATTO550N/ATCACGGATAATGTC | | ATCACGGATAATGTC |
| 44 | /mal/TCACTACTATTAGTACT | /5ATTO550N/AGTACTAATAGTAGTGA | | AGTACTAATAGTAGTGA |
| 45 | /mal/GCCAGAATGCCA | | /5Alex647N/TGGCATTCTGGC | TGGCATTCTGGC |
| 46 | /mal/GACTCGAACCTGAG | | /5Alex647N/CTCAGGTTCGAGTC | CTCAGGTTCGAGTC |
| 48 | /mal/GCACGGCAAAGTG | /5ATTO550N/CACTTTGCCGTGC | | CACTTTGCCGTGC |
| 49 | /mal/ATAACGCCTCGTATC | /5ATTO550N/GATACGAGGCGTTAT | | GATACGAGGCGTTAT |
| 51 | /mal/CGTGCGGGAAAAT | | /5Alex647N/ATTTTCCCGCACG | ATTTTCCCGCACG |
| 52 | /mal/ACCAGGTGATTGCAT | | /5Alex647N/ATGCAATCACCTGGT | ATGCAATCACCTGGT |
| 55 | /mal/AGGTCAACTCGCAC | /5ATTO550N/GTGCGAGTTGACCT | | GTGCGAGTTGACCT |
| 56 | /mal/GGTCACATGGTCGTT | | /5Alex647N/AACGACCATGTGACC | AACGACCATGTGACC |
| 57 | /mal/GCGGATTTCGTATTT | /5ATTO550N/AAATACGAAATCCGC | | AAATACGAAATCCGC |
| 58 | /mal/CGTCAGTACTTTCAG | /5ATTO550N/CTGAAAGTACTGACG | | CTGAAAGTACTGACG |
| 59 | /mal/GCTTATTATGGACTTC | | /5Alex647N/GAAGTCCATAATAAGC | GAAGTCCATAATAAGC |
| 60 | /mal/ACAAAACTGCTGTCG | | /5Alex647N/CGACAGCAGTTTTGT | CGACAGCAGTTTTGT |
| 61 | /mal/TCTTATTCCCGAATA | | /5Alex647N/TATTCGGGAATAAGA | TATTCGGGAATAAGA |
| 62 | /mal/TAGGGGAACAGGTTG | /5ATTO550N/CAACCTGTTCCCCTA | | CAACCTGTTCCCCTA |
| 63 | /mal/AATTAGCTTAAGAGAGT | /5ATTO550N/ACTCTCTTAAGCTAATT | | ACTCTCTTAAGCTAATT |
| 65 | /mal/GATAAATATTTTACAGAGT | /5ATTO550N/ACTCTGTAAAATATTTATC | | ACTCTGTAAAATATTTATC |
| 66 | /mal/TACGTGCTTTGGTT | /5ATTO550N/AACCAAAGCACGTA | | AACCAAAGCACGTA |
| 67 | /mal/GACGACGAAGGC | | /5Alex647N/GCCTTCGTCGTC | GCCTTCGTCGTC |
| 68 | /mal/CTTCTTGTGGAACC | /5ATTO550N/GGTTCCACAAGAAG | | GGTTCCACAAGAAG |
| 69 | /mal/CCCGGCAGTT | | /5Alex647N/AACTGCCGGG | AACTGCCGGG |
| 70 | /mal/AACCAAACTGACCG | | /5Alex647N/CGGTCAGTTTGGTT | CGGTCAGTTTGGTT |
| 71 | /mal/TCACCCCCAGC | /5ATTO550N/GCTGGGGGTGA | | GCTGGGGGTGA |
| 72 | /mal/AACGCGACGGAT | | /5Alex647N/ATCCGTCGCGTT | ATCCGTCGCGTT |
| 74 | /mal/ATTTGCTTCGACGA | /5ATTO550N/TCGTCGAAGCAAAT | | TCGTCGAAGCAAAT |
| 75 | /mal/CGCTTGGGTGTTTA | /5ATTO550N/TAAACACCCAAGCG | | TAAACACCCAAGCG |
| 76 | /mal/CTGTCGGTCGTCA | | /5Alex647N/TGACGACCGACAG | TGACGACCGACAG |
| 77 | /mal/ATTTCAACAAATATTGTT | | /5Alex647N/AACAATATTTGTTGAAAT | AACAATATTTGTTGAAAT |
| 79 | /mal/GCCGGAAGTGGT | | /5Alex647N/ACCACTTCCGGC | ACCACTTCCGGC |
| 80 | /mal/CGGGGCACCA | | /5Alex647N/TGGTGCCCCG | TGGTGCCCCG |
| 81 | /mal/CAAGGAACTACCGA | | /5Alex647N/TCGGTAGTTCCTTG | TCGGTAGTTCCTTG |

#### Table S5

##### Abs Mix 1: 4 °C o/n

| Antigen | Oligo | Clone | Dilution |
|---|---|---|---|
| CD105 | 69 | MJ7/18 | 1:20 |
| CD127 | 79 | polyclonal | 1:20 |
| CD150 | 24 | mShad150 | 1:100 |
| CD169 | 46 | polyclonal | 1:50 |
| CD27 | 32 | LG.3A10 | 1:50 |
| CD34 | 59 | RAM34 | 1:20 |
| CD38 | 44 | 90 | 1:200 |
| CD45 | 62 | 30-F11 | 1:200 |
| CD48 | 15 | HM48-1 | 1:200 |
| CD61 | 49 | 2C9.G2 | 1:200 |
| CD90 | 63 | G7 | 1:300 |
| ESAM | 58 | 1G8 | 1:100 |
| GFP-Ab | 67 | FM264 | 1:20 |
| LepR | 38 | polyclonal | 1:300 |
| Ly6C | 41 | HK1.4 | 1:100 |
| MHCII | 36 | M5/114.15.2 | 1:800 |
| NG2 | 23 | polyclonal | 1:50 |
| OPN | 2 | polyclonal | 1:400 |
| PD-L1 | 81 | 10F.9G2 | 1:100 |
| Perilipin | 70 | polyclonal | 1:100 |
| Runx2 | 14 | 232902 | 1:200 |
| Sca-1 | 56 | D7 | 1:100 |
| Sirpα | 42 | P84 | 1:50 |
| TCRgd | 71 | UC7-13D5 | 1:50 |
| Tie2 | 57 | TEK4 | 1:50 |

##### Abs Mix 2: 3h RT

| Antigen | Oligo | Clone | Dilution |
|---|---|---|---|
| B220 | 3 | RA3.3/6.1 | 1:100 |
| CD106 | 76 | 429 | 1:100 |
| CD117 | 30 | 2B8 | 1:200 |
| CD11b | 5 | M1/70 | 1:100 |
| CD11c | 26 | HL3 | 1:50 |
| CD135 | 61 | 113308 | 1:50 |
| CD16/32 | 28 | 2.4G2 | 1:400 |
| CD21/35 | 72 | 7G8 | 1:50 |
| CD3 | 6 | 17A2 | 1:200 |
| CD31 | 66 | MEC13.3 | 1:200 |
| CD4 | 7 | RM4-5 | 1:400 |
| CD41 | 75 | MWREg30 | 1:500 |
| CD47 | 65 | miap410 | 1:300 |
| CD49d | 68 | R1-2 | 1:50 |
| CD5 | 77 | 53-7.3 | 1:50 |
| CD71 | 21 | C2F2 | 1:400 |
| CD86 | 51 | GL-1 | 1:100 |
| CD8 | 11 | 53-6.7 | 1:400 |
| Endomucin | 80 | polyclonal | 1:100 |
| F4/80 | 48 | T45-2342 | 1:200 |
| ICOS | 60 | C398.4A | 1:20 |
| IgM | 33 | II/41 | 1:50 |
| Ki-67 | 52 | B56 | 1:50 |
| Ly6G | 55 | 1A8 | 1:400 |
| NKp46 | 45 | 29A1.4 | 1:20 |
| PD-1 | 74 | 29F.1A12 | 1:200 |
| Podoplanin | 29 | 8.1.1 | 1:100 |
| Siglec H | 20 | 551 | 1:100 |
| TER-119 | 43 | TER-119 | 1:400 |

### Table S6

| Cycle | Exposure time (s) | Ab | Oligo | Clone | Dilution | Exposure time (s) | Ab | Oligo | Clone | Dilution | Exposure time (s) |
|---|---|---|---|---|---|---|---|---|---|---|---|
| 1 | 1/40 | Runx2 | 14 | 232902 | 1:200 | 1/100 | Ki-67 | 52 | B56 | 1:50 | 1/15 |
| 2 | 1/20 | CD150 | 24 | mShad150 | 1:100 | 1/60 | GFP-Ab | 67 | FM264 | 1:20 | 1/15 |
| 3 | 1/10 | ESAM | 58 | 1G8 | 1:100 | 1/30 | CD127 | 79 | polyclonal | 1:20 | 1/8.5 |
| 4 | 1/600 | Tie2 | 57 | TEK4 | 1:50 | 1/30 | ICOS | 60 | C398.4A | 1:20 | 1/8.5 |
| 5 | 1/600 | TCRgd | 71 | UC7-13D5 | 1:50 | 1/30 | CD34 | 59 | RAM34 | 1:20 | 1/12 |
| 6 | 1/600 | CD11b | 5 | M1/70 | 1:100 | 1/50 | CD135 | 61 | 113308 | 1:50 | 1/6 |
| 7 | 1/600 | CD49d | 68 | R1-2 | 1:50 | 1/70 | CD105 | 69 | MJ7/18 | 1:20 | 1/15 |
| 8 | 1/600 | Podo | 8 | 8.1.1 | 1:100 | 1/15 | NKp46 | 45 | 29A1.4 | 1:20 | 1/8.5 |
| 9 | 1/600 | CD31 | 66 | MEC13.3 | 1:200 | 1/30 | CD5 | 77 | 53-7.3 | 1:50 | 1/15 |
| 10 | 1/600 | CD38 | 44 | 90 | 1:200 | 1/50 | CD27 | 32 | LG.3A10 | 1:50 | 1/20 |
| 11 | 1/600 | LepR | 38 | polyclonal | 1:300 | 1/15 | CD11c | 26 | HL3 | 1:50 | 1/15 |
| 12 | 1/600 | PD-1 | 74 | 29F.1A12 | 1:200 | 1/20 | Sca-1 | 56 | D7 | 1:100 | 1/12 |
| 13 | 1/600 | CD47 | 65 | miap410 | 1:300 | 1/40 | CD21/35 | 72 | 7G8 | 1:50 | 1/15 |
| 14 | 1/600 | TER-119 | 43 | TER-119 | 1:400 | 1/120 | NG2 | 23 | polyclonal | 1:50 | 1/15 |
| 15 | 1/600 | F4/80 | 48 | T45-2342 | 1:200 | 1/50 | Perilipin | 70 | polyclonal | 1:100 | 1/25 |
| 16 | 1/600 | CD117 | 30 | 2B8 | 1:200 | 1/50 | Sirpα | 42 | P84 | 1:50 | 1/15 |
| 17 | 1/600 | CD48 | 15 | HM48-1 | 1:200 | 1/30 | IgM | 33 | II/41 | 1:50 | 1/25 |
| 18 | 1/600 | CD90 | 63 | G7 | 1:300 | 1/60 | CD3 | 6 | 17A2 | 1:200 | 1/15 |
| 19 | 1/600 | CD45 | 62 | 30-F11 | 1:200 | 1/70 | CD169 | 46 | polyclonal | 1:50 | 1/25 |
| 20 | 1/600 | CD61 | 49 | 2C9.G2 | 1:200 | 1/30 | CD106 | 76 | 429 | 1:100 | 1/25 |
| 21 | 1/600 | Ly6G | 55 | 1A8 | 1:400 | 1/100 | B220 | 3 | Ra3-6B2 | 1:100 | 1/15 |
| 22 | 1/600 | SigH | 20 | 551 | 1:100 | 1/8.5 | Emcn | 80 | polyclonal | 1:100 | 1/15 |
| 23 | 1/600 | CD4 | 7 | RM4-5 | 1:400 | 1/60 | PD-L1 | 81 | 10F.9G2 | 1:100 | 1/8.5 |
| 24 | 1/600 | CD8 | 11 | 53-6.7 | 1:400 | 1/70 | CD115 | 29 | 460615 | 1:100 | 1/40 |
| 25 | 1/600 | CD41 | 75 | MWREg30 | 1:500 | 1/300 | CD86 | 51 | GL-1 | 1:100 | 1/12 |
| 26 | 1/600 | OPN | 2 | polyclonal | 1:400 | 1/300 | Ly6C | 41 | HK1.4 | 1:100 | 1/20 |
| 27 | 1/600 | MHCII | 36 | M5/114.15.2 | 1:800 | 1/300 | CD16/32 | 28 | 2.4G2 | 1:400 | 1/30 |
| 28 | 1/600 | blank | | | | 1/8.5 | CD71 | 21 | C2F2 | 1:400 | 1/175 |
| 29 | 1/600 | blank | | | | 1/30 | blank | | | | 1/8.5 |
| 30 | 1/600 | blank | | | | 1/100 | blank | | | | 1/40 |
| 31 | 1/600 | blank | | | | 1/300 | DRAQ5 | | | 1:100 | 1/25 |

### Table S7

| **Lysis Buffer** | **Manufacturer & Catalog No.** | **Volume/gram** |
|---|---|---|
| Trimethylglycine (TMG), 5 M | MilliporeSigma, #B0300 | 100 µL |
| Molecular biology-grade water | Thermo Fisher Scientific, #AM9932 | 10.79 µL |
| Tris base solution, 1.5 M | MilliporeSigma, #T1699 | 16.67 µL |
| Hydrochloric acid solution (HCl), 1 M | MilliporeSigma, #H9892 | 6.9 µL |
| Potassium chloride solution (KCl) | Thermo Fisher Scientific, #J62422 | 25 µL |
| Disodium EDTA ($Na_2EDTA$), 500 mM | Thermo Fisher Scientific, #J15694 | 0.14 µL |
| Polysorbate 20 solution (Tween 20), 10% v/v | MilliporeSigma, #P9416 | 0.5 µL |
| **DNA Storage Buffer** | | |
| Tris, 1 M, PH 8.0 | MilliporeSigma, #T1699 | 1 mL |
| Disodium EDTA ($Na_2EDTA$), 500 mM | Thermo Fisher Scientific, #J15694 | 200 µL |
| Sodium azide ($NaN_3$) | Sigma-Aldrich, #S8032 | 0.02 g |
| Polysorbate 20 solution (Tween 20), 10% v/v | MilliporeSigma, #P9416 | 50 µL |
| Molecular biology-grade water | Thermo Fisher Scientific, #AM9932 | 98.75 mL |
| **Reverse transcription (RT) Mix** | | |
| DNA storage buffer | as above | 66.5 µL |
| Kapa HiFi HotStart plus dNTPs (Kapa dNTPs mix), 10 mM | Roche, #07958889001 | 30 µL |
| RT primer, GTGACTGGAGTTCAGACGTGTGCTCTTCCGATCTTTTTTTTTTGTTTTTTTTTGTTTTTTTTTV, 50µM | biomers | 3.5 µL |
| **Lysis Mix** | | |
| Lysis Buffer | as above | 160 µL |
| RT Mix | as above | 100 µL |
| Proteinase K, 800 U/mL | NEB, #P8107 | 40 µL |
| **Template-switching reverse-transcription (TSRT) Mix** | | |
| Dithiothreitol solution (DTT), 50 mM | MilliporeSigma, #43816 | 100 µL |
| Magnesium chloride ($MgCl_2$), 200 mM | Thermo Fisher Scientific, #AM9530G | 20 µL |
| Proteinase K inhibitor, 5 mM | MilliporeSigma, #539470 | 5 µL |
| RNase inhibitor, 20x | Thermo Fisher Scientific, #AM2696 | 25 µL |
| Template-switch oligo (TSO), CdUACACGACGCdUCdUdUCCGAdUCdUNNNNNrGrGrG | biomers | 25 µL |
| RevertAid H Minus reverse transcriptase | Thermo Fisher Scientific, #EP0451 | 25 µL |
| **PCR Buffer** | | |
| Molecular biology-grade water | Thermo Fisher Scientific, #AM9932 | 169 µL |
| Kapa HiFi HotStart plus dNTPs (Kapa Fidelity Buffer), 5x | Roche, #07958889001 | 100 µL |
| Trimethylglycine (TMG), 5 M | MilliporeSigma, #B0300 | 100 µL |
| Disodium EDTA ($Na_2EDTA$), 500 mM | Thermo Fisher Scientific, #J15694 | 6 µL |
| **PCR Mix** | | |
| PCR Buffer | as above | 375 µL |
| Kapa HiFi HotStart plus dNTPs (Polymerase), 1 U/µL | Roche, #07958889001 | 20 µL |
| Uracil-DNA glycosylase (UNG, UDG), 5 U/µL | NE BioLab, #M0280 | 5 µL |

Note: All solutions prepared for 100 reactions
